# MMV687794 blocks *Plasmodium falciparum* invasion of red blood cells by targeting a Surface-associated Lipid-Interacting Rhoptry Protein, *Pf*SLIRP

**DOI:** 10.64898/2026.02.24.707847

**Authors:** Dawson B. Ling, Ghizal Siddiqui, Christopher A. MacRaild, Yoshiki Yamaryo-Botté, Madeline G. Dans, Betty Kouskousis, Greta E. Weiss, Zahra Razook, Somya Mehra, William Nguyen, Monica Chen, Alyssa E. Barry, Brad E. Sleebs, Tania F. de Koning-Ward, Cyrille Y. Botté, Darren J. Creek, Alan F. Cowman, Niall D. Geoghegan, Kelly L. Rogers, Brendan S. Crabb, Hayley E. Bullen, Paul R. Gilson

## Abstract

Invasion of red blood cells (RBCs) by the human malaria parasite, *Plasmodium falciparum*, drives disease. During invasion, the parasite pushes its way into the RBC while wrapping the RBC membrane around itself to establish the parasitophorous vacuole, a stable niche within the RBC where the parasite grows. To better understand invasion, we investigated the mechanism of action of an invasion-inhibitory compound, MMV687794 (MMV794). Lattice light-sheet microscopy revealed that MMV794 blocks parasite entry by preventing parasitophorous vacuole formation. *In vitro* drug resistance selection of parasites with MMV794 found mutations to the α/β hydrolase, PF3D7_0403800, and engineering one of these mutations (C36W) into parasites by CRISPR/Cas9 recapitulated the resistance phenotype. Expansion microscopy demonstrated that this protein is expressed in schizonts, localising to the surface of rhoptries, which are specialised apical secretory organelles that function during invasion. Lipidomics and proteomics analyses of C36W parasites uncovered widespread changes to lipid composition/homeostasis and altered abundance of proteins involved in invasion, indicating a role for PF3D7_0403800 in invasion-associated lipid metabolism. Finally, we used solvent-induced proteome profiling and reactivity assays to confirm drug-target engagement. Together, our findings identify a novel <u>S</u>urface-associated <u>L</u>ipid-Interacting <u>R</u>hoptry <u>P</u>rotein (*Pf*SLIRP) that coordinates lipid metabolism at the rhoptries to enable RBC invasion.

## Introductions

In 2024, there were an estimated 282 million cases of malaria worldwide, resulting in 610,000 deaths, an overall increase since 2019, with *Plasmodium falciparum* (the most lethal species) accounting for 96% of cases^1^. This rise in cases is, in part, due to the development of parasite resistance to all current antimalarials^2^, including partial resistance to the gold-standard artemisinin combination therapy^3–12^. This presents a major hurdle for malaria eradication and emphasises the need for new antimalarials, particularly with novel mechanisms of action to avoid cross-resistance with existing antimalarials^13^.

Merozoite invasion of red blood cells (RBCs) occurs via a series of tightly coordinated events ^14^ and presents a novel target as it is essential to the proliferation of the disease-causing blood-stage parasites, yet is not currently targeted by clinical antimalarials. The obligate intracellular parasite must quickly internalise within a RBC to survive as the extracellular merozoite form rapidly loses its invasive capacity^15,16^. Blocking invasion would therefore reduce the parasite burden and, ultimately, malaria symptoms^17^. To this end, we previously screened the Medicines for Malaria Venture (MMV) Pathogen Box, a library of 400 compounds, for inhibitors of blood-stage *P. falciparum* invasion^18^. Of the hit compounds, MMV687794 (MMV794) appeared to be invasion-specific and was selected for further study^18^. Using *in vitro* drug resistance selection and genomic analysis, we identified two mutations in MMV794-resistant parasites within the putative α/β hydrolase PF3D7_0403800 (Δ5’UTR and C36W), designating it as the putative target of resistance to MMV794.

PF3D7_0403800 is a serine hydrolase ^19^ belonging to the α/β hydrolase superfamily because it has been predicted to contain an α/β hydrolase fold, which is composed of a core β-sheet made up of eight β-strands that run parallel (except for β2), interconnected by α-helices^20–22^. Like other α/β hydrolase proteins, PF3D7_0403800 also contains a catalytic triad comprised of nucleophile-histidine-acid residues (Ser-His-Asp) borne on loops that give rise to its hydrolase activity^20^. The biological function of PF3D7_0403800 in *P. falciparum* has not been studied, but it is predicted to be essential or at least important for blood-stage *P. falciparum* growth; PF3D7_0403800 is refractory to *piggyBac* transposon insertional mutagenesis^23^ and conditional mutation of the catalytic triad from Ser-His-Asp to Ser-Ala-Asp leads to defective replication^24^. The essentiality of PF3D7_0403800, combined with the potential of MMV794 to presumably inhibit it and invasion specifically, indicates that this protein represents an attractive drug target.

In this study, the C36W mutation was introduced into wild-type (3D7) parasites, leading to recapitulation of the MMV794 resistance phenotype, suggesting PF3D7_0403800 is the likely target of this compound. We also performed functional assays and lattice light-sheet microscopy to pinpoint the MoA of MMV794 in blood-stage *P. falciparum* invasion, demonstrating the compound specifically inhibits invasion when added to schizonts, later preventing merozoites from internalisation into RBCs during invasion. Lastly, we investigated the biological function of PF3D7_0403800 and its interaction with MMV794. Our findings indicate that PF3D7_0403800 localises to the rhoptries, contributes to parasite lipid metabolism, and interacts with MMV794 likely through a covalent mechanism.

## Results

### MMV794 inhibits merozoite development during schizogony

To validate the potency of MMV687794 (MMV794; Figure 1A) synthesised in-house (Figure S1) against blood-stage *P. falciparum* growth, we used the colorimetric-based 72-h lactate dehydrogenase (LDH) growth assay^25^. 3D7 parasites were treated with a two-fold titration of 10 µM MMV794 over nine concentrations, which demonstrated that the compound inhibited parasite growth over 72-h with an EC_50_ of 332 nM (Figure 1B and S2A) as previously shown^18^. To confirm that the activity of MMV794 was confined to the stage of invasion^18^, functional assays were performed using transgenic 3D7 parasites containing the bioluminescent marker Nanoluciferase (Nluc) fused to the N-terminal domain of Hyp1 (Hyp1-Nluc), which is exported into the infected RBC cytosol^26^.

**Figure 1.**
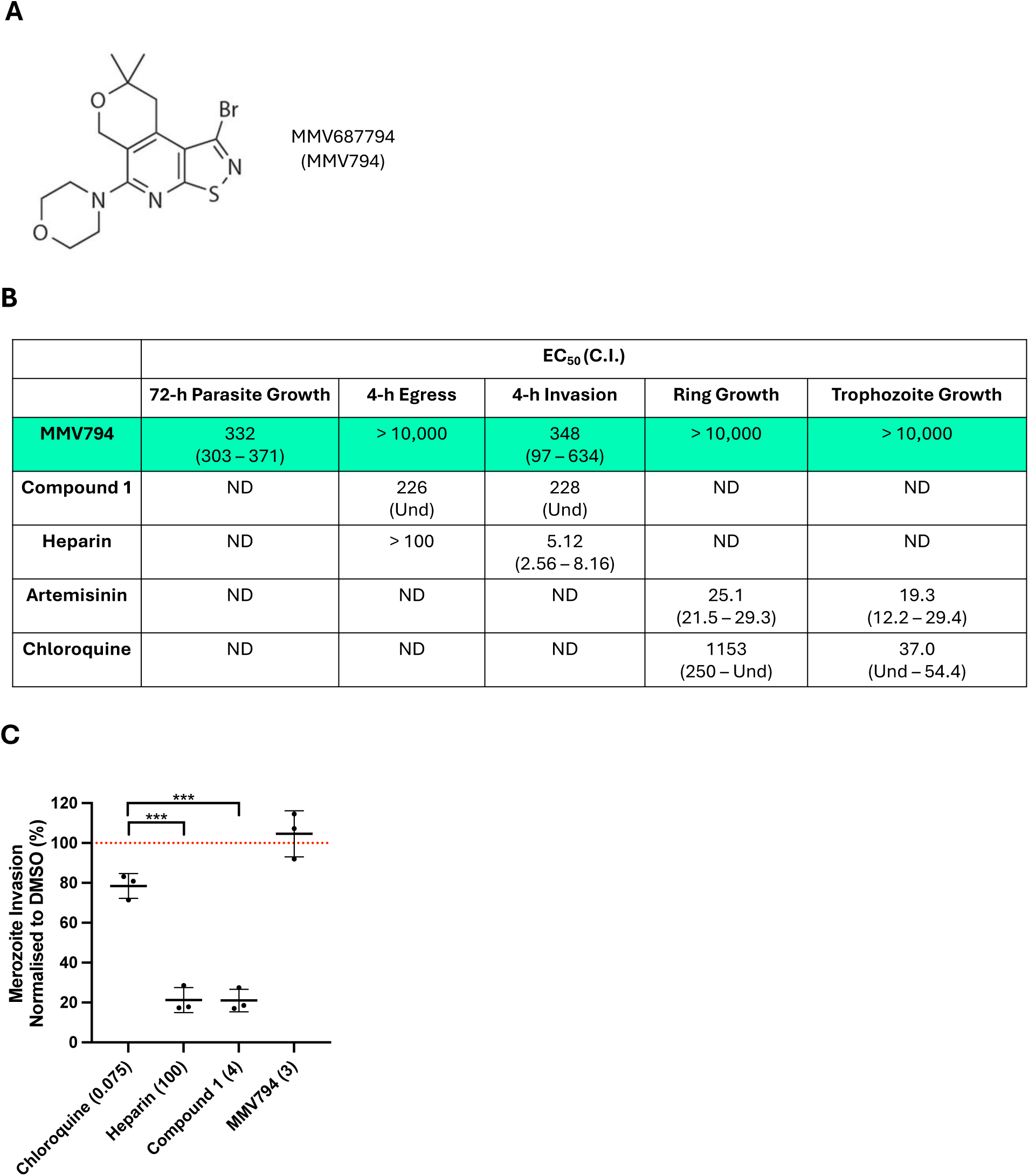
MMV794 inhibits invasion when added to schizonts. **A** Structure of MMV687794 (MMV794). **B** Tabular summary of MMV794 activity against 72-h parasite growth, 4-h egress and invasion, and ring and trophozoite growth. MMV794 inhibits invasion but not egress, ring or trophozoite growth. Compound 1 (egress inhibitor) and heparin (invasion inhibitor) were used as controls for the 4-h egress and invasion assay. Artemisinin and chloroquine (parasite growth inhibitors) were used as controls for the ring and trophozoite growth assays. Mean EC50 values were derived from dose response curves (Figure S2 and S3) and determined from three biological replicates, each with two or three technical replicates. C.I. = 95% confidence interval. ND = not determined. Und = undefined. EC50 values are in nM (except for Heparin, in µg/mL). **C** MMV794 did not inhibit merozoite invasion. Chloroquine (parasite growth inhibitor) was used as a negative control and did not inhibit merozoite invasion. Heparin and compound 1 (merozoite invasion inhibitors) both decreased merozoite invasion as expected. All values were normalised to 0.1% DMSO (vehicle control). Compound concentrations are indicated in brackets in nM (µg/mL for heparin). Error bars represent the standard deviation of the mean of three biological replicates, each with three technical replicates. Statistical analyses were performed on GraphPad Prism 10 using ordinary one-way ANOVA with Dunnett’s multiple comparison test between chloroquine (negative control) and other compounds. *** *p* < 0.001 (Heparin, *p* = 0.0009; Compound 1, *p* = 0.0007). ND = not determined. Und = undefined.

To establish the invasion-blocking potency of newly synthesised MMV794, the 4-h egress and invasion assay was performed as described^18^ (Figure S2B), whereby mature schizonts were incubated in the presence of a titration of MMV794 or control compounds (compound 1, egress inhibitor or heparin, invasion inhibitor) as well as RBC for 4 h to allow merozoites to egress and invade RBCs. During the 4-h window, compound 1 blocked egress (EC_50_ = 226 nM) and subsequent invasion (EC_50_ = 248 nM) and heparin (EC_50_ = 5.12 µg/mL) blocked invasion as expected (Figure 1B and S2C), while MMV794 did not inhibit egress but blocked invasion (EC_50_ = 348 nM) as previously reported (Figure 1B and S2D)^18^.

To confirm that MMV794 does not have a broader activity across the asexual cell cycle, ring- and trophozoite-stage growth assays^18^ (Figure S3A) were performed. In this assay, either rings (0-16 hpi) or trophozoites (18-36 hpi) were treated for 4 h with titrations of MMV794 or the control compounds artemisinin and chloroquine. Drugs were then removed, and parasites were grown for a further 24 h (rings) or 48 h (trophozoites) at 37 °C. The EC_50_ of each compound was subsequently calculated to determine the stage at which it exhibits most activity. Artemisinin inhibited the growth of rings EC_50_ = 25 nM) and trophozoites (EC_50_ = 19 nM) at similar potencies (Figure 1B and S3B) as expected^27^. Chloroquine inhibited trophozoites (EC_50_ = 37 nM) more effectively than rings (EC_50_ = 1153 nM) over 4 h, consistent with its stage-specific mechanism of action (MoA)^28^. MMV794 did not inhibit ring or trophozoite growth even at the highest concentration (10 µM) over the 4-h treatment window, indicating that the compound does not target the intracellular development of either parasite stage (Figure 1B and S3B).

The 4-h egress and invasion assay measures the effect of a compound on merozoite development within the schizont, egress from the RBC, RBC invasion, and ring-stage formation. Since MMV794 did not affect parasite egress (Figure 1B and S2D), we sought to dissect whether it directly inhibits the final stages of merozoite development, or merozoite invasion and subsequent conversion to a ring. To do this, the purified merozoite invasion assay^18,29^ (Figure S4) was performed. In this assay, merozoites are mechanically released from the iRBC and incubated for 30 mins with fresh RBCs and either MMV794 at 3 µM (10× 72-h parasite growth EC_50_) or control compounds chloroquine (growth inhibitor; 75 nM), compound 1 or heparin (invasion inhibitors; 4 µM and 100 µg/mL, respectively). After a 30-min incubation, compounds were removed, and parasites were grown for 24 h to permit Nanoluciferase expression, measured as a proxy for growth. Heparin and compound 1 blocked merozoite invasion of RBCs as expected, reducing invasion to < 20 %, while chloroquine did not significantly reduce merozoite invasion (Figure 1C). MMV794 did not inhibit merozoite invasion in this 30-min period in contrast to the 4-h invasion assay as found previously^18^, suggesting that its inhibitory effect may not be through preventing merozoite invasion or blocking early ring stage development (Figure 1C), rather exerting its inhibitory effect during schizont/merozoite maturation. Its potency in the 4-h egress/invasion assay (EC_50_ = 348 nM) closely matches that for 72-h parasite growth (EC_50_ = 332 nM; Figure 1B), suggesting MMV794 specifically targets this process. Importantly, because egress is unaffected (Figure 1B), this targeted maturation step appears to be unimportant for egress but is specifically required for invasion.

### MMV794 prevents internalisation of merozoites during invasion

Within the schizont, merozoites develop secretory organelles required for successful invasion of host cells. These organelles, termed rhoptries, micronemes, and dense granules, contain specialised proteins that are sequentially and precisely released before, during and after invasion to facilitate successful entry into the host RBC^14,30^. Since MMV794 inhibits RBC invasion by possibly disrupting merozoite development, it could disrupt the development of secretory organelles or proteins therein. To gain insight into how MMV794 treatment affects parasite development and the subsequent invasion process, live-cell imaging of late-stage schizonts (44–48 hpi) was performed in the presence of 3 µM MMV794 (10× 72-h parasite growth EC_50_) or 0.1% DMSO (vehicle control). A combined total of 47 schizont ruptures (egresses) were observed in the presence of either DMSO or MMV794 across seven imaging sessions using either brightfield or lattice light-sheet microscopy.

To ensure both treatment groups had similar numbers of surrounding RBCs to invade, the number of times merozoites released from a single RBC egress event subsequently contacting a RBC for ≥ 0.25 s was quantified and expressed as RBC contact per egress. This analysis found that merozoites contacted RBCs at a similar frequency in both treatments; each egress released merozoites that contacted RBCs 12 times (± 12.10) in DMSO and 10 times (± 11.50) in MMV794, indicating they had similar opportunities to contact and invade RBCs (Figure 2A). During normal merozoite invasion, the first obvious visual change following RBC contact is the deformation of the RBC membrane by the merozoite (Figure 2B ‘RBC deformation’), which involves ligand-receptor binding between the merozoite and RBC surfaces^31,32^. To examine whether MMV794 prevented these ligand-receptor interactions, we quantified the proportion of merozoites that deformed RBCs following contact. Post-RBC contact, a comparable proportion of merozoites deformed RBCs in either DMSO (27.7 ± 25.7%) or MMV794 (23.9 ± 29.7%) treatment (Figure 2C), indicating that MMV794 does not prevent the merozoite-RBC surface ligand-receptor interactions required for RBC deformation.

**Figure 2.**
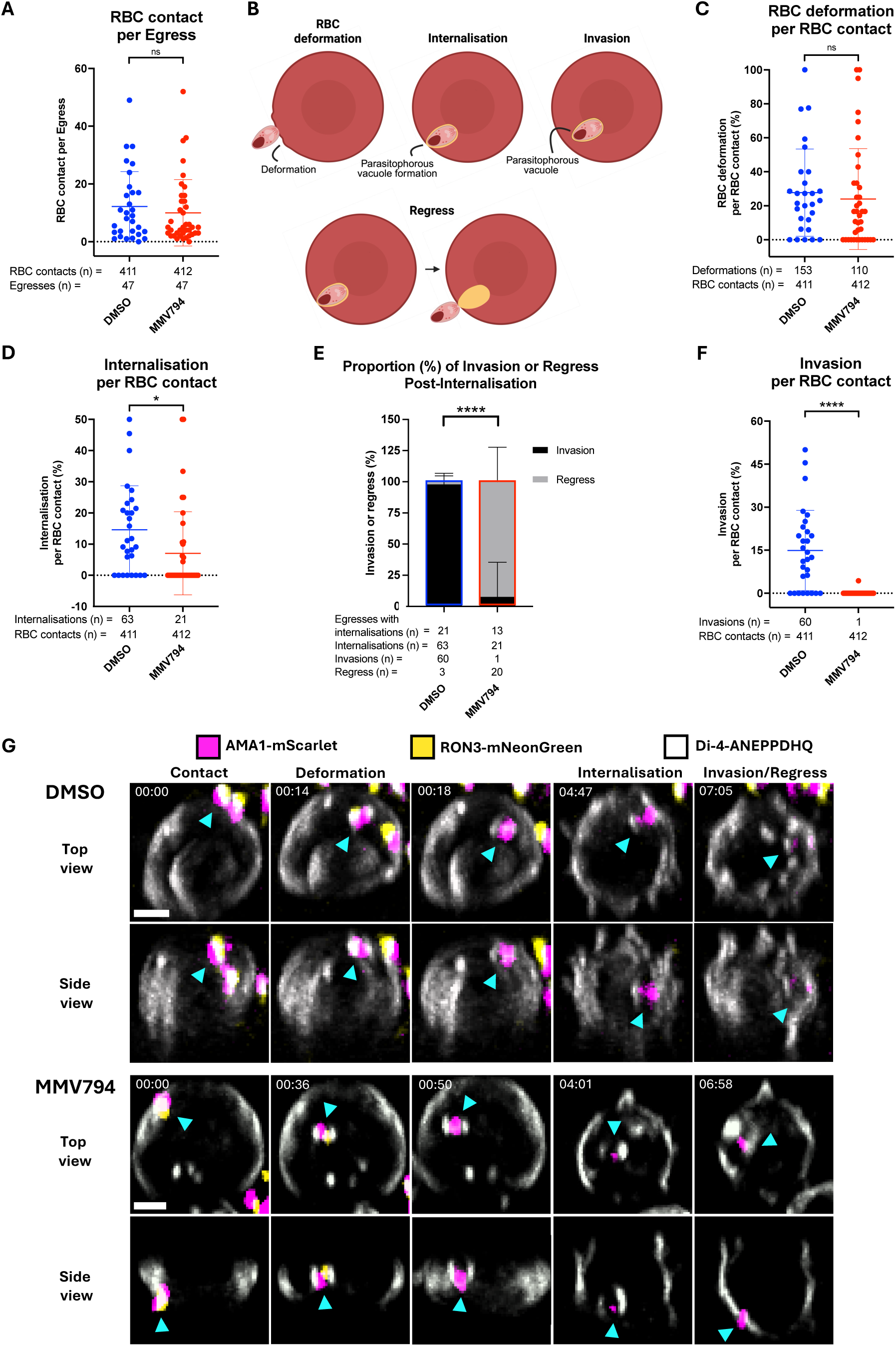
MMV794 prevents internalisation of merozoites during invasion. Live-cell imaging of schizonts was performed immediately after treatment with either DMSO vehicle control (0.1%) or MMV794 (3 µM) to capture 47 schizont ruptures (egresses) and subsequent invasion events for each treatment group. **A** The average number of times the merozoites released from one egress contacted a RBC showed that merozoites in both treatment groups contacted a RBC a similar number of times. **B** Schematics of parameters measured post-RBC contact. RBC deformation: the merozoite deforming the RBC surface after contact. Internalisation: the merozoite completely entering the RBC as it forms the parasitophorous vacuole. Invasion: the merozoite completely entering the RBC and remains within the RBC in the fully formed parasitophorous vacuole. Regress: the merozoite exits the RBC after internalisation. Figure created using BioRender.com. **C** MMV794 treatment did not affect the number of times a merozoite proceeded to deform the RBC after contact when compared to the DMSO vehicle control. **D** MMV794 treatment reduced the proportion of merozoites that proceeded to internalise themselves within the RBC after contact when compared to the DMSO vehicle control. **E** After merozoite internalisation, MMV794 treatment significantly increased the proportion merozoites that regressed from the RBC when compared to the DMSO vehicle control. **F** MMV794 treatment reduced the proportion of merozoites that remained internalised within the RBC when compared to the DMSO vehicle control. Error bars represent the standard deviation of the mean of seven biological replicates. Statistical analyses were performed on GraphPad Prism 10 using Welch’s *t*-test. * *p* < 0.05, **** *p* < 0.0001. ns = not significant. **G** Representative frames from 4D Lattice Light Sheet live cell microscopy during invasion of MMV794 or DMSO treated parasites. AMA1-mScarlet: merozoite surface (magenta). RON3-mNeonGreen: merozoite rhoptries (yellow). Di-4-ANEPPDHQ: RBC membrane (white). MMV794 treatment prevents merozoites from successfully internalising within the RBCs, leading to regress of the merozoite (indicated by arrow), which can be seen protruding from the invasion site on the RBC surface following regress. Arrows (cyan) indicate merozoite of interest. Scale bar = 2 µm.

RBC deformation is followed by moving junction formation and internalisation of the merozoite within the RBC. Moving junction formation is primarily driven by the secretion of rhoptry neck proteins, which form the RON complex on the RBC surface and bind to apical membrane antigen 1 on the merozoite surface^33–35^. The moving junction is a nexus between the merozoite and the RBC, from which the merozoite engages the glideosome to drive internalisation within the RBC and parasitophorous vacuole formation^30,36,37^. To examine whether MMV794 disrupts merozoite internalisation, defined as the process by which the merozoite completely drives itself into the RBC (Figure 2B ‘Internalisation’), we quantified the proportion of merozoites that proceeded to internalise themselves within RBCs following contact. MMV794 treatment reduced the proportion of merozoites that were internalised within RBCs (7.0 ± 13.3%; *p* = 0.0294) when compared to DMSO treatment (14.6 ± 14.1%; Figure 2D), suggesting MMV794 treatment may inhibit moving junction formation, glideosome activity or parasitophorous vacuole formation.

Post-internalisation, the RBC undergoes echinocytosis while the merozoite converts into a ring-stage parasite. If sealing of the nascent parasitophorous vacuole is impaired, the merozoite instead exits/regresses from the RBC. This phenotype has previously been observed following disruption of the moving junction^38–41^ or the glideosome^18,42^ or RBC lipid composition^32^. To examine whether MMV794 interferes with these processes and causes merozoites to regress from the RBC, all merozoites that were internalised within RBCs (Figure 2B ‘Internalisation’) were tracked for 15 mins, and the proportion that remained internalised (invasion; Figure 2B ‘Invasion’) or regressed was quantified (Figure 2B ‘Regress’). All RBC egress events leading to an internalisation/s were monitored (Figure 2E ‘Egresses with internalisations’). In the presence of MMV794, most internalised merozoites (92.31%; *p* < 0.0001) regressed from the RBC when compared with vehicle control (DMSO)-treated parasites (2.14%; Figure 2E), indicating that MMV794 blocks invasion by impairing parasitophorous vacuole sealing, likely through disruption of moving junction and/or glideosome function. This regression phenotype largely accounted for the invasion-blocking activity of MMV794, quantified as the proportion of merozoites that remained internalised (invaded) following contact (Figure 2F). MMV794 treatment significantly reduced the proportion of merozoites that remained internalised, i.e. completed invasion of a RBC (0.11 ± 0.71%; *p* < 0.0001) compared with DMSO-treated parasites (14.90 ± 13.97%; Figure 2F).

Parasite regression following MMV794 treatment is demonstrated in live-cell imaging frames of RON3-mNeonGreen/AMA1-mScarlet merozoites using lattice light-sheet microscopy, with RBC membranes stained with Di-4-ANEPPDHQ (Figure 2G). This enabled the tracking of rhoptry bulb contents (RON3-mNeonGreen), the parasite (AMA1-mScarlet), and the RBC membrane during invasion. Similar to the DMSO control (Video S1), MMV794-treated merozoites were able to deform the RBC, secrete their rhoptry bulb contents as demonstrated by the diminishing RON3-mNeonGreen signal, and subsequently proceed to internalise within the RBC. However, unlike the DMSO control, in which the merozoite remains internalised within the RBC and transitions into an amoeboid ring as the AMA1-mScarlet signal diminishes, the MMV794-treated merozoite regressed from the RBC post-internalisation (Video S2). Overall, this suggests that MMV794 prevents completion of invasion by promoting merozoite regression from the RBC through disrupted parasitophorous vacuole sealing, likely as a result of impaired moving junction and/or glideosome.

### MMV794-resistant parasites contain mutations within the *P. falciparum* α**/**β hydrolase, PF3D7_0403800 (*pfslirp*): a putative membrane-associated lipid-binding protein

To identify the potential target/s of MMV794, *in vitro* drug resistance selection was performed. Five populations (A-E) of 10^8^ wild-type mixed stage clonal 3D7 parasites were cycled on and off 3 µM MMV794 (10× 72-h parasite growth EC_50_) alongside a DMSO control well and tested for resistance using the 72-h LDH parasite growth assay. Two of the MMV794-selected parasite populations (B and C) gained ∼3-fold resistance (Figure 3A) and were subsequently cloned by limiting dilution^43^. This yielded two clonal parasite lines from population B (Pop B) and three from population C (Pop C), which retained 4-to-5-fold resistance against MMV794 when compared to wild-type 3D7 control parasites (Figure S5). To investigate the genetic changes conferring MMV794 resistance in these clonal parasites, genomic DNA was extracted from Pop B and C clones and the 3D7 parent, and whole-genome sequenced using MinION technology^42,44^. Variant analyses were performed on the aligned genomic sequences from these parasites, identifying 16 single nucleotide polymorphisms (SNPs) and five structural variants across 17 genes (Table S1). There was only one gene containing either SNPs or structural variants that was present across all resistant clones: PF3D7_0403800, a putative α/β hydrolase. Specifically, population B clones contained a 2644 bp (Pop B_F10) and 2646 bp (Pop B_F6) deletion upstream of PF3D7_0403800 (Δ5’UTR) (Table S1), while population C clones (Pop C_C6, Pop C_C8, Pop C_D6), contained a nonsynonymous SNP in PF3D7_0403800. This SNP resulted in an amino acid substitution at position 36 from cysteine to tryptophan (C36W; Table S1). Together, this suggests that PF3D7_0403800 may be involved in the MoA of MM794 and will henceforth be referred to as *pfslirp* (<u>S</u>urface-associated <u>L</u>ipid-Interacting <u>R</u>hoptry <u>P</u>rotein).

**Figure 3.**
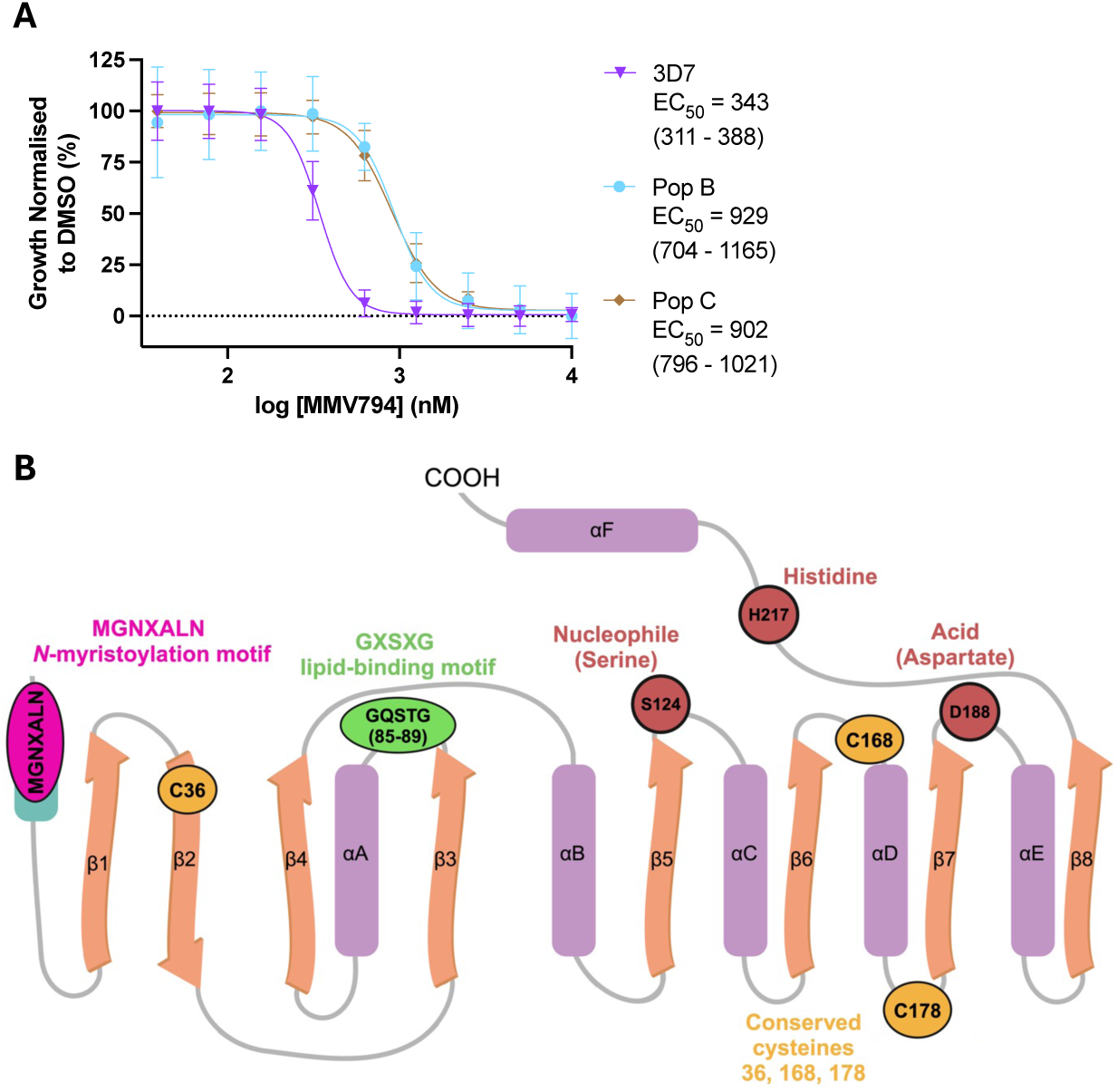
MMV794-resistant parasites contain mutations to the *P. falciparum* α/β hydrolase, *Pf*SLIRP. **A** Two MMV794-resistant parasite populations were generated (Pop B and C) by exposure of clonal 3D7 *P. falciparum* parasites to 10× 72-h parasite growth EC50 of MMV794 (3 µM). EC50s were determined by standard 72-h growth inhibition assays. Values were normalised to samples treated with 0.1% DMSO (vehicle control). Error bars represent the standard deviation of the mean of three biological replicates, each with three technical replicates. Mean of EC50 and 95% confidence interval (C.I.) values are indicated. **B** The four conserved features of PF3D7_0403800 across the *Plasmodium* genus were mapped onto the α/β hydrolase fold in 2D. Magenta denotes *N*-myristoylation motif, yellow denotes three conserved cysteines at positions 36, 168 and 178, green denotes the GXSXG lipid-binding motif and red denotes the catalytic triad composed of the nucleophile (serine 124), acid (aspartate 188) and histidine (127). Figure created using BioRender.com.

To investigate the potential biological function of *Pf*SLIRP, basic local alignment search tool (BLAST) searches^45^ and multiple sequence alignments of *Pf*SLIRP against other Apicomplexan parasites and *Plasmodium spp.* were performed. This identified potential *Pf*SLIRP orthologues likely only present in blood-borne parasites, including *Hepatocystis*, *Babesia*, *Theileiria* and other *Plasmodium spp.* (Table S2 and S3). This also revealed conserved features within the α/β hydrolase fold of *Pf*SLIRP (Figure 3B, S6 and S7), with these proteins sharing: 1) an *N*-myristoylation motif that likely localises the protein to a membrane^46^. This is concordant with a previous proteomic study showing that *Pf*SLIRP abundance decreased in the presence of an inhibitor against *N*-myristoyl transferase (*N*-myristoylation)^47^. 2) A GXSXG lipid-binding motif commonly found in other *Plasmodium* α/β hydrolase proteins that have functions in parasite lipid metabolism by acting on esters^48,49^ and phospholipids^50–53^. 3) The highly conserved nucleophile-histidine-acid catalytic triad, characteristic of the canonical α/β hydrolase fold^20,54^ (except for *P. cynomolgi* that lacks the nucleophilic serine). 4) Three highly conserved cysteines, with C36 being highly conserved only in *Hepatocystis* and *Plasmodium* parasites. Altogether, this suggests *Pf*SLIRP is a membrane-associated protein likely involved in parasite lipid metabolism, and the conserved cysteines, especially C36, may be important for this function.

### The C36W mutation in *Pf*SLIRP confers resistance to MMV794

To validate that the *Pf*SLIRP C36W mutation found in MMV794-resistant clones Pop C_C6, Pop C_C8 and Pop C_D6 conferred resistance to MMV794, CRISPR/Cas9 was used to insert this mutation into wild-type 3D7 parasites (Table S4 and S5)^55^. This genome editing approach was not pursued for introduction of the Δ5′UTR mutation as the extreme A+T-rich nature of the 5′UTR in *P. falciparum* was anticipated to preclude reliable construct assembly and homology-directed repair.

The CRISPR/Cas9 construct contained a recodonised version of *Pf*SLIRP (wildtype (C36C) or mutant (C36W)), a C-terminal 3× haemagglutinin (HA) tag, followed by a conditional knockdown *glmS* ribozyme to facilitate downstream studies of *Pf*SLIRP (Figure 4A and S8). After parasites were recovered from transfection, clonal selection by limiting dilution was performed. Integration of the construct was confirmed by PCR (Figure S9) and western blotting using antibodies to the 3× HA-tag to show that *Pf*SLIRP appeared at the expected size of 83.4 kDa (Figure 4B). The CRISPR wild-type (C36C) parasite line will be referred to as C-WT, and the mutant (C36W) as C-C36W.

**Figure 4.**
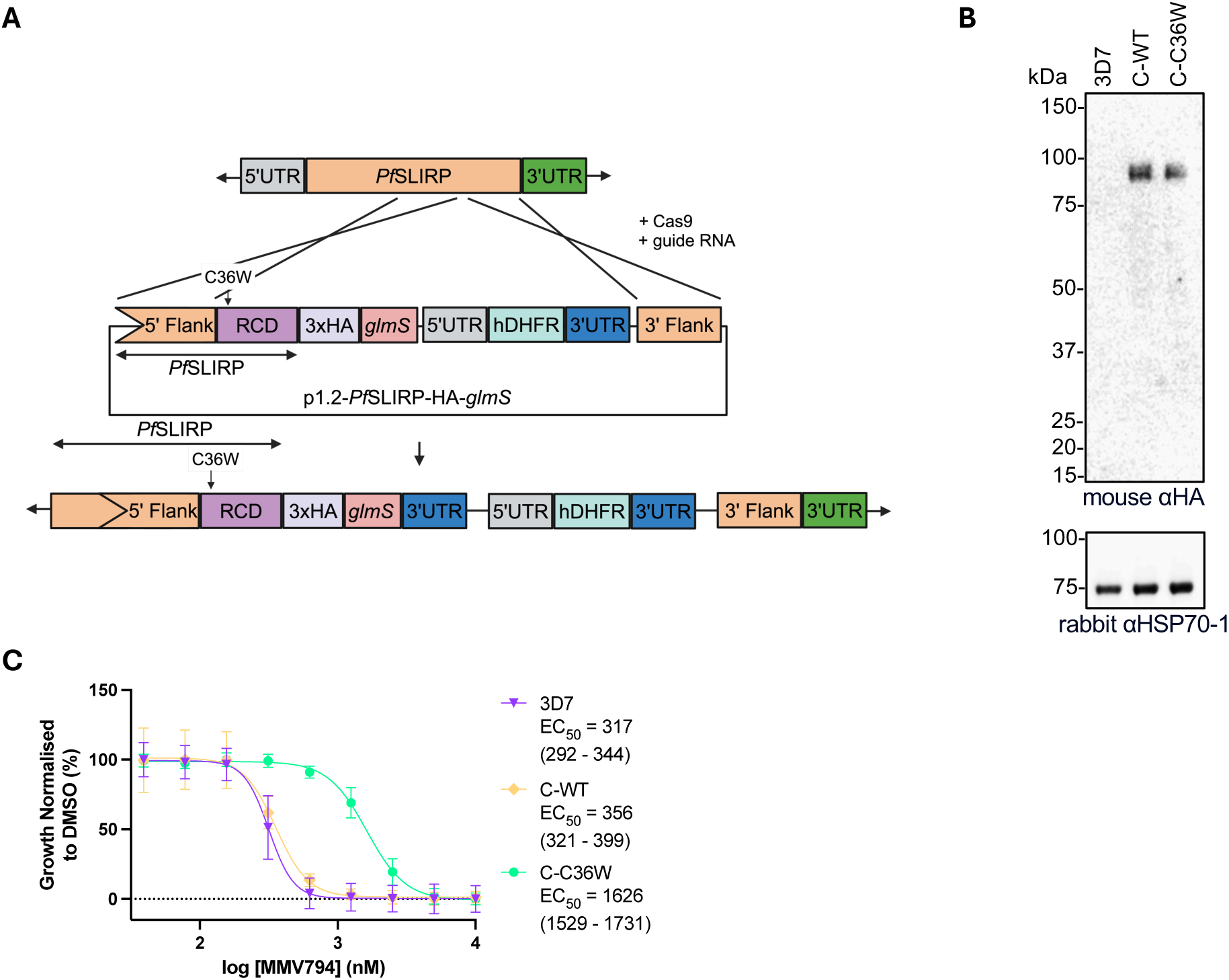
The C36W mutation in *Pf*SLIRP confers resistance against MMV794. **A** Wild-type or C36W mutant *pfslirp* genes were introduced into 3D7 parasites via CRISPR-Cas9 cloning strategy with donor plasmids p1.2-*Pf*SLIRP(WT)-HA-*glmS* (C-WT) and p1.2-*Pf*SLIRP(C36W)-HA-*glmS* (C-C36W). A 3× haemagglutinin (HA) epitope tag was included in the construct for *Pf*SLIRP detection. RCD = recodonised. *glmS* = glucosamine-6-phosphate riboswitch ribozyme. hDHFR = human dihydrofolate reductase selection marker. The double-headed arrow marks the genomic region corresponding to the *pfslirp* gene. Figure created using BioRender.com. **B** Western blot of proteins extracted from 3D7, C-WT and C-C36W demonstrate that *Pf*SLIRP was appended with HA, visualising at the expected size (83.4 kDa). Mouse anti-HA was used for probing *Pf*SLIRP, and rabbit anti-HSP70-1 was used as a loading control for the western blots. **C** Dose response curves of 72-h parasite growth assays showed that C-C36W parasites possessed 4-to-5-fold MMV794 resistance when compared to C-WT and 3D7 parasites. Values were normalised to 0.1% DMSO and EC50 values and their 95% confidence intervals in nM are indicated in brackets. Error bars represent the standard deviation of the mean of biological triplicates, each with three technical replicates.

To assess whether the C36W mutations conferred resistance to MMV794, the 72-h LDH growth assay was performed on parental 3D7, C-WT and C-C36W parasites. This demonstrated that the C36W mutation conferred 5-fold resistance to MMV794, when compared to the parental 3D7 parasites and C-WT (Figure 4C), indicating *Pf*SLIRP drives the mechanism of resistance against MMV794.

### *Pf*SLIRP associates with the rhoptry membrane at the schizont stage

The localisation of *Pf*SLIRP has not previously been reported, thus, to gain insight into the potential function of *Pf*SLIRP, its localisation was established within C-WT and C-C36W parasites via indirect immunofluorescence assays (IFAs) using anti-HA antibodies. Initial IFAs were performed on ring, trophozoite and schizont-stage C-WT parasites co-labelled with antibodies against parasitophorous vacuole membrane marker, exported protein 2 (EXP2)^56^. *Pf*SLIRP expression was only detectable in schizonts, with apical punctate labelling (Figure 5A). IFAs were also performed on C-WT schizonts co-labelled with the rhoptry neck marker RhopH3^57^. Super resolution microscopy was performed with SRRF (super-resolution radial fluctuations) analysis^58^, which revealed that *Pf*SLIRP takes on a donut-shaped distribution that appears to wrap around RhopH3 (Figure 5A). This *Pf*SLIRP localisation profile aligns with a prior observation wherein the *P. berghei Pf*SLIRP orthologue (PBANKA_1001500) also localised to the rhoptries^59^. Given that *Pf*SLIRP drives the mechanism of resistance of the invasion-blocking compound MMV794, its localisation at the rhoptries suggests that *Pf*SLIRP may play a role in invasion.

**Figure 5.**
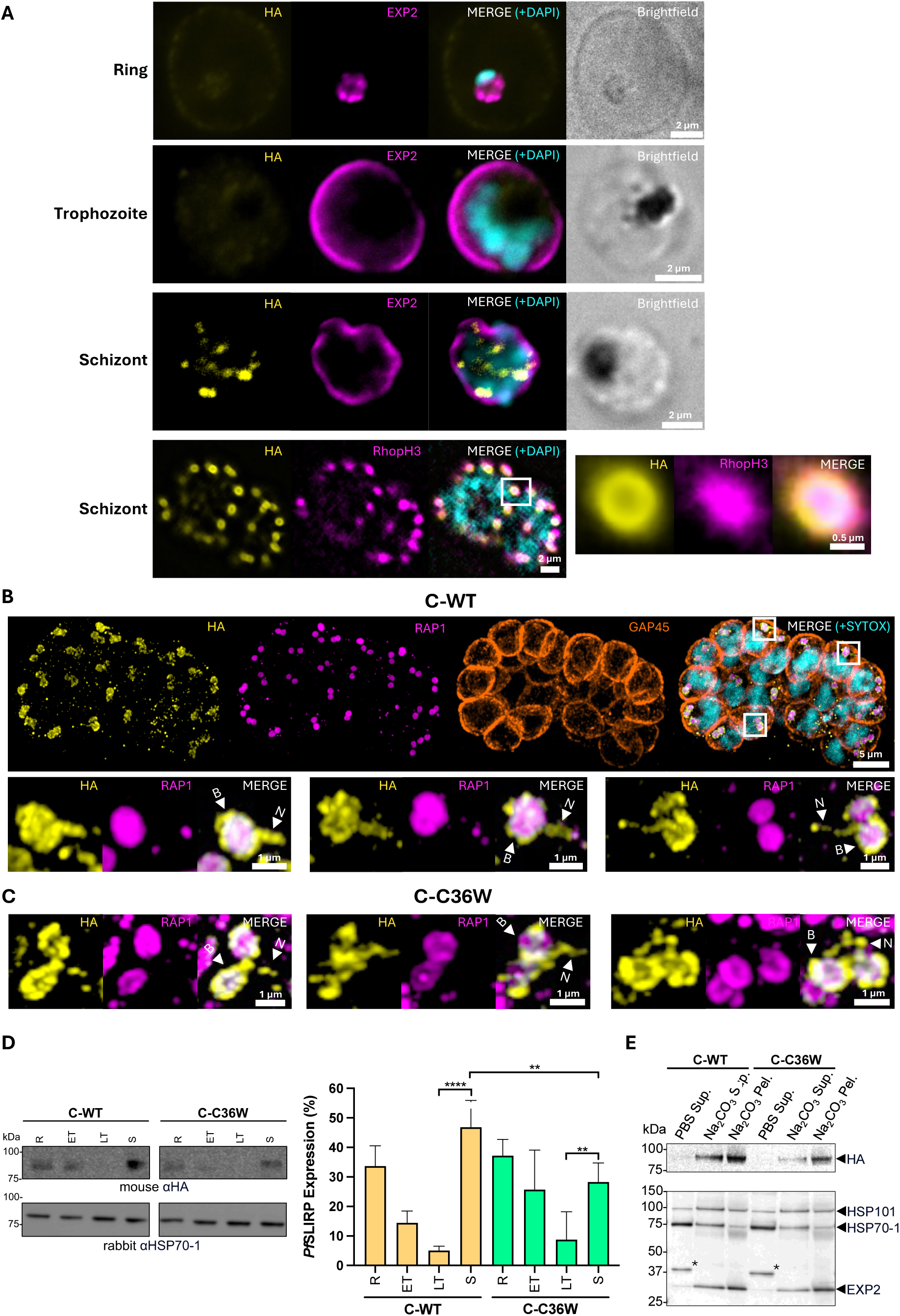
*Pf*SLIRP is highly expressed in schizonts and associates with the rhoptry membrane. **A** 2D widefield immunofluorescence assay (IFA) images of C-WT parasites showing HA-tagged *Pf*SLIRP in schizonts at the periphery of rhoptry bulbs. EXP2 marks the parasitophorous vacuole membrane, RhopH3 marks rhoptry bulbs, and DNA is stained with DAPI. Super-Resolution Radial Fluctuations (SRRF) processing was applied to enhance spatial resolution. The white box in the merged schizont image indicate zoomed area to the right. **B** Confocal maximum intensity projections of expanded C-WT schizonts illustrating wild-type *Pf*SLIRP–HA localisation at the rhoptry membrane. RAP1 marks rhoptry bulbs, GAP45 labels the merozoite periphery, and SYTOX™ blue stains DNA. The white boxes in the merged image indicate zoomed areas at the bottom. B = rhoptry bulb, N = rhoptry neck. **C** Confocal maximum intensity projections of rhoptries from expanded C-C36W schizonts (Figure S11) illustrating C36W *Pf*SLIRP-HA localisation at the rhoptry membrane. **D** Western blot of parasite proteins from ring (R), early trophozoite (ET), late trophozoite (LT), and schizont stages, probed with anti-HA to detect *Pf*SLIRP. HSP70-1 serves as a loading control. Densitometry values were normalised to HSP70-1, a constitutively-expressed housekeeping protein. *Pf*SLIRP expression peaks at the schizont stage for C-WT, and the ring stage for C-C36W. Western blot is representative of three biological replicates (refer to Figure S12 for all replicates). Error bars represent the standard deviation of the mean of three biological replicates, each with two technical replicates. Statistical analyses were performed with GraphPad Prism 10, with Welch’s *t*-test between the late trophozoite and schizont stages within both parasite lines, and between the same stages across both parasite lines. ** *p* < 0.01, **** *p* < 0.0001. No bar indicates not significant. **E** Sequential lysis of C-WT and C-C36W schizont saponin pellets by freeze–thaw and sodium carbonate extraction. Western blot shows *Pf*SLIRP is found in the carbonate soluble (Na2CO3 Sup.) and insoluble (Na2CO3 Pel.) fractions but not in the PBS soluble fraction (PBS Sup.). Control proteins included soluble protein *Pf*HSP70-1, peripheral membrane protein *Pf*HSP101, and integral membrane protein *Pf*EXP2. Mouse anti-HA was used for *Pf*SLIRP detection; rabbit antibodies were used to detect the control proteins. * denotes cross-reactive bands produced by rabbit HSP70-1 antibody (refer to Figure S14 for all cross-reactive bands produced by the antibody).

Rhoptry proteins commonly localise to specific rhoptry zones e.g. within the rhoptry neck or bulb, or on the surface. Therefore, to determine the suborganellar localisation of *Pf*SLIRP, ultrastructure expansion microscopy (U-ExM)^60–62^ was applied to C-WT schizonts. IFAs were then performed on these samples and imaged using a LSM980 with Airyscan 2 (Zeiss). As an experimental control, IFAs performed on U-ExM samples of C-WT schizonts using established parasite markers such as NHS Ester (protein density) and Glideosome-Associated Protein 45 (GAP45) (inner membrane complex/merozoite periphery) were consistent with previous studies^36,61^. NHS ester most densely stains the protein-rich rhoptries at the wider end of the merozoite, while GAP45 marks the merozoite periphery (Figure S10).

To determine the suborganellar localisation of *Pf*SLIRP, C-WT schizonts were labelled with antibodies to HA (*Pf*SLIRP), RAP1 (rhoptry bulb), and GAP45, then with secondary antibodies conjugated to Alexa Fluor 488, 594 and 647, respectively. SYTOX™ BLUE was used to stain the nuclei. *Pf*SLIRP signal surrounded RAP1, thus localising to the surface of the rhoptry bulb (Figure 5B). Interestingly, the *Pf*SLIRP signal extended beyond that of RAP1 and therefore it also localises to the rhoptry neck (Figure 5B). To examine whether the C36W mutation alters the localisation of *Pf*SLIRP and therefore its potential function, the same U-ExM IFAs were performed. Resultant images revealed that the C36W *Pf*SLIRP protein had the same localisation profile as the wild type (Figure 5C). Collectively, this suggests *Pf*SLIRP has an important role during schizogony at the rhoptry membrane, and the C36 cysteine does not appear to be individually essential for its localisation.

To further inform the biological function of *Pf*SLIRP, its blood-stage profile was established by extracting proteins from C-WT and C-C36W parasites at various stages, including rings (0–12 hpi), early trophozoites (18–24 hpi), late trophozoites (30–36 hpi) and schizonts (40–46 hpi). To generate the parasites used in these assays, parasite cultures were sorbitol synchronised for harvesting immediately (rings) or later within the same cycle after delaying at RT (trophozoites). Schizonts were also treated with the reversible protein kinase G (PKG) inhibitor ML10^63^ to prevent egress. *Pf*SLIRP abundance was visualised via western blotting using antibodies to HA and the constitutively expressed housekeeping protein (HSP70-1), which served as a loading control (Figure 5D). The relative expression (%) of *Pf*SLIRP was calculated by normalising its signal to HSP70-1, which was converted to a percentage of total *Pf*SLIRP signal across the entire blot. This revealed that wild-type *Pf*SLIRP (C-WT) is most highly expressed in schizonts with 47% of total *Pf*SLIRP expression found in this stage, followed by ring (34%), early trophozoite (14%) and late trophozoite (5%) (Figure 5D). This expression profile agrees with previous transcriptomics studies^64–67^ but contrasts with our localisation results, which showed that *Pf*SLIRP is only detectable in schizonts, perhaps because it requires the formation of rhoptries to concentrate its localisation signal.

To examine whether the C36W mutation modifies the blood-stage expression profile of *Pf*SLIRP, the same western blot analysis was performed. Interestingly, the main difference detected was that C36W *Pf*SLIRP was most highly expressed in rings (37% of total expression), driven by a strong reduction in schizonts expression to 28% (Figure 5D; C-WT vs. C-C36W, *p* = 0.0026). Again, expression was intermediate in early trophozoites (26%) and lowest in late trophozoites (9%) (Figure 5D). Taken together, *Pf*SLIRP expression most likely peaks in the schizont stage, and the C36W mutation may reduce its abundance at this stage.

*Pf*SLIRP is predicted to contain an *N*-myristoylation motif and is enriched with other *N*-myristoylated proteins^47^. This post-translational modification associates proteins with membranes in the absence of a transmembrane domain. To investigate the membrane association of *Pf*SLIRP, carbonate extraction was performed on C-WT and C-C36W parasites. Saponin pellets were harvested from schizonts and sequentially lysed, first via freeze-thaw to rupture cell membranes and release soluble proteins (PBS supernatant). The resulting pellets were then lysed by incubation with sodium carbonate to release peripheral membrane proteins (Na_2_CO_3_ supernatant), with the final pellet containing integral membrane proteins (Na_2_CO_3_ pellet). Loading controls for each fraction included the soluble protein HSP70-1^68,69^, peripheral membrane protein HSP101 and integral membrane protein EXP2^56^. The controls behaved as expected while *Pf*SLIRP (wild type and C36W) was detected in the carbonate soluble and insoluble fractions, appearing more concentrated in the latter (Figure 5E). This was consistent with that expected of an *N*-myristoylated protein.

### Conditional knockdown of *Pf*SLIRP does not affect parasite fitness

To investigate whether *Pf*SLIRP is essential for blood-stage *P. falciparum*, parasite growth was examined by microscopy of C-WT cultures treated with glucosamine (GlcN) to conditionally knockdown *Pf*SLIRP expression. C-WT and C-36W knockdown efficiency was investigated by treating parasites with 2.5mM glucosamine for 1.5 cell cycles (from early trophozoites to late schizonts the following cycle), followed by western blotting and densitometry analysis. HSP70-1 was used as a loading control. These data revealed that C-WT and C-C36W parasites achieved 87% *Pf*SLIRP knockdown following this treatment (Figure 6A and B). To investigate if *Pf*SLIRP knockdown affected the growth of the parasite, parasite growth rate ± 2.5 mM GlcN was monitored over 264 h (5.5 cell cycles) via brightfield microscopy of Giemsa-stained thin blood smears. The fold change in parasitaemia of the C-WT ± 2.5 mM GlcN cultures was determined by dividing the total parasitaemia at every cycle (48 h) by the parasitaemia of the previous cycle, which was determined through microscopic counts of Giemsa-stained thin blood smears. This showed that the parasites grew at comparable rates, with similar fold changes in parasitaemia in untreated C-WT (8.58 ± 4.84-fold change) versus treated C-WT (5.98 ± 3.22-fold change) cultures at 240 h (Figure 6C). Images taken illustrate that 2.5 mM GlcN treatment of C-WT cultures did not affect parasite growth rate or gross morphology throughout the 264-h treatment period (Figure 6D). These results suggest that 87% *Pf*SLIRP knockdown may be insufficient to observe a growth or gross morphological defect, with low levels of *Pf*SLIRP expression remaining adequate to sustain normal levels of growth.

**Figure 6.**
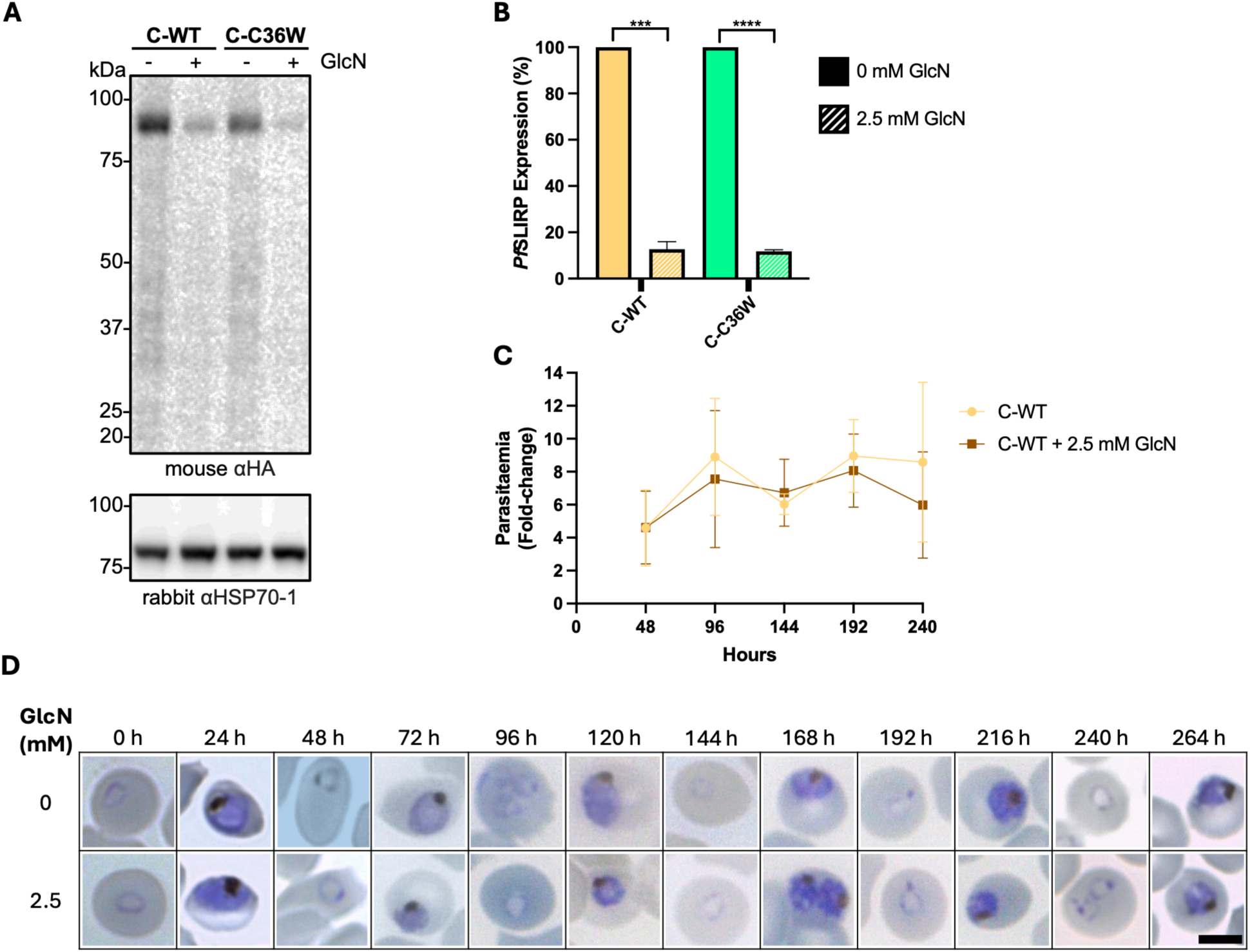
Conditional knockdown of *Pf*SLIRP does not aXect parasite growth or gross morphology. **A** Western blot of proteins extracted from C-WT and C-C36W parasites ± 2.5 mM glucosamine (GlcN) for 72 h (early trophozoites to schizont) demonstrate *Pf*SLIRP knockdown. Western blots are representative of three biological replicates (refer to Figure S15 for all biological replicates). Mouse anti-HA was used to probe *PfS*LIRP, and rabbit anti-HSP70-1 was used as a loading control. **B** Densitometry analysis of *Pf*SLIRP (mouse anti-HA) normalised to HSP70-1 demonstrates that 87% knockdown of *Pf*SLIRP can be achieved in both C-WT and C-C36W parasites. Error bars represent the mean of three biological replicates. Statistical analyses were performed on GraphPad Prism 10 with Welch’s *t*-test for each parasite line ± 2.5 mM. *** *p* < 0.001, **** *p* < 0.0001 (C-WT, *p* = 0.0005; C-C36W, *p* < 0.0001). No bar indicates not significant. **C** The fold-change in parasitaemia of C-WT parasites ± 2.5 mM GlcN for 264 h were measured by examination of **D** Giemsa-stained thin blood smears every 24 h, showing the C-WT parasites grew comparably ± 2.5 mM GlcN every cycle (48 h). Error bars represent the standard deviation of three biological replicates. Scale bar = 5 µm.

### The C36W mutation in *Pf*SLIRP alters parasite lipid homeostasis

*Pf*SLIRP contains a predicted lipid-binding motif GXSXG (GQSTQ). Therefore, we hypothesised that while the conditional knockdown of *Pf*SLIRP did not affect parasite growth rate or gross morphology, it may impact parasite lipid metabolism. To investigate this, we performed mass spectrometry-based lipidomic analyses on total lipids extracted from C-WT and C-C36W parasites treated ± 2.5 mM GlcN for 72 h to analyse lipid abundance. Since *Pf*SLIRP localises to the rhoptry surface and mutation of its C36 confers resistance to the invasion-blocking compound MMV794, we hypothesised that *Pf*SLIRP likely plays a role in parasite invasion of RBCs. Thus, alterations to the lipidome would be observable in parasites that had recently invaded new RBCs. To test this, 3D7, C-WT, and C-C36W parasites were sorbitol-synchronised and cultured to the early trophozoite stage (18-24 hpi), then treated ± 2.5 mM GlcN for 72 h until reaching the schizont stage in the subsequent cycle (Figure 7A). To prevent the second-cycle schizonts rupturing, parasites were treated for 16 h with 30 nM of the reversible egress inhibitor ML10, 56 h into the 72-h treatment window at the mid-trophozoite stage (24-30 hpi) (Figure 7A). Stalled schizont cultures were subsequently washed with cRPMI to remove ML10 and allow schizont rupture and release of merozoites. After a 4-h invasion period into fresh RBCs, parasites were pelleted, quenched, lysed with saponin, and metabolites extracted for LC-MS analysis. To examine the lipidome of C-WT and C-C36W ± 2.5 mM GlcN parasites, the relative abundance (Mol %) of 12 lipid classes was calculated using the internal standard normalisation method as previously reported^70–72^. This revealed that *Pf*SLIRP knockdown did not significantly alter the lipidome in either C-WT or C-C36W parasites (Figure 7B and C). A control assay also confirmed that the metabolome of C-WT parasites was comparable to that of 3D7, and the 3D7 metabolome was unaffected by 2.5 mM GlcN (Figure S16). These data suggest that *Pf*SLIRP may remain partially functional in lipid metabolism at low expression levels, or that changes solely due to *Pf*SLIRP knockdown are too subtle to detect using whole-parasite lipid extracts, given its rhoptry localisation.

**Figure 7.**
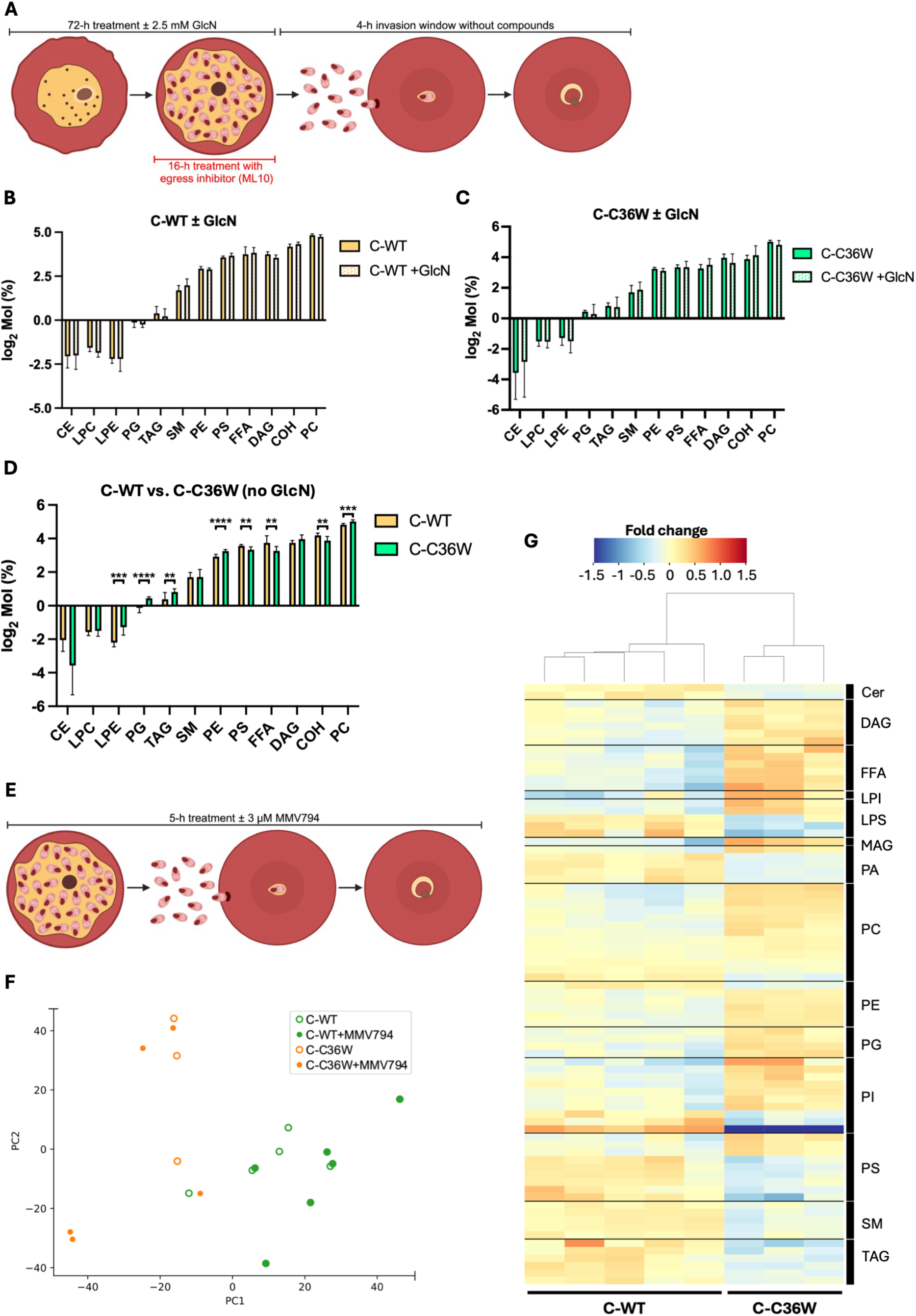
Lipidomics analyses reveal that the C36W mutation in *Pf*SLIRP perturbs the parasite lipidome. **A** Early trophozoite C-WT and C-C36W parasites were treated ± 2.5 mM glucosamine (GlcN) for 72 h. At 56 h, ML10 was added for 16 h to stall parasites at the schizont stage. After GlcN/ML10 removal at 72 h, parasites were allowed to egress and invade for 4 h before lipids were harvested for liquid chromatography-mass spectrometry (LC-MS) analysis. Figure created using BioRender.com. Relative abundance (log2 Mol%) of 12 lipid classes showed that *Pf*SLIRP knockdown (+ GlcN) did not significantly alter lipid composition between **B** C-WT or **C** C-C36W parasites +/- GlcN. **D** Eight lipid classes showed significant abundance changes between C-WT and C-C36W parasites. Error bars represent the standard deviation of five biological replicates, each with two technical replicates. Statistical analyses were performed in GraphPad Prism 10 using Welch’s *t*-test (C-WT vs C-WT + GlcN, C-C36W vs C-C36W + GlcN, and C-WT vs C-C36W). ** *p* < 0.01, *** *p* < 0.001 **** *p* < 0.0001. No bar indicates not significant. Lipid classes: cholesteryl-esters (CE), lysophosphatidylcholine (LPC), lysophosphatidylethanolamine (LPE), phosphatidylglycerol (PG), sphingomyelin (SM), triacylglycerol (TAG), cholesterol (COH), diacylglycerol (DAG), free fatty acid (FFA), phosphatidylcholine (PC), phosphatidylethanolamine (PE) and phosphatidylserine (PS). **E** Purified schizonts were treated ± 3 µM MMV794 for 5 h during egress and invasion, then harvested for metabolomic analysis. Figure created using BioRender.com **F** Principal component analysis of global metabolites from untreated (open) or treated (filled) C-WT (orange) and C-C36W (green) parasites shows distinct metabolomes between C-WT and C-C36W, with MMV794 having no major effect. Points represent individual biological replicates. **G** Heatmap of relative metabolite abundance of 79 significantly altered lipids (*p* < 0.02) between untreated C-WT and C-C36W. Hierarchical clustering indicates sample similarity. Lipids are organised vertically by class and within each class by abundance profile. Additional lipid classes: phosphatidylinositol (PI), lysophosphatidylserine (LPS), ceramide (Cer), phosphatidic acid (PA), monoglyceride (MG). Each column represents one biological replicate. Colour scale represents log2 fold change.

In contrast, the lipid content and homeostasis differed significantly between C-WT and C-C36W parasites, with C-C36W showing significantly higher levels of lysophosphatidylethanolamine (LPE), phosphatidylglycerol (PG), triacylglycerol (TAG), phosphatidylethanolamine (PE) and phosphatidylcholine (PC) but significantly lower levels of phosphatidylserine (PS), free fatty acid (FFA) and cholesterol (COH) (Figure 7D). These broad shifts in lipid class abundance suggest that the C36W mutation disrupts *Pf*SLIRP function and perturbs parasite lipid metabolism more extensively than *Pf*SLIRP knockdown alone.

To dissect the effect of the C36W mutation and MMV794 on the parasite lipidome, further untargeted LC-MS-based metabolomic analyses were conducted on total lipids extracted from C-WT and C-C36W parasites ± 3 µM MMV794. Briefly, purified C-WT and C-C36W schizonts were allowed to invade RBCs for 5 h ± 3 µM MMV794, and metabolites were extracted from the resulting iRBCs (Figure 7E). Approximately 3000 metabolites were putatively identified by exact mass and predicted retention time^73^. Consistent with the above finding that C-C36W parasites have a significantly perturbed lipidome, principal component analysis (PCA) shows the metabolic signatures for untreated C-WT and C-C36W are distinct with clear separation (Figure 7F). To calculate which metabolites were perturbed due to the C36W mutation, the fold change in the metabolites of C-C36W parasites was determined relative to those in C-WT parasites. This detected 260 significantly perturbed putative metabolites in C-C36W parasites (Figure S17). 113 of the perturbed features were putatively annotated as phospholipids including phosphatidylcholine (PC), phosphatidylethanolamine (PE), phosphatidylserine (PS) and phosphatidylinositol (PI) (Figure S17), representing a clear enrichment of this lipid class. Other lipid classes, including fatty acids and lysophospholipids were also abundant, whereas clear enrichment of other metabolite classes was not apparent. We therefore used established trends in chromatographic retention to confirm and extend the identification of lipid species in this dataset. This resulted in 713 high-confidence lipid annotations including phospholipids, neutral lipids, lysophospholipids, sphingolipids and FFAs. Of these, 79 showed significantly perturbed abundance in C-C36W parasites, and this included phospholipids (PC, PE, PS, PI, phosphatidic acid (PA), PG, PI, PS), sphingolipids (ceramide (Cer), sphingomyelin (SM)), neutral lipids (DAG, monoacylglycerol (MAG), TAG) and lysophospholipids (lysophosphatidylinositol (LPI), LPS) and FFAs (Fig 7G). When compared with the lipid perturbations measured in our targeted lipidomics experiment (Figure 7D), we observed robust correlations in measured fold-changes for PC, lysophosphatidylcholine (LPC) and FFAs, indicative of consistent lipidomic perturbations involving these lipid classes across these two experiments.

PCA of parasite metabolomes also illustrated that addition of MMV794 did not further alter the metabolic profiles of C-WT and C-C36W under these conditions (Figure 7F). The results suggest that MMV794 does not affect the parasite metabolome despite blocking invasion within this 5-h invasion window. Alternatively, it is more likely that MMV794 acts on lipid pathways within the rhoptries involving *Pf*SLIRP, inducing changes that may be critical for invasion but are too subtle to be detected from whole parasite lipid extracts, as observed above in parasites with *Pf*SLIRP knockdown. Furthermore, the phospholipid perturbations in C-C36W parasites were not recapitulated by MMV794 or GlcN treatment of C-WT parasites. These data indicate that activities of other proteins/enzymes involved in parasite lipid metabolism may have also changed in MMV794-resistant parasites, potentially serving as compensatory mechanisms to counteract the compound.

### MMV794-resistance mutations perturb the parasite proteome and decrease *Pf*SLIRP expression

To investigate whether the mutations in *Pf*SLIRP perturb the abundance of *Pf*SLIRP and potentially other proteins, we performed untargeted proteomics analyses of MMV794-resistant parasite lines Pop B_F10 (Δ5’UTR; Figure S5), Pop C_C6 (C36W; Figure S5), C-C36W, C-WT and parental 3D7 parasites. Briefly, segmented schizont lysates were subjected to extraction and purification for LC-MS/MS analysis, and relative protein abundances were determined using label-free quantification via data-independent acquisition (LFQ-DIA) proteomics. This identified 5,887 proteins, of which 4,358 were from *Plasmodium* (Table S6). PCA and hierarchical clustering analyses were performed on the relative abundances of these proteins in the different parasite lines. Similar proteomic signatures were detected between the *in vitro* generated MMV794-resistant parasites (Pop C_C6 and Pop B_F10), and the transgenic parasites (C-WT and C-C36W), while the parental 3D7 parasites had a distinct proteomic signature (Figure 8A and B). A subset of 28 proteins was detected as significantly perturbed based on pairwise comparisons across the five parasite lines. Statistical significance was assessed using limma empirical Bayes–moderated *t-*tests for each defined contrast (*p* < 0.05, log_2_ fold change ≥ 0.25), and these proteins are represented in a separate heatmap (Figure 8C). Specifically, six proteins were upregulated in all mutants (Pop B_F10, Pop C_C6 and C-C36W) compared to C-WT and 3D7 parasites, while 22 were downregulated (Figure 8C). *Pf*SLIRP abundance was significantly decreased in C-C36W (*p* = 0.0117), Pop B_F10 (*p* < 0.0001), and Pop C_C6 (*p* < 0.0001) compared to 3D7 parasites, indicating that MMV794 resistance-conferring mutations decrease the expression of the protein (Figure 8D).

**Figure 8.**
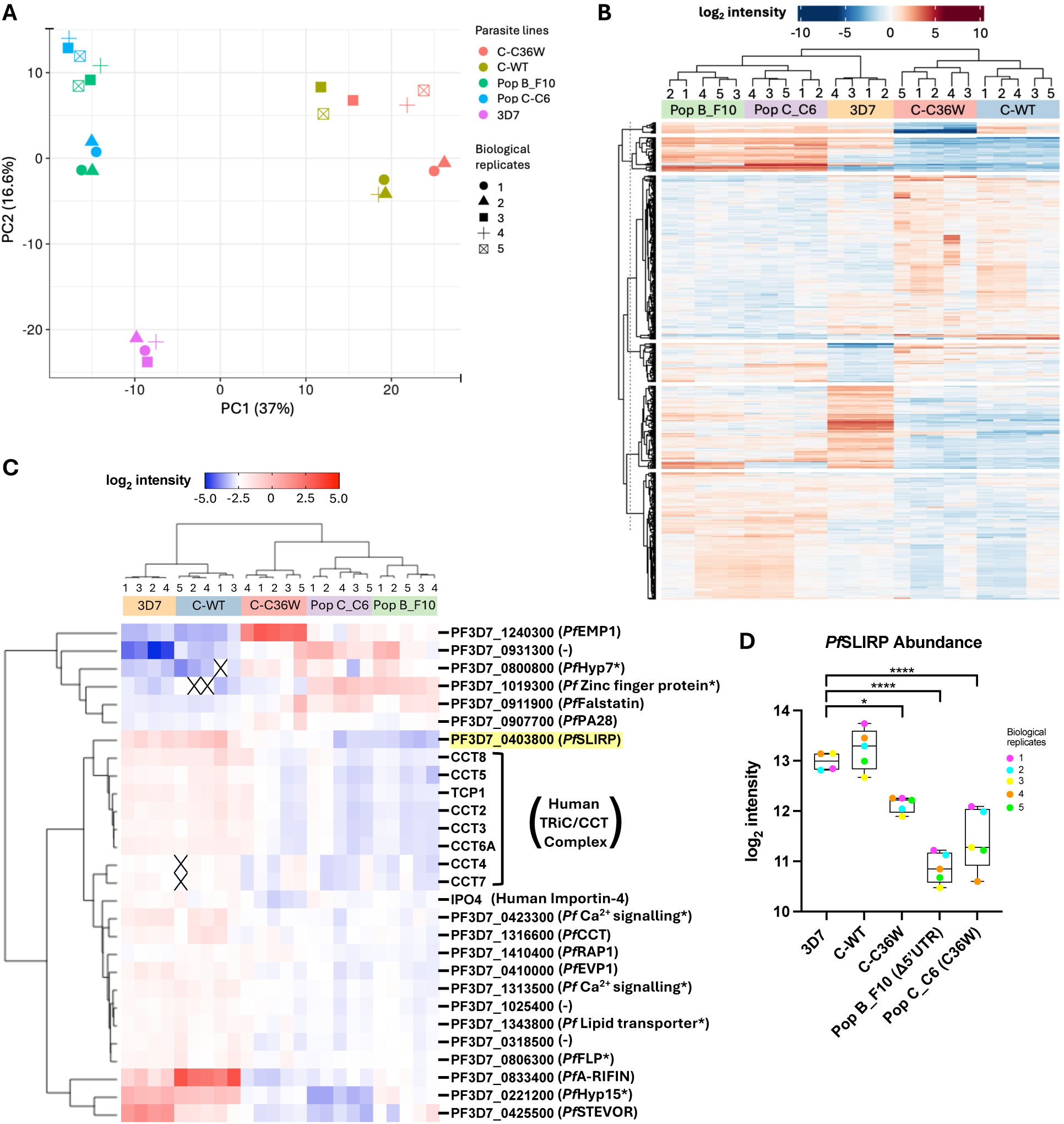
MMV794-resistance mutations perturb the parasite proteome and decrease *Pf*SLIRP expression. Untargeted comparative proteome profiling of 3D7, Pop B_F10, Pop C_C6, C-WT and C-C36W parasites (purified schizonts). **A** Principal component analysis of global proteome profiles from the different parasite lines demonstrate that similar profiles are observed between Pop B_F10 and Pop C_C6 parasites, C-WT and C-C36W parasites, while 3D7 parasites are different from all the other parasite lines. **B** Hierarchical clustering analysis of the relative abundance of 5887 proteins demonstrate different proteomes across the five parasite lines, with 28 proteins being significantly perturbed. Vertical clustering shows sample similarities and horizontal clustering shows the relative abundance of the 5887 proteins. **C** Hierarchical clustering analysis of the relative abundance of the 28 significantly perturbed proteins (including *Pf*SLIRP; highlighted in yellow). Vertical clustering shows sample similarities and horizontal clustering shows the relative abundance of the 28 significantly perturbed proteins. Each column represents a single biological replicate. The colour scale bar represents log2 intensity (median-centred). X = not identified. Protein names/functions are indicated in brackets. (*) = uncharacterised proteins. (-) = unknown. **D** Across the five parasite lines, *Pf*SLIRP abundance (log2 intensity) is significantly decreased in C-C36W, Pop B_F10 and Pop C_C6 parasites, when compared to 3D7 parasites. Statistical analyses were performed on GraphPad Prism 10 with ordinary one-way ANOVA with Dunnett’s multiple comparison test between 3D7 and other parasite lines. * *p* < 0.05, **** *p* < 0.0001.

This proteomics analysis also revealed that *Pf*SLIRP mutants dysregulated parasite proteins associated with pathways in: 1) lipid metabolism, as they have reduced abundance of a lipid transfer protein (PF3D7_1343800; *Pf*VPS13LP2^74^) and *Pf*CCT, the rate-limiting enzyme in phosphatidylcholine synthesis^75^ (Figure 8C). 2) Vesicular trafficking and fusion, underpinned by downregulated *Pf*RAP1^76^ and *Pf*EVP1^77^, which facilitate trafficking at the rhoptries and tubovesicular network, respectively (Figure 8C). 3) Invasion, by upregulating the host cysteine protease inhibitor Falstatin^78^, and downregulating a ferlin-like protein (*Pf*FLP) important for egress^79^ (Figure 8C). These changes likely reflect compensatory adaptations within the parasites to counteract the MoA of MMV794, which blocks invasion by perturbing *Pf*SLIRP, a lipid-interacting protein at the rhoptries. We also observed perturbations in other proteins, of which only a few have been studied, including virulence factors (STEVOR, A-RIFIN, *Pf*EMP1), the human TRiC/CCT complex, and the 20S proteasome regulator *Pf*PA28 (Figure 8C).

### MMV794 demonstrates target engagement with *Pf*SLIRP in blood-stage *Plasmodium falciparum*

While the *Pf*SLIRP C36W mutation mediates resistance to MMV794, this does not establish that MMV794 directly targets *Pf*SLIRP. To investigate potential drug-target engagement between MMV794 and *Pf*SLIRP, we performed drug sensitisation assays in which *Pf*SLIRP expression was reduced in the presence of MMV794. If *Pf*SLIRP was the sole direct target of MMV794, reduced *Pf*SLIRP levels would be expected to increase parasite sensitivity to the compound. To test this, parasite cultures were pre-treated ± 2.5 mM GlcN for 48 h (ring to ring in the following cycle). Subsequently, GlcN-treated parasites were maintained on 2.5 mM GlcN for the 72-h assay, resulting in a total of 120 h of GlcN exposure. We previously achieved 87% *Pf*SLIRP knockdown (Figure 6A and B) following 72-h treatment with 2.5 mM GlcN. This 120-h GlcN treatment ensured maximal *Pf*SLIRP knockdown that may be sufficient to produce an observable change in the MMV794 parasite growth EC. The growth EC was calculated for each treatment group. MMV794 exhibited similar EC s against 3D7 parasites treated ± 2.5 mM GlcN, indicating that parental 3D7 parasites remained sensitive to MMV794 and that 120-h GlcN treatment did not significantly affect parasite growth (Figure 9A). As expected, C-C36W parasites without *Pf*SLIRP knockdown (-GlcN) remained 5–6-fold more resistant to MMV794 than 3D7 and C-WT -GlcN parasites (Figure 9A). Unexpectedly, in C-WT parasites, *Pf*SLIRP knockdown conferred 4-fold resistance to MMV794 when compared to the C-WT -GlcN control (Figure 9A; *p* < 0.0001), illustrating that reduced *Pf*SLIRP expression is linked to MMV794 resistance. Despite this shift in EC_50_, knockdown of *Pf*SLIRP does not fully recapitulate the shift in EC_50_ seen in C-C36W -GlcN parasites when compared to C-WT or 3D7 (-GlcN), suggesting that while reduced levels of *Pf*SLIRP do provide MMV794 resistance, the C36W mutation further enhances this drug resistance phenotype (Figure 9A). Importantly, knockdown of *Pf*SLIRP in C-C36W parasites did not alter MMV794 sensitivity, likely because they have already reached maximal resistance to the compound given their combination of reduced *Pf*SLIRP abundance and the C36W mutation (Figure 9A).

**Figure 9.**
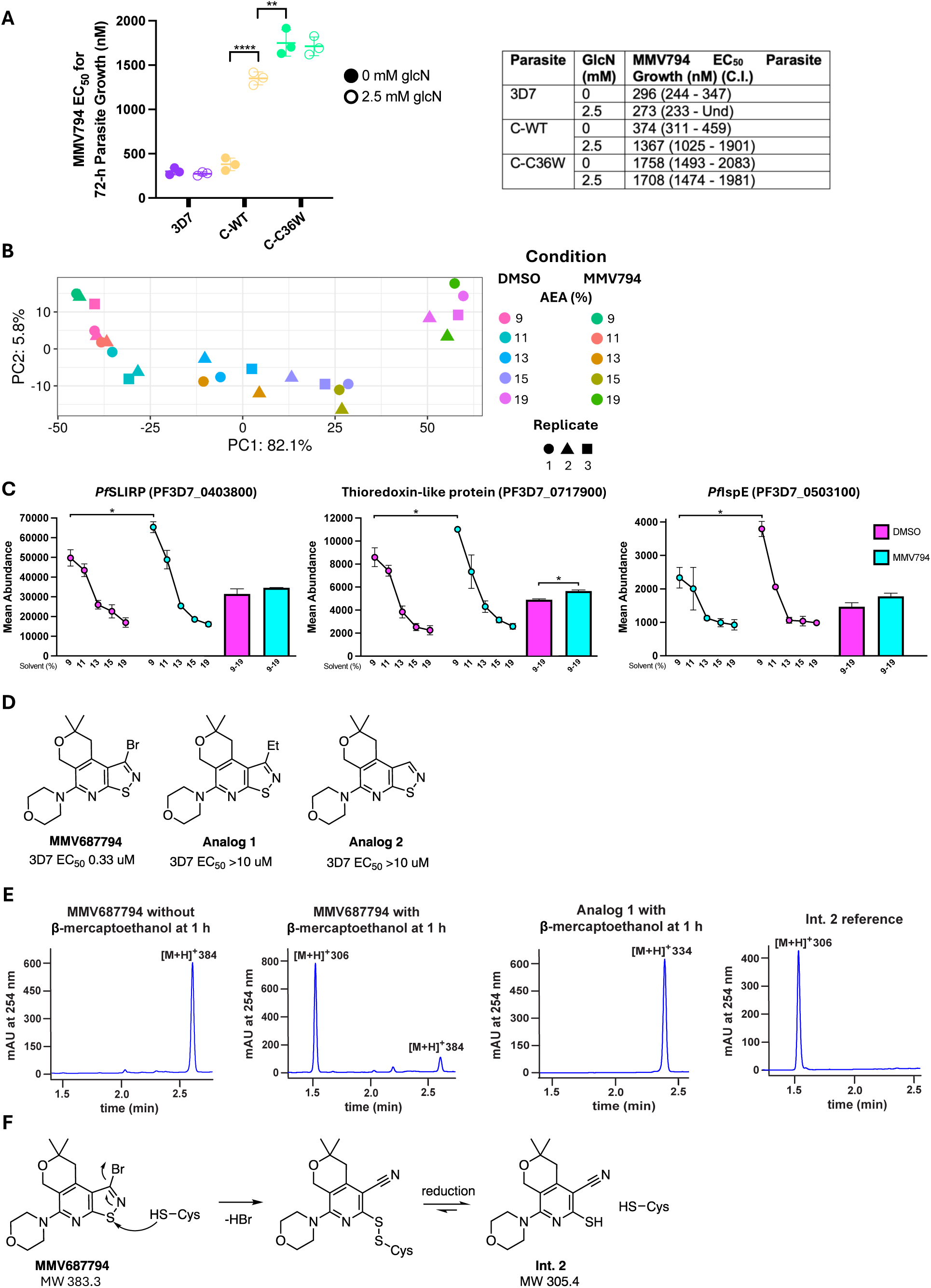
MMV794 engages *Pf*SLIRP, likely through its C36. **A** Mean 72-h growth inhibition EC50s of MMV794 treated 3D7, C-WT and C-C36W parasites ± 2.5 mM GlcN for 120 h. Mean and standard deviation of three biological replicates completed in technical triplicate. No change in MMV794 EC50 was observed in 3D7 +/- GlcN (negative control for GlcN treatment). C-WT + 2.5 mM GlcN parasites gained 4-fold MMV794 resistance, when compared to untreated C-WT. Untreated C-C36W parasites were 5-fold resistant to MMV794, which was unaltered by GlcN treatment. Values were normalised to vehicle control (0.1% DMSO). Error bars represent the mean of three biological replicates, each with three technical replicates. C.I. represents 95% confidence intervals for EC50 values. Statistical analyses were performed on GraphPad Prism 10 with ordinary one-way ANOVA with Dunnett’s multiple comparison test between C-WT + 2.5 mM GlcN and untreated C-WT or C-C36W. ** *p* < 0.01, **** *p* < 0.0001. **B** Solvent-induced protein precipitation of 3D7 parasite lysate treated with either DMSO (1%) or MMV794 (3 µM) exposed to a mixture of acetone/ethanol/acetate (AEA) at 9, 11, 13, 15, and 19%. Principal component analysis of the global proteome profiles demonstrates clear separation in PC2, demonstrating that most changes in the proteome is due to solvent destabilisation. **C** Three parasite proteins, including *Pf*SLIRP, were stabilised by MMV794 when challenged with AEA at varying concentrations. Protein mean abundances were plotted as curves at 9, 11, 13, 15 and 19% AEA concentrations or as bar graphs at 9-19% AEA. Error bars represent the standard deviation of the mean of three (DMSO) or two (MMV794) replicates. Statistical analyses were performed on GraphPad Prism 10 using Welch’s *t*-test between DMSO and MMV794 at all solvent concentrations (9, 11, 13, 15 and 19% AEA) and at solvent gradient (9-19% AEA). * *p* < 0.05. No bar indicates not significant. **D** Structures of MMV687794 and inactive analogs. **E** Reactivity of β-mercaptoethanol with MMV687794 in a PBS buffered solution incubated for 1 h at 37 oC. LCMS analysis shows the degradation of MMV687794 to product Int. 2 in the presence of β-mercaptoethanol, but not with Analog 1. A reference LCMS analysis is shown of Int. 2. **F** Proposed mechanism showing the reversible reactivity of the 3-bromo isothiazole warhead with the *PfSLIRP* Cys36 that proceeds via a disulfide intermediate that subsequently is reduced to give product Int. 2.

To further dissect MMV794-*Pf*SLIRP engagement and investigate other potential MMV794 binding partners, solvent-induced proteome profiling (SPP) was employed. This approach identifies significant protein-ligand interactions/stabilisations by measuring the abundance of soluble proteins stabilised by MMV794, challenged with organic solvents that denature proteins^80–83^. Binding of the compound to its target protein protects it from denaturation, which can then be detected through mass spectrometry. To perform SPP, soluble protein lysates were obtained from segmented 3D7 schizonts, incubated with either 1% DMSO or 3 µM MMV794, and challenged with increasing concentrations (9, 11, 13, 15, 19%) of acetone:ethanol:acetate (AEA). This was followed by centrifugation to collect soluble proteins, which should contain a higher abundance of proteins stabilised by MMV794. Soluble protein fractions were subjected to global proteome (LC-MS/MS) analysis, and mean protein abundance was calculated using data-independent acquisition mass spectrometry analysis (DIA-MS)^84^. This analysis identified 5,069 proteins, of which 3,785 were *Plasmodium* proteins (Table S7). PCA of protein mean abundance in parasites treated with either DMSO or MMV794 across all AEA concentrations illustrates clear separation in PC1, accounting for 82.1% of the variance (Figure 9B). This establishes that the predominant source of variation in the parasite proteome is driven by solvent-induced denaturation (increasing AEA concentration; Figure 9B). MMV794 demonstrated significant target engagement (*p* < 0.05) with three *Plasmodium* proteins having increased mean abundances compared to DMSO, at specific AEA concentrations or across the AEA concentration gradient (Figure 9C). *Pf*SLIRP was notably stabilised by MMV794 at 9% AEA (*p* = 0.0155), demonstrating that MMV794 indeed engages with *Pf*SLIRP (Figure 9C). MMV794 also engaged two proteins, including the 4-diphosphocytidyl-2C-methyl-D-erythritol kinase^85^ (*Pf*IspE; PF3D7_0503100) at 9% AEA (*p* = 0.0103), and an uncharacterised thioredoxin-like protein (PF3D7_0717900) at 9% AEA (*p* = 0.0356) and across the AEA concentration gradient (*p* = 0.0162).

Finally, we sought to interrogate the mechanism by which MMV794 interacts with *Pf*SLIRP. MMV794 belongs to an isothiazolinone-related compound class, which has been shown to covalently modify cysteine residues in a monoacylglycerol lipase^86^, an enzyme from the same hydrolase family as *Pf*SLIRP. Hence, to investigate whether MMV794 may act via a similar mechanism in *Pf*SLIRP, we assessed the intrinsic reactivity of MMV794 with thiol groups using β-mercaptoethanol as a cysteine mimic. MMV794 was incubated ± excess β-mercaptoethanol at 37 °C for 1 h, and the reaction products were analysed by LC-MS. In the presence of β-mercaptoethanol, a shift in mass corresponding to thiol adduct product (Int. 2) was detected (Figure 9E), indicating that MMV794 contains an electrophilic warhead capable of reacting with free thiol groups on cysteines that proceeds via a covalent adduct to afford Int. 2 (Figure 9F). Analog 1 and 2 that do not contain a bromo leaving group are inactive against *P. falciparum* 3D7 parasites (Figure 9D) consistent with data showing Analog 1 does not covalently react with β-mercaptoethanol (Figure 9E). Overall, these findings support a model in which MMV794 targets *Pf*SLIRP by covalently interacting with its C36 residue.

## Discussion

Here, we demonstrate that MMV794 is a schizont-specific inhibitor that targets *Pf*SLIRP, a membrane-associated lipid-interacting α/β hydrolase possibly unique to blood-stage Apicomplexa. This results in defective invasion of RBCs both before and after internalisation and by disrupting PV sealing during invasion. Resistance to MMV794 was driven by mutations (C36W and Δ5’UTR) and knockdown of *Pf*SLIRP. Localisation and lipidomics studies confirmed that *Pf*SLIRP is at the rhoptry surface and has a role in parasite lipid metabolism. Collectively, MMV794 has revealed *Pf*SLIRP as a novel <u>S</u>urface-associated <u>L</u>ipid-Interacting <u>R</u>hoptry <u>P</u>rotein, positioned at the intersection of rhoptry trafficking and invasion-related lipid metabolism.

Our SPP experiment confirmed that MMV794 directly targets *Pf*SLIRP, and the β-mercaptoethanol reactivity assay indicates that this interaction could occur through covalent attack of the 3-bromoisothiazole warhead of the compound at C36^86^. This covalent engagement appears highly deleterious, as resistant parasites have evolved two distinct strategies to avoid it. One resistance pathway is exemplified by the C36W variant (Pop C_C6), in which the small nucleophilic cysteine is replaced by a bulky indole-containing tryptophan located ∼19 Å from the catalytic serine (Figure S18; Video S3). This distance argues against a role for C36 in catalytic regulation^87^. Instead, substitution with tryptophan would abolish the reactive thiol required for covalent engagement by the electrophilic warhead and remodel the local surface topology^88,89^, thereby preventing covalent modification and protecting the protein from MMV794 binding. Interestingly, C36W parasites exhibit reduced *Pf*SLIRP abundance in our proteomics dataset, suggesting that while C36W prevents covalent inactivation, it also compromises protein stability or translation efficiency, resulting in lower steady-state levels. This was consistent with our observations that *glmS* knockdown of the C36W transgenic line had no impact on its MMV794-resistance phenotype. A second, transcriptional resistance strategy was observed in two other resistant populations (Pop B_F10 and Pop B_F6), in which a ∼2600 bp upstream deletion reduced *Pf*SLIRP expression. This regulatory lesion likely disrupts promoter or enhancer elements, decreasing transcription and thereby limiting the pool of *Pf*SLIRP available for interaction. The *glmS* knockdown of the wild-type line phenocopied this resistance, reinforcing that reduction of *Pf*SLIRP abundance alone is sufficient to mitigate MMV794-driven toxicity. In this context, the compound becomes ineffective because there is little substrate to modify. Together, these findings suggest that the MoA of MMV794 may arise from drug-induced destabilisation or misfolding of *Pf*SLIRP following covalent adduction at C36. Thus, the parasite can escape MMV794 either by structurally preventing drug binding (C36W) or by limiting the amount of vulnerable protein (Δ5’UTR). Alternatively, MMV794 may act as a prodrug activated by *Pf*SLIRP, such that reduced *Pf*SLIRP expression would decrease MMV794 activation, hence the MMV794 resistance observed in the transgenic wild-type parasites when *Pf*SLIRP is knocked down. A similar prodrug-activating mechanism has been documented for *Pf*PARE, an esterase in the α/β hydrolase family that converts pepstatin esters into their active form^49^. While our data are consistent with *Pf*SLIRP C36 acting as a reactive nucleophile for MMV794, we cannot exclude the possibility that C36 may serve in a catalytic prodrug activation process rather than being a direct mechanistic target. Other targets stabilised by MMV794 in the SPP study included a functionally uncharacterised thioredoxin-like protein (PF3D7_0717900) and *Pf*IspE (PF3D7_0503100)^85,90,91^, an important enzyme for isoprenoid synthesis in the apicoplast and is thus highly unlikely to be involved in invasion. Furthermore, we could not find additional evidence for the involvement of these proteins in the MoA of MMV794 as they were not mutated in the resistance selection.

*Pf*SLIRP knockdown (87%) did not affect blood-stage parasite growth rate or lipid profile. As *Pf*SLIRP is refractory to piggyBac mutagenesis^23^, it may remain functional when knocked down to low expression levels, similar to other essential proteins such as plasmepsin V^92,93^, caseinolytic-protease chaperone^94,95^ and cdc2-related protein kinase 4^96^. The low *Pf*SLIRP transcript counts across the blood-stage^64–67^, proteomics expression unique to schizonts^97^ and weak western blot detection support this. The essentiality of *Pf*SLIRP for blood-stage parasite growth may be confirmed using an inducible knockout system such as *loxP*/DiCre^98^.

While *Pf*SLIRP knockdown did not affect the parasite lipidome, mutation to the highly conserved C36 residue significantly altered the lipidome across two independent analyses, with concordant changes seen in PC, LPC and FFA in particular. The lipidomic analyses results were consistent with the predicted function of *Pf*SLIRP as a lipid-interacting α/β hydrolase, supporting its proposed role as a (lyso)phospholipase as reflected by reduced FFA in C36W mutant parasites and increased phospholipid substrates (putatively here PC and LPC) required for FFA generation^99–101^. However, pinpointing the exact perturbation caused by the C36W mutation is challenging as *P. falciparum* has multiple overlapping mechanisms that produce these abundant lipids (PC, PE, PG, DAG). For example, *P. falciparum* can generate the major phospholipids PS, PE and PC *de novo* via two, three, and four routes in normal conditions, respectively^102^. Future experiments may address this by through protein lipid overlay assays with a recombinant form of wild-type or C36W *Pf*SLIRP to directly assess its lipid-binding capacity.

Proteomic profiling of MMV794-resistant parasites also revealed altered proteomes, including proteins linked to invasion such as *Pf*RAP1, indicating diminished rhoptry vesicular trafficking. The comparable growth of MMV794-resistant parasites to wild-type parasites in culture supports previous findings^76^ that while *Pf*RAP1 is involved in rhoptry biogenesis by trafficking other rhoptry proteins like *Pf*RAP2, its low abundance is likely sufficient to support parasite invasion. *Pf*FLP, implicated in RBC gamete egress in *P. berghei*^79^ and microneme vesicular trafficking *T. gondii*^103^, was also notably downregulated in resistant parasites. Concurrent with the upregulation of Falstatin, a cysteine protease inhibitor proposed to enable selective cleavage at the invasion site for efficient RBC invasion^78^, these changes signify possible compensatory adaptations in parasites to preserve controlled invasion dynamics amid disrupted invasion/rhoptry (*Pf*SLIRP) function. In MMV794-resistant parasites, perturbed parasite lipid metabolism is also evidenced by the downregulation of CTP:phosphocholine cytidylyltransferase (*Pf*CCT), the rate-limiting enzyme in the synthesis of phosphatidylcholine, the most abundant (48%) phospholipid in iRBC membrane^104–106^. Reductions in *Pf*EVP1 were also observed in resistant parasites, a protein required for vesicular lipid import at the iRBC membrane^77^. These changes at the protein level align with our lipidomic findings that MMV794-resistant parasites exhibit changes in lipid metabolism pathways. Mutations to *Pf*SLIRP likely alter its function and prompt compensatory changes in other lipid-interacting proteins, ultimately reshaping the cellular lipid composition. Disruption of *Pf*SLIRP and invasion and secretory trafficking pathways may have created downstream consequences for virulence factor export by affecting ER–Golgi trafficking, increasing misfolded or misrouted STEVOR^107^, RIFIN^108,109^ and *Pf*EMP1^110^, and selecting for parasites that reduce STEVOR and A-RIFIN while maintaining *Pf*EMP1 variants^111^ better suited to stress. The accompanying reduction in *Pf*EMP1 export is consistent with decreased host TRiC/CCT abundance^112^, while accumulation of disorganised cargo likely triggers a proteostasis response, including *Pf*PA28 upregulation to enhance proteasome activity.

Lattice light-sheet microscopy revealed that MMV794 treatment of schizonts resulted in fewer merozoites initiating RBC internalisation during invasion, and those that do mostly fail to complete internalisation. This defect was unlikely due to total rhoptry secretion failure, as RON3 is released, but rather reflects incomplete or misassembled rhoptries that are sufficient for stable merozoite-RBC attachment but insufficient for invasion pore sealing. Similar phenotypes have been observed in parasites with protein expression knockdown of moving junction protein AMA1^40^ or subtilisin-like protease 2 (SUB2)^113^, which is responsible for shedding merozoite surface proteins such as MSP1 and AMA1 during invasion. Like these mutants, MMV794-treated parasites can form a moving junction and initiate internalisation but cannot seal the pore, resulting in regression. RBC internalisation by *P. falciparum* involves coordinated mechanical and receptor interactions coupled with lipid remodelling at the host-parasite interface. The moving junction not only anchors the merozoite but also serves as a conduit for transferring proteins and lipids from apical organelles^30,32,114^. Although the PVM largely derives from the RBC membrane^32^, phospholipids injected at the merozoite apex appear in both the plasma membrane and PVM post-invasion^115^. Similar multilamellar lipid aggregates are expelled by *P. knowlesi* merozoites^116–118^, likely aiding membrane curvature and fluidity. These lipid dynamics may support signalling events such as trafficking heterotrimeric G proteins in lipid rafts to the PVM^119^, which affects host membrane deformability^120–123^. Furthermore, cholesterol has also been shown to play a critical role in resealing the RBC membrane post-entry, likely by facilitating RBC membrane curvature during merozoite invasion^32^. Hence, *Pf*SLIRP inhibition may alter the lipid composition of the nascent PVM, making it mechanistically difficult for the merozoite to wrap the RBC membrane around itself and thereby reducing invasion efficiency. When MMV794-treated parasites do manage to invade, they subsequently encounter difficulties resealing the RBC membrane, a process that presumably also requires optimal membrane curvature and lipid composition.

We propose that *Pf*SLIRP integrates vesicular trafficking and lipid-driven rhoptry biogenesis during schizogony, organising invasion and lipid-modifying proteins within rhoptries to ensure moving junction competency (Figure 10). Although MMV794-treated parasites form moving junctions and secrete rhoptry contents, they may fail to deliver the complete rhoptry payload required to assemble a functional junctional complex capable of effective lipid remodelling and internalisation. Future localisation of fluorescently-tagged *Pf*SLIRP, alongside co-immunoprecipitation of its interacting partners, will help define its precise at the rhoptries. As resistance to existing antimalarial drugs intensifies, uncovering novel vulnerabilities in parasite biology from mechanistic studies offers new avenues for therapeutic intervention.

**Figure 10.**
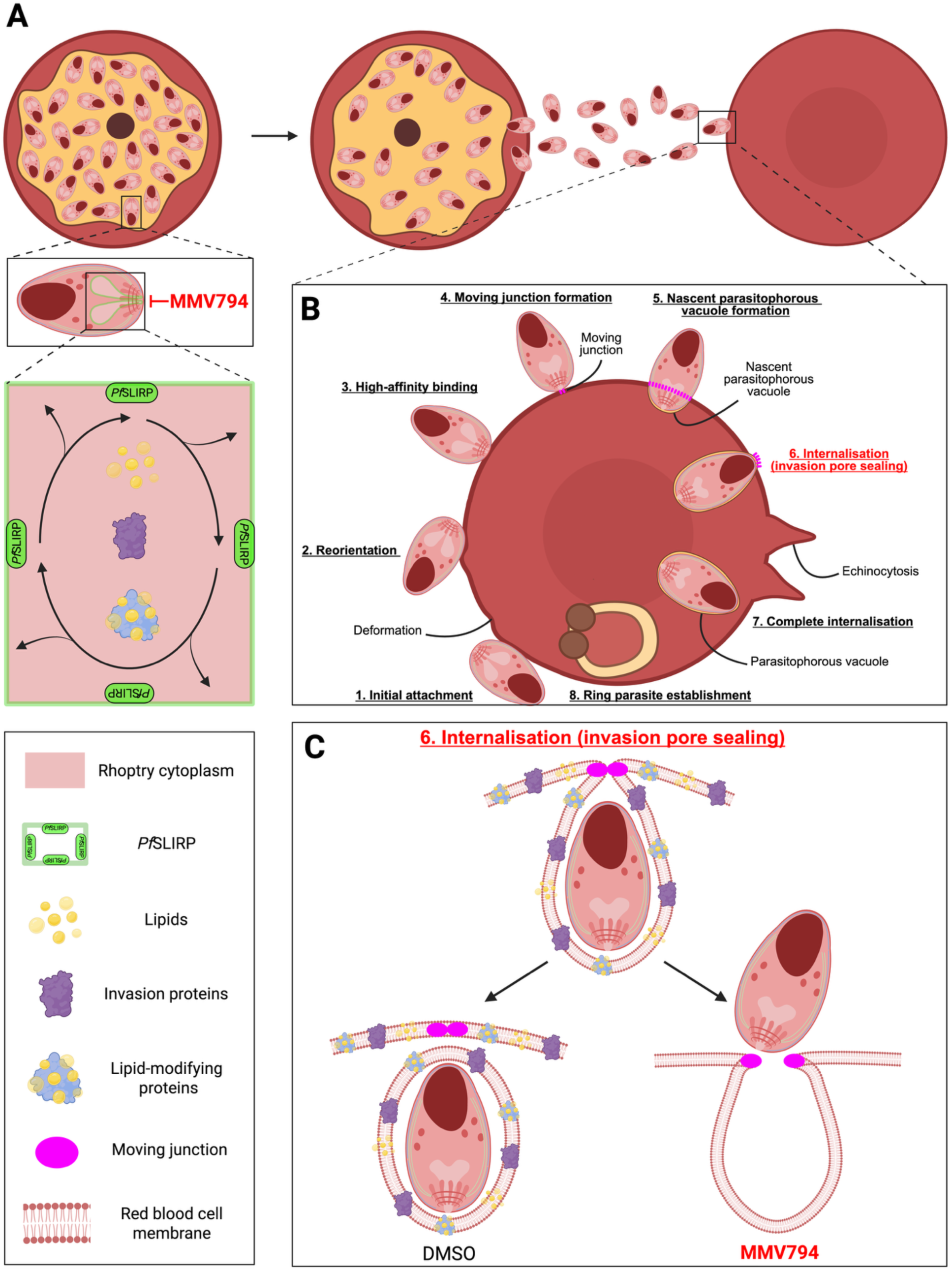
Proposed function of *Pf*SLIRP in blood-stage *P. falciparum* invasion. **A** MMV794 inhibits *Pf*SLIRP found at the rhoptry surface, which contributes to rhoptry function by trafficking of lipids, invasion-related and lipid-modifying proteins during schizogony. **B** Inhibition of *Pf*SLIRP by MMV794 blocks invasion downstream by preventing internalisation of the merozoite within the red blood cell (RBC). **C** MMV794 treatment permits the formation of a moving junction sufficient for merozoite entry into the RBC, but this junctional complex is insufficient for sealing the invasion pore, likely due to incomplete delivery of rhoptry contents needed for RBC membrane remodelling.

## Methods

### Chemistry Methods

The methods for synthesis of MMV687794 and its analogs are outlined in the Supplementary Information.

### *In vitro* culturing of *P. falciparum*

Parasite culturing^124^, sorbitol synchronisation^125^, Percoll purification^126^, and magnet purification^127^ were performed with slight modifications, and are fully detailed in the Supplementary Information.

### Colorimetric and fluorescent plate-based assays

The 72-h lactate dehydrogenase growth assay and the Nanoluciferase 4-h egress and invasion, trophozoite growth, purified merozoite invasion assays were performed as previously described^18^. The Nanoluciferase ring growth assay was adapted from the early ring inhibition assay^18^ and was performed on mixed-stage ring parasites (0-12 hpi).

### Widefield live-cell imaging

Parasites were imaged similarly as described^14^. Two days prior to imaging, 3D7 ring-stage parasite cultures (≥5 % rings) were sorbitol synchronised, resuspended to 2% HCT, delayed at RT for 7 hours and returned to incubation at 37 °C for 16 h until next day. Then, parasite culture medium was replenished containing 30 nM (ML10) (MRT-0207065, BEI Resources) and parasite cultures were again delayed at RT for 7 hours before returning to incubation at 37 °C for 16 h until next day. On the day of imaging, the chambers of an 8-well Nunc™ Lab-Tek™ II chambered cover glass (Thermo Scientific) were rinsed 3× with 1 mL warm cRPMI. The ML10-treated synchronous 3D7 schizont cultures were washed (500 g/5-10 mins) in cRPMI and diluted to 0.1% HCT. Then, 200 µL of the 0.1% HCT culture was dispensed into the wells of the chambered cover glass. Parasites were allowed to settle onto the coverglass for 30 mins before imaging on a Zeiss Cell Observer Z1 inverted widefield fluorescent microscope equipped with a Plan-Apochromat 100×/1.40 Oil DIC objective and a humidified gas chamber (1% O_2_, 5% CO_2_ and 94% N_2_) at 37 °C and an AxioCam 702 Mono camera. Egress and invasion events were captured at 4 frames per second for 20 mins and cell behaviour was examined manually using Fiji software.

### Scoring of invasion events

To examine the effects of MMV794 on merozoite invasion, parameters such as the number of egresses, RBC contact, deformation and internalisation were analysed manually (Table S6). Statistical analyses were performed between vehicle control (0.1% DMSO)- and compound-treated parasites from live cell videos on GraphPad Prism 9 using Welch’s *t*-test.

### Lattice light-sheet live-cell imaging

RON3-mNG-AMA1-mSc parasites were prepared for imaging as above (widefield live-cell imaging) with an additional Percoll purification step on the day of imaging to obtain ≥80% schizont parasite pellets. For RBC membrane staining, RBCs were washed once in iRPMI in a 1.5 mL microcentrifuge tube (3,000 rpm/2 mins) and resuspended at 0.5% HCT in iRPMI stained with 1.5 µM Di-4-ANEPPDHQ (Thermo Scientific) for 1 h at 37 °C. Then, stained RBCs were washed (3,000 rpm/2 mins) 3× in imaging medium (cRPMI with 10 µM Trolox (Santa Cruz Biotechnology)). For imaging, the parasite pellet was mixed with the stained RBCs and loaded onto an 8-well chambered coverslip (µ-slide 8-well high glass bottom; Ibidi) to 0.1% HCT and ∼60% parasitaemia, containing either 3 µM MMV794 or 0.03% DMSO, to a final volume of 200 µL in imaging medium.

Parasite cultures were allowed to settle for 30 mins before imaging on the Zeiss Lattice Lightsheet 7. Samples were illuminated with laser light at 488 nm, 560 nm, or 640 nm, via a 13.3×/0.44 illumination objective, generating a light sheet with dimensions of 30 µm length and 1 µm thickness. Emitted fluorescence was collected through a 44.83×/1.0 detection objective and directed to a dual camera system through a 565 nm beamsplitter and a quad-notch 405/488/561/640 nm multi-band stop filter before camera 1 and a 525-550 nm bandpass filter before camera 2. The 488 nm laser was used to excite mNeonGreen (RON3) and Di-4-ANEPPDHQ, the 561 nm laser for mScarlet (AMA1). 290 × 200 µm × 30 µm volumes were scanned with 0.4 µm intervals at 2 ms exposure time (per frame) for 30 mins. Acquired data was deskewed and deconvolved as previously. Deskewed and deconvolved data were processed using Imaris 10.2 software.

### *In vitro* drug resistance selection (on and off drug cycling)

Resistance selection was performed as described^80^ with 3 µM MMV794 and 0.03% DMSO (vehicle control) for three cycles.

### Generation of p1.2-*Pf*SLIRP-HA-*glmS* constructs for CRISPR-Cas9 transfection

Methods for generation of the p1.2-*Pf*SLIRP-HA-*glmS* plasmid used for CRISPR-Cas9 transfection^55^ into 3D7 parasites were fully described in the Supplementary Information.

### Saponin lysis

Parasite pellets were obtained from saponin lysis^128^ with washes performed in 1× PBS+Protease Inhibitor Cocktail Tablet (PI; Roche) as detailed in the Supplementary Information.

### gDNA extraction

gDNA was extracted from parasite pellets (from saponin lysis) as previously^42^, fully described in Supplementary Information.

### Whole genome sequencing and variants analyses

MMV794-resistant parasites and parental 3D7 parasites were whole-genome sequenced, and raw sequencing data was converted to DNA nucleotides, demultiplexed and identified for single nucleotide polymorphism and structural variants^42,129,130^. The genomic sequencing data is available from the European Nucleotide Archive; accession number PRJEB89677).

### Sanger sequencing

Plasmid DNA (100 ng/µL) or PCR product (10 ng/µL) was sent for Sanger sequencing at Micromon Genomics (Monash University, Clayton) with primers listed in Tables S4 and S5. Sequences received were examined by Sequencher® DNA sequence analysis software (or SnapGene V5.3.3).

### Bioinformatics

Basic local alignment search tool (BLAST) searches^45^ of *Pf*SLIRP (PF3D7_0403800) were performed using predicted protein sequence from PlasmoDB^131^ against other Apicomplexan parasites and *Plasmodium spp.* to identify potential orthologous genes. All multiple sequence alignments were performed using Clustal Omega^132^ to identify conserved regions of *Pf*SLIRP, which were mapped onto the predicted 3D structure from AlphaFold 3^21,22,133^.

### Western blotting sample preparation (*Pf*SLIRP-HA validation, *Pf*SLIRP parasite stage expression, *Pf*SLIRP knockdown, carbonate extraction)

To validate the HA tagging of *Pf*SLIRP, mixed-stage (predominantly trophozoites and schizonts) 10 mL parasite cultures at 4% HCT and ≥ 5% parasitaemia were harvested via saponin lysis.

To examine stage-specific expression of *Pf*SLIRP, individual 30 mL ring (0-12 hpi), early trophozoite (18-24 hpi), late trophozoite (30-36 hpi), and schizonts (40-46 hpi) cultures were prepared. Contamination of asynchronous parasites was minimised by 2× sorbitol treatment of cultures. Post-sorbitol treatment, ring cultures were harvested immediately, early trophozoites and late trophozoites were harvested upon reaching the desired stage, schizonts were treated with 30 nM ML10 and Percoll purified before harvest via saponin lysis.

To examine the level of *Pf*SLIRP knockdown, early-trophozoite (18-24 hpi) C-WT and C-C36W parasites at 1% parasitaemia were treated with 2.5 mM glucosamine for 72 h until they reached the schizont stage, which were harvested via saponin lysis.

The carbonate extraction protocol was adapted from Grüring *et al.*^134^ and fully described in the Supplementary Information.

Densitometry analyses were conducted using the Image Studio™ v.1.0 programme and adjusted to loading control (HSP70-1):

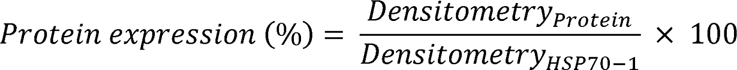

### Indirect immunofluorescence assays (IFAs)

IFAs were prepared either through iRBC smears or cultures of Percoll-purified iRBCs. Images were captured using a Zeiss Axio Observer Z1 inverted widefield microscope with a Plan-Apochromat 100×/1.40 Oil DIC objective and the super-resolution radial fluctuations (SRRF) algorithm^58^ was used to capture super-resolution images. All images were processed using Fiji software.

IFAs performed on iRBC smears using thin smears made on glass slides (acetone/methanol fixation) or iRBCs cultures settled onto poly-L-lysine- (Sigma-Aldrich) coated 10 mm round coverslips (paraformaldehyde/glutaraldehyde fixation) were prepared as described^135^, fully detailed in the Supplementary Information. Fixed cells were blocked in 3% BSA/0.02% TX100 in PBS for 1 hour at RT. To minimise non-specific binding, 1 mL primary antibodies in 3% BSA/0.02% TX100 were incubated with RBCs at 20% HCT at RT for 90 mins. The cross-adsorbed primary antibodies were then added to the cells overnight at 4 °C. Cells were washed with 0.02% TX100 in PBS to remove unbound antibodies and probed with Alexa Fluor secondary antibodies as above. Washes were performed as before. Coverslips were mounted on slides with 2-5 µL of VECTASHIELD® Antifade Mounting Medium with DAPI (Vector Laboratories).

### Ultrastructure expansion microscopy (U-ExM) IFA

U-ExM^60–62^ was performed with slight modifications, protocol is fully detailed in Supplementary Information.

For staining, the gels were washed 2× for ≥ 10 mins with 1× PBS on a platform rocker with 1 mL 2% BSA (Merck Life Science) at RT for ≥ 20 mins. The gels were then probed with 1 mL of primary antibodies in 2% BSA at 4 °C overnight in the dark on a platform rocker, washed 3× for ≥ 5 mins with 1× PBS at RT on a platform rocker, then probed with 1 mL of secondary antibodies in 1× PBS at RT for 2-3 h in the dark on a platform rocker. The gels were washed 2× for ≥ 5 mins with 1× PBS at RT in the dark on a platform rocker. Then, the gels were transferred into a 90 mm petri dish and washed 2× for ≥ 10 mins with Mili-Q water at RT in the dark on a platform rocker for expansion.

For imaging, #1.5 25 mm round glass coverslips (Fisher Biotec) were coated with 0.1 mg/mL poly-D-lysine for ≥ 1 h at RT, and gels were mounted onto the coverslips in an Attofluor™ cell chamber (Thermo Scientific). The chamber was then placed onto a Piezo stage and imaged on the Zeiss laser scanning microscope 980 with Airyscan 2 using the Plan-Apochromat 63×/1.40 Oil DIC objective. The following settings were used to acquire z-stacks with Airyscan 2: full Z-stack per Track, optimal sampling (0.148 µm), bidirectional frame scanning, 35 × 35 × 150 nm voxel size, 2 sampling rate, 0.70 µs pixel dwell time, 850 V detector gain, and 0.5% (405 nm), 1% (488 nm), 1% (561 nm) and 0.3% (639 nm) laser power. Images were processed using Imaris 10.2.

### Time course analysis of parasite growth post-*Pf*SLIRP knockdown

C-WT parasites at 4% HCT and 1% parasitaemia were treated ± 2.5 mM GlcN over 264 h (5.5 cell cycles) starting from rings, with media and GlcN replenished daily. Parasite cultures were diluted with fresh RBCs when trophozoite parasitaemia reached ≥ 5%. Parasites were monitored daily via brightfield microscopy of Giemsa-stained thin blood smears of cultures and cell counting of 1000 cells including RBCs and iRBCs. The parasite growth rate was calculated by dividing the parasitaemia every cycle (48 h) by the parasitaemia of the cycle before. Dilution factors were also used to calculate parasitaemia at timepoints when parasite cultures were diluted. Images were taken on a brightfield microscope Leica DM750 equipped with the digital microscope camera Leica ICC50 HD.

### MMV794 sensitisation assay

3D7, C-WT and C-C36W parasites were treated with ± 2.5 mM GlcN for 48 h (rings to rings) and LDH assays (as described above) with MMV794 were performed on these parasite cultures ± 2.5 mM GlcN. Parasites initially treated with GlcN were continually treated with 2.5 mM GlcN for the duration of the 72-h LDH assay, totalling 120 h of GlcN exposure (*Pf*SLIRP knockdown).

### Baseline parasite metabolomic analysis of C-WT

Methods for analysing baseline metabolome of C-WT parasites compared to 3D7 parasites ± 2.5 mM GlcN are outlined in the Supplementary Information.

### Parasite lipidomics analysis (post-*Pf*SLIRP knockdown)

C-WT and C-C36W parasite cultures were prepared as follows: 30 mL ring-stage parasite cultures at 2% HCT and 2-3% parasitaemia were sorbitol treated, grew to early trophozoites, split into 2× 10 mL cultures each, and treated ± 2.5 mM GlcN for 72 h until parasites reached the schizont stage. During the 72 h-GlcN treatment period, the cultures were also treated with 30 nM ML10 at the 56 h timepoint overnight for 16 h to stall parasites at the late-schizont stage (44-48 hpi) at the 72 h timepoint. Then, these cultures were monitored to ensure schizont fitness via microscopy examination of Giemsa-stained thin blood smears. The schizont cultures were pelleted (500 g/10 mins), washed 1× with 50 mL cRPMI, dispensed into new culture dishes with 50 mL cRPMI and incubated at normal culture conditions for 4 h for invasion. Parasite invasion was validated via monitoring of cultures via microscopy examination of Giemsa-stained thin blood smears. Then, the cultures were pelleted (500 g/10 mins), resuspended in 10 mL cRPMI and quenched in a dry ice-ethanol slurry until they reached 10 °C. Finally, the cultures were pelleted (500 g/10 mins), and the pellets were harvested via 0.15% saponin lysis.

Parasite lipids were extracted by chloroform-methanol (chloroform:methanol:water 1:3:1 (v/v/v))^70^ and fully detailed in the Supplementary Information. Briefly, parasite lipids were extracted from saponin pellets spiked with internal standards (tridecanoic acid and phosphatidylcholine) and SPLASH® LIPIDOMIX® Mass Spec Standard (Avanti), using chloroform:methanol:water 1:3:1 (v/v/v) with periodic sonication. The samples (organic phase) were dried down in a SpeedVac with the drying rate set to high and stored at - 20 °C until subsequent gas or liquid chromatography-mass spectrometry analysis. Samples were analysed in random sequence along with blank samples.

Analysis of total parasite lipids by LCMS are fully described in the Supplementary Information. Briefly, dried-down lipids were reconstituted in methanol and incubated at 30 °C with vigorous vortexing. Total lipids in methanol were injected into an Agilent 1290 infinity/Infinity II LCMS system equipped with ZORBAX Eclipse Plus C18, 2.1 × 100 mm, 1.8 µm columns (Agilent) with Infinity II inline filter, 0.3 µm (Agilent) maintained at 45 °C. Samples were separated by the charge in the gradient of solvent A (water:acetonitrile:isopropanol, 5:3:2, (v/v/v), 1 mM ammonium formate) and solvent B (isopropanol:acetonitrile:water, 90:9:1 (v/v/v), 1 mM ammonium formate). MS (Agilent 6495c triple quadrupole) was operated in for targeted analysis with DMRM (dynamic multiple reaction monitoring) with 650 ms of cycle time. Acquisition DMRM method used was referred to^136^ with a modification for *P. falciparum* lipid species. The resultant LCMS data was subjected to targeted analysis using Mass Hunter Quantification software (Agilent). Each lipid species was quantified using a calibration curve of each representative lipid with known abundance. All solvents used for this LCMS analysis were purchased from Sigma-Aldrich.

### Parasite lipidomics analysis (post-MMV794 treatment)

C-WT and C-C36W parasite cultures were prepared as follows: 30 mL ring-stage parasite cultures at 4% HCT and ≥ 10% parasitaemia were sorbitol treated and stalled at the early schizont stage (40-44 hpi) with overnight (16 h) treatment with 30 nM ML10 at the early trophozoite stage. Magnet purification was performed on the 30 mL cultures to isolate schizonts, which were split equally into 2× 10 mL cultures. The cultures were treated with ± 3 µM MMV794 (10× 72-h parasite growth EC_50_), incubated at normal culture conditions for invasion for 5 h and pelleted (500 g/10 mins). 160 µL of the supernatant was aliquoted for the analysis of MMV794 abundance, while the rest was aspirated. The parasite pellet was quenched in 1 mL ice-cold 1× PBS, pelleted (1,000 g/5 mins) and the supernatant was aspirated.

Parasite lipids were extracted via chloroform-methanol (chloroform:methanol:water 2:6:1 (v/v/v)) as fully described in the Supplementary Information. The samples (organic phase) were dried down for lipidomics in the Savant® DNA110 SpeedVac® concentrator (Thermo Scientific) with the drying rate set to high. Dried-down samples were wrapped in Parafilm and stored at -80 °C until LCMS analysis.

Analyses of MMV794 and total parasite lipids by LCMS are fully described in the Supplementary Information. Briefly, dried-down samples were reconstituted in butanol:methanol:water (45:45:10) and sonicated at RT for 30 mins and centrifugated (21,000 g/10 mins) to remove cell debris. A pooled biological quality control was included to identify metabolites and to monitor downstream sample stability. Samples were injected into a Dionex UltiMate™ rapid separation liquid chromatography system (Thermo Scientific) equipped with Supelco® Ascentis Express® C8 100 × 2.1 mm, 2.7 µm reversed-phase column (Merck) with a Luna C8 2 × 2 mm, 5 µm guard column (Phenomenex) maintained at 40 °C. Samples were separated by the change in gradient of solvent A (water:isopropanol, 60:40. (v/v), 8 mM ammonium formate) and solvent B (isopropanol:water, 98:2, (v/v) 8 mM ammonium formate and 2 mM formic acid (FA)). MS was operated in full scan mode with positive and negative polarity switching at 70,000 resolution with detection range of 140-1,300 *m/z* to collect positive and negative ion mode data.

The LCMS data obtained was subjected to untargeted processing^137^ using open-source software identification and evaluation of metabolomics data from LCMS (IDEOM)^73^, fully described in the Supplementary Information.

### Untargeted comparative proteome profiling

Preparation of parasite cell lysates used for MS/MS analysis were performed from an adapted protocol of Siddiqui *et al.*^97^ and are clearly detailed in the Supplementary Information.

### Assessment of MMV687794 warhead reactivity using β-mercaptoethanol

The intrinsic warhead reactivity with β-mercaptoethanol was assessed following an adaptation of a previously described procedure^138^, outlined in the Supplementary Information.

### Solvent proteome profiling

3D7 parasite lysates were prepared as follows: 30 mL ring-stage parasite cultures at 4% HCT and ≥ 10% parasitaemia were sorbitol treated and stalled at the early schizont stage (40-44 hpi) with overnight (16 h) treatment with 30 nM ML10 at the early trophozoite stage. The cultures were washed 1× with 30 mL cRPMI and pelleted (500 g/10 mins). Parasite cell lysate used for LC-MS/MS analysis were prepared as described^83^, fully detailed in the Supplementary Information.

### LC-MS/MS analysis with Orbitrap Astral using LFQ-DIA

Samples were analysed as described^83^, detailed in the Supplementary Information

### Analysis of SPP LC-MS/MS data

The proteomics data was processed as described^83^, detailed in the Supplementary Information. Proteins were considered to show significant stabilisation only if they had a *p* value < 0.05 and a log2 fold change ≥ 1.2 between control and experimental groups.

## Acknowledgements

We thank Lifeblood Biological Resources Australia and the Australian Red Cross Blood Service for providing the human red blood cells. We thank BEI Resources, NIAID, NIH and LifeArc (Stevenage, UK) for supplying ML10 (MRT-0207065), NR-56525. We thank Molly P. Schneider for her assistance in setting up 96-well plates for cloning selection of parasites by limiting dilution. We thank Emma T. Gill and Lachlan W. Whitehead for assistance in 3D-printing of coverslip mould used for U-ExM imaging. We thank Thomas C. McLean and Stephen W. Scally for their expertise on structural biology. This research was supported by the Australian Society for Parasitology Researcher Exchange, Travel and Training Funding Scheme, and the Burnet Institute InnoVate program. This research was supported by the Commonwealth through Australian Government Research Training Program Scholarships to Dawson B. Ling and Madeline G. Dans. This research was supported by the Victorian State Government Operational Infrastructure Support Grant (Institutional grant) and Australian Government NHMRC Independent Research Institutes Infrastructure Support Scheme. This research was supported by grants from Agence Nationale de la Recherche, France, (Project ApicoLipiAdapt ANR-21-CE44-0010; Project Apicolipidtraffic ANR-23-CE15-0009-01; Project OIL ANR-24-CE15-2171-02; Project Plasmohost ANR-25-CE11-3606), The Fondation pour la Recherche Médicale (FRM EQU202103012700), Laboratoire d’Excellence Parafrap, France (ANR-11-LABX-0024), Centre national de la recherche scientifique, France (International Associated Laboratory-International Research Program Apicolipid project), the Université Grenoble Alpes (IDEX ISP Apicolipid) and Région Auvergne Rhone-Alpes, France, for the lipidomics analyses platform (IRICE Project GEMELI), Centre Franco-Indien pour la Promotion de la Recherche Avancée (CEFIPRA; Collaborative Research Program Grant (Project 6003-1), IARDP grant (#2023-0414)) to Cyrille Y. Botté. This research was supported by grants from the National Health and Medical Research Council (NHMRC), Australia, Investigator Grants 1194535, 1176955, 1177431 to Alan F. Cowman, Ideas Grant 2012271 to Kelly L. Rogers, Synergy Grant 1185354 to Paul R. Gilson.

## Author Information

### Contributions

Study design and planning: D.B.L., G.S., C.A.B., Y.Y.B., M.G.D., W.N., B.E.S., C.Y.B. Performed experiments and/or data analysis: D.B.L., G.S., C.A.B., Y.Y.B., M.G.D., B.K., G.E.W., Z.R., S.M., M.C. Provided funding: A.E.B., B.E.S., T.F.dK-W., C.Y.B., D.J.C., K.L.R., A.F.C., B.S.C., P.R.G. Provided supervision: C.A.M. (M.C.), D.J.C. (M.C.), T.F.dK-W. (M.G.D.), H.E.B. (D.B.L.), P.R.G. (D.B.L.). Manuscript writing: D.B.L., H.E.B., and P.R.G. with contributions from other authors.

