## Supplementary Information for "MMV687794 blocks *Plasmodium falciparum* invasion of red blood cells by targeting a Surface-associated Lipid-Interacting Rhoptry Protein, *Pf*SLIRP"

### Supplementary Figures

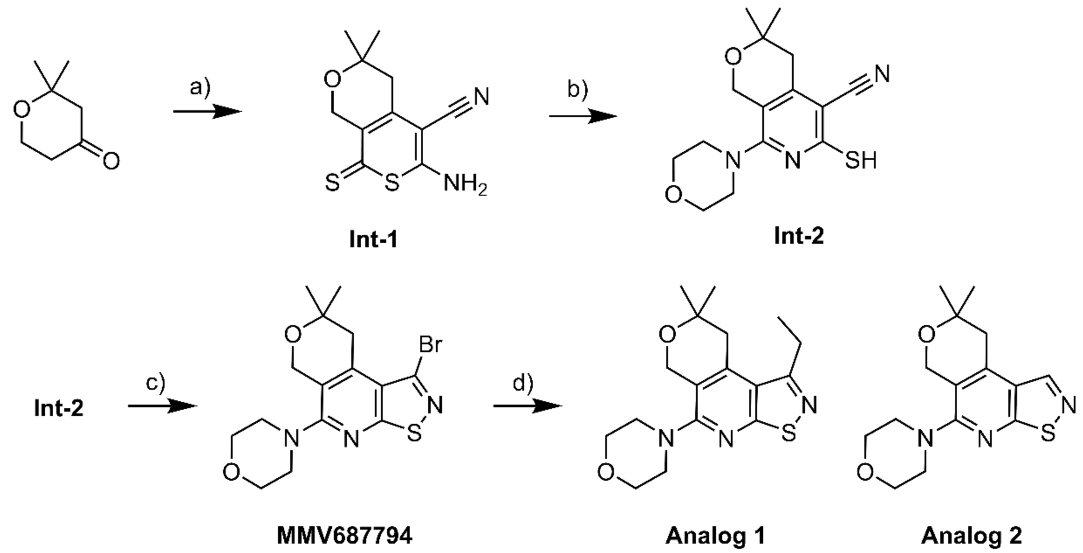


**Figure S1. Synthetic route to access MMV687794 and analogs 1 and 2.** Reagents and conditions: (a) $CS_2$, malononitrile, MeOH, triethylamine, 20 °C; (b) morpholine, ethanol, 78 °C; (c) bromine, $CHCl_3$, 60 °C; (d) diethyl zinc, $Pd(PPh_3)_2Cl_2$, dry THF, 66 °C.

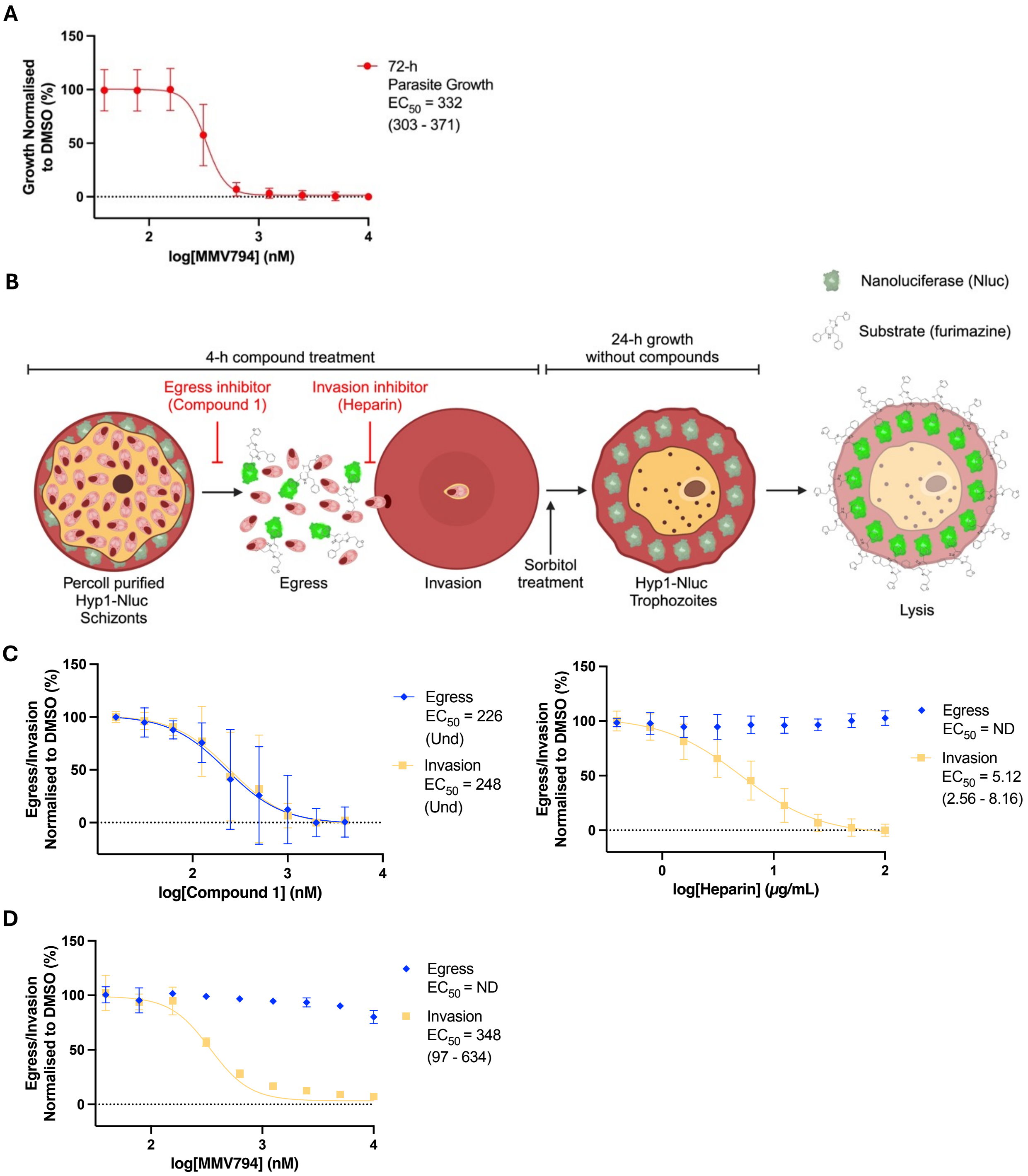
A
Growth Normalised to DMSO (%)
150
100
50
0
2
3
4
log[MMV794] (nM)
72-h
Parasite Growth
$EC_{50}$ = 332
(303 - 371)
B
Nanoluciferase (Nluc)
Substrate (furimazine)
4-h compound treatment
24-h growth
without compounds
Egress inhibitor
(Compound 1)
Invasion inhibitor
(Heparin)
Percoll purified
Hyp1-Nluc
Schizonts
Egress
Invasion
Sorbitol
treatment
Hyp1-Nluc
Trophozoites
Lysis
C
Egress/Invasion
Normalised to DMSO (%)
150
100
50
0
2
3
4
log[Compound 1] (nM)
Egress
$EC_{50}$ = 226
(Und)
Invasion
$EC_{50}$ = 248
(Und)
Egress/Invasion
Normalised to DMSO (%)
150
100
50
0
0
1
2
log[Heparin] (μg/mL)
Egress
$EC_{50}$ = ND
Invasion
$EC_{50}$ = 5.12
(2.56 - 8.16)
D
Egress/Invasion
Normalised to DMSO (%)
150
100
50
0
2
3
4
log[MMV794] (nM)
Egress
$EC_{50}$ = ND
Invasion
$EC_{50}$ = 348
(97 - 634)

**Figure S2. MMV794 inhibits invasion when added to schizonts. A** Dose response of 72-h 3D7 parasite growth for MMV794. **B** Schematic of 4-h egress and invasion assay. Hyp1-Nluc schizonts were allowed to egress and invade in the presence of compounds for 4 h. Supernatant was removed after the 4-h egress and invasion window. Compounds and residual schizonts were removed by sorbitol treatment and washes, and successfully invaded parasites were grown for 24 h, lysed at the trophozoite stage. Furimazine was added to supernatant (post 4-h window) and parasite lysate to generate bioluminescence as a measure of egress and invasion, respectively. Figure created using BioRender.com. **C** Dose response curves of egress and invasion for control compounds, compound 1 (egress inhibitor) and heparin (invasion inhibitor) were as expected. **D** MMV794 inhibits invasion but not egress. C.I. represents 95% confidence intervals for $EC_{50}$ values. Error bars represent the standard deviation of the mean of three biological replicates, each with two or three technical replicates.

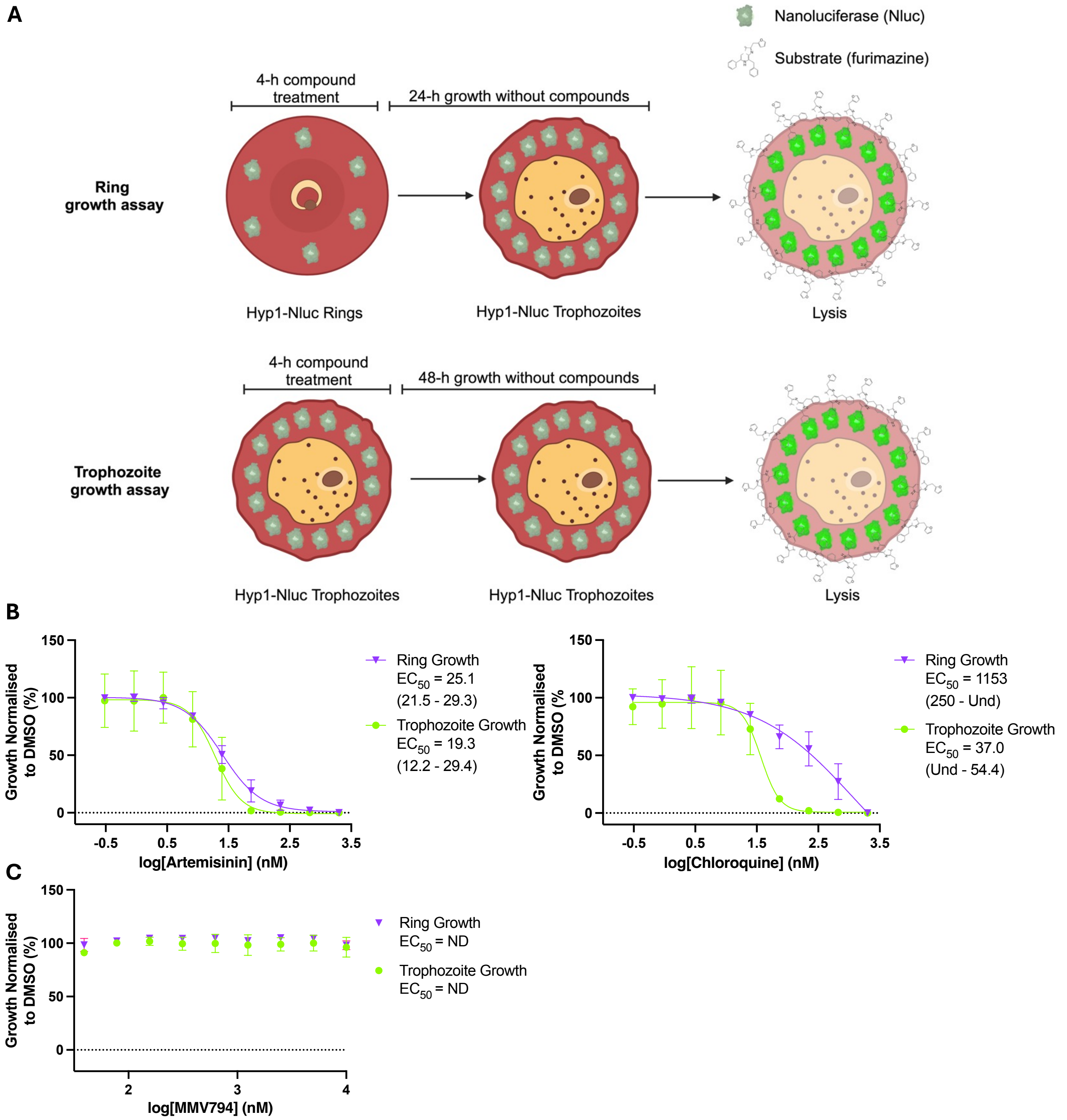
A
Nanoluciferase (Nluc)
Substrate (furimazine)
4-h compound treatment
24-h growth without compounds
Ring growth assay
Hyp1-Nluc Rings
Hyp1-Nluc Trophozoites
Lysis
4-h compound treatment
48-h growth without compounds
Trophozoite growth assay
Hyp1-Nluc Trophozoites
Hyp1-Nluc Trophozoites
Lysis
B
Growth Normalised to DMSO (%)
log[Artemisinin] (nM)
Ring Growth
EC50 = 25.1
(21.5 - 29.3)
Trophozoite Growth
EC50 = 19.3
(12.2 - 29.4)
Growth Normalised to DMSO (%
log[Chloroquine] (nM)
Ring Growth
EC50 = 1153
(250 - Und)
Trophozoite Growth
EC50 = 37.0
(Und - 54.4)
C
Growth Normalised to DMSO (%)
log[MMV794] (nM)
Ring Growth
EC50 = ND
Trophozoite Growth
EC50 = ND

**Figure S3. MMV794 does not inhibit ring or trophozoite growth. A** Schematics of ring and trophozoite growth assays. Hyp1-Nluc ring and trophozoites were treated with compounds for 4 h. Compounds were removed by washing, and successfully invaded parasites were grown for 24 or 48 h, lysed at the trophozoite stage, and furimazine was added to generate bioluminescence as a measure of ring or trophozoite growth. Figure created using BioRender.com. **B** Dose response curves of ring and trophozoite growth assays for control compounds, artemisinin and chloroquine (parasite growth inhibitors) were as expected, with chloroquine being more potent against trophozoite growth. **C** MMV794 did not inhibit ring or trophozoite growth after 4 h treatment. Error bars represent the standard deviation of the mean of three biological replicates, each with two or three technical replicates.

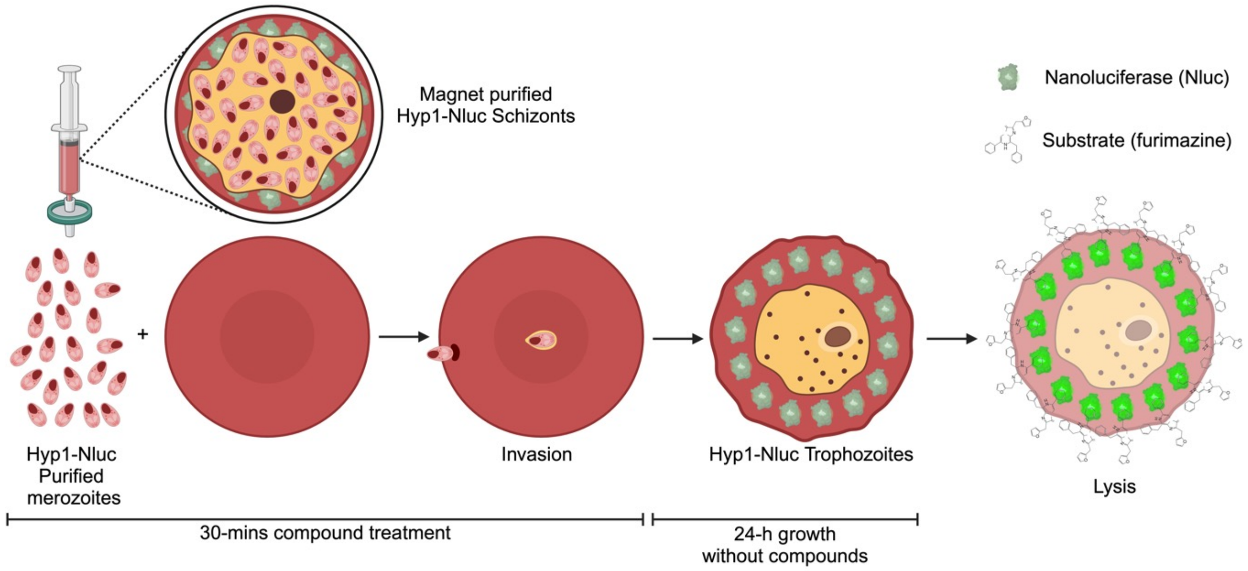


**Figure S4. Schematic of purified merozoite invasion assay.** Merozoites were isolated from late-stage magnet-purified Hyp1-Nluc schizonts and added to fresh red blood cells (RBCs) for 30 mins in the presence of compounds and agitation to stimulate invasion. Compounds and residual merozoites were removed by washing, and successfully invaded parasites were grown for 24 h, lysed at the trophozoite stage, and furimazine was added to generate bioluminescence as a measure of merozoite invasion. Figures created using BioRender.com.

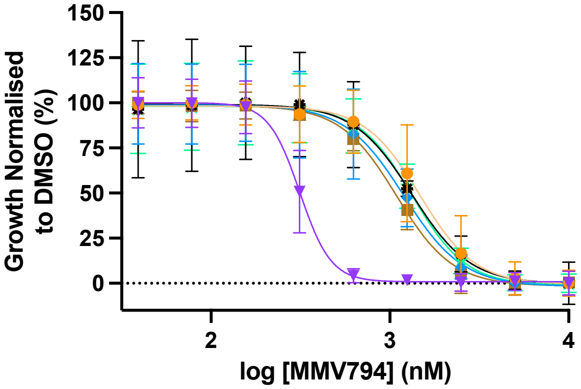


| | $EC_{50}$ (nM) (C.I.) |
|---|---|
| WT_3D7_F3 | 314 (275 – 357) |
| Pop B_F10 | 1474 (1140 – 1971) |
| Pop B_F6 | 1198 (870 – 1676) |
| Pop C_C6 | 1327 (863 – 2486) |
| Pop C_C8 | 1090 (955 – 1241) |
| Pop C_D6 | 1336 (1014 – 1807) |

**Figure S5. 72-h LDH growth assays of clonal MMV794-resistant parasites.** MMV794-resistant parasite populations B and C were cloned by limiting dilution to yield two clones of population B (Pop B_F10 and Pop B_F6) and three clones of population C (Pop C_C6, Pop C_C8 and Pop C_D6). The clonal parasites were tested against MMV794 for 72-h growth inhibition. Values were normalised to 0.1% DMSO (vehicle control). Error bars represent the standard deviation of the mean of three biological replicates, each with three technical replicates. Mean of $EC_{50}$ and 95% confidence interval (C.I.) values are indicated.

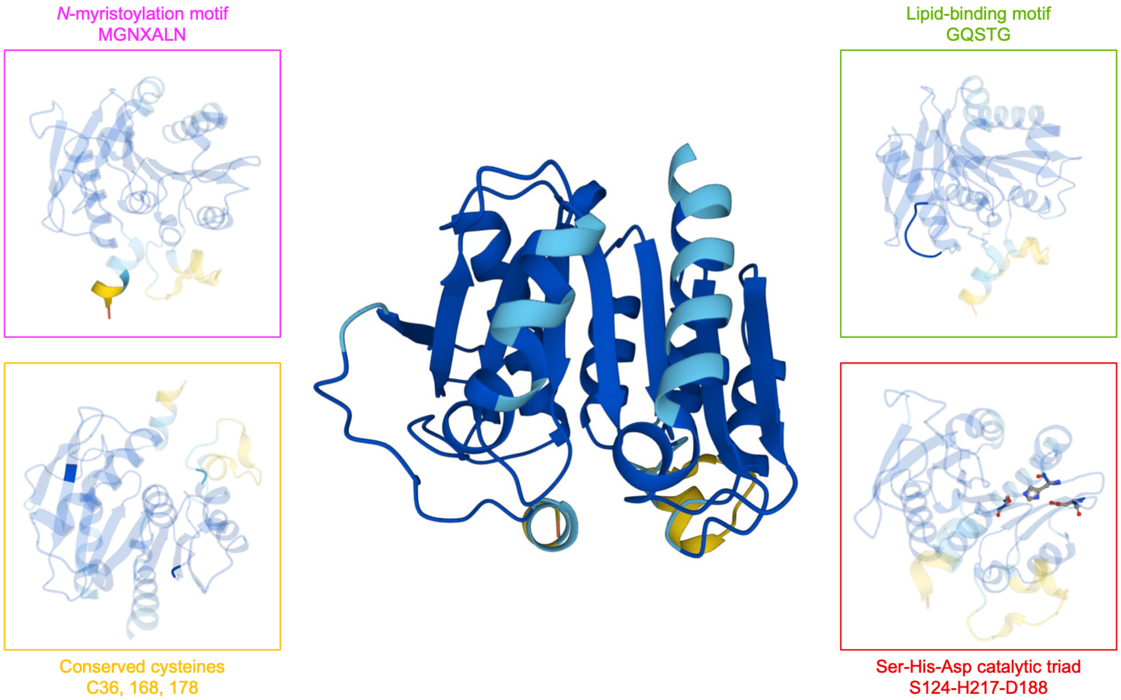


**Figure S6. 3D model of the α/β hydrolase fold in *Pf*SLIRP.** Model was predicted by AlphaFold (Jumper *et al.*, 2021; Varadi *et al.*, 2024). Magenta denotes *N*-myristoylation motif, orange/yellow denotes three conserved cysteines at positions 36, 168 and 178, green denotes the GXSXG lipid-binding motif and red denoting the catalytic triad composed of the nucleophile (serine 124), acid (aspartate 188) and histidine (127). The conserved features are highlighted in circles (2D) and boxes and opaque regions (3D) of the protein. Residues are colour-coded by AlphaFold per-residue model confidence score between 0-100: very high > 90 (blue), high > 70 (cyan), low > 50 (yellow) and orange < 50 (very low).

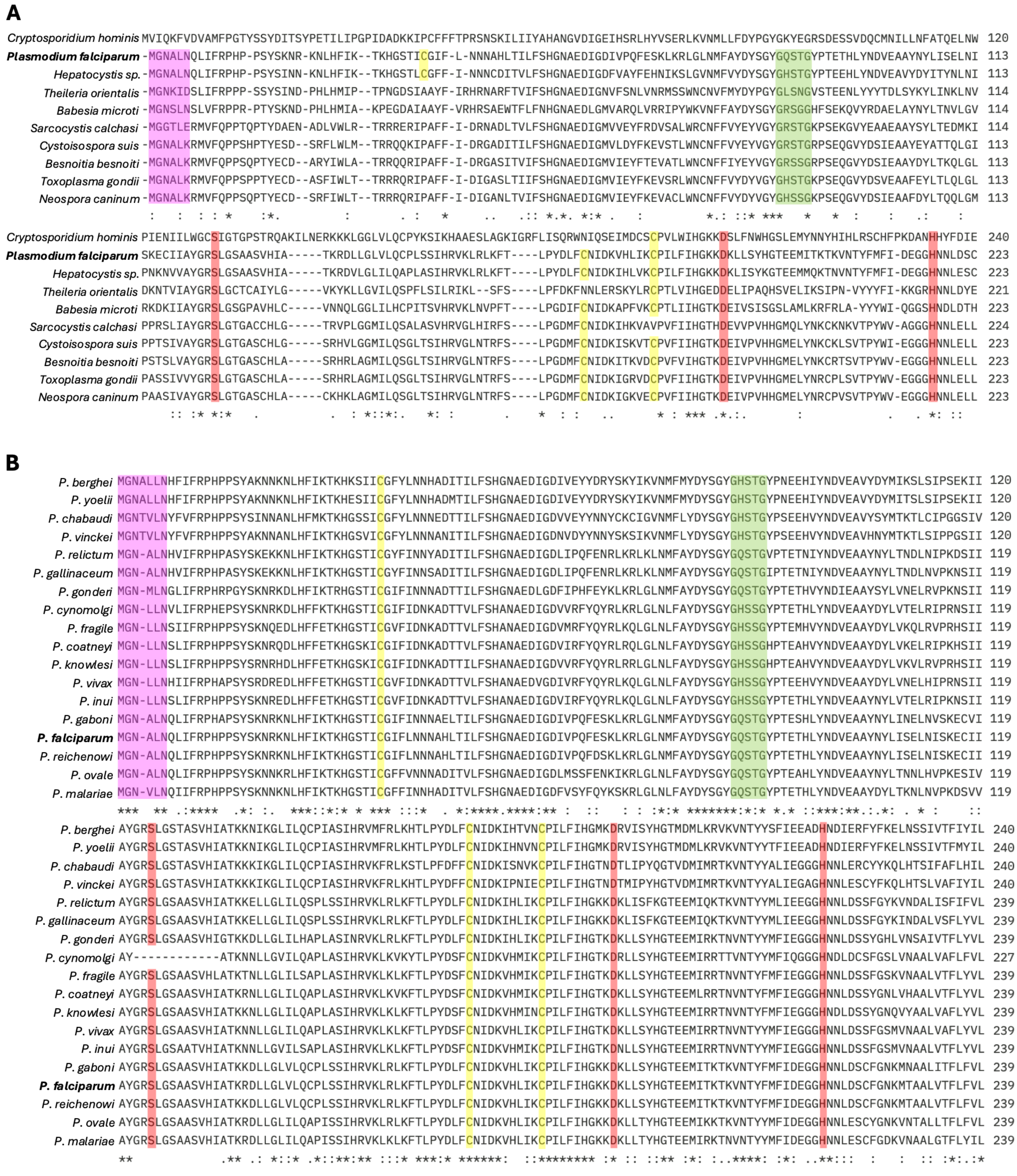


**Figure S7. Multiple sequence alignment of *Pf*SLIRP (*Plasmodium falciparum*) against its potential orthologs. A** In other Apicomplexan parasites. **B** In other *Plasmodium spp*. (*) = fully conserved residues; (:) = Strongly conserved residues; (.) = Weakly conserved residues. The *N*-myristoylation motif, three conserved cysteine residues, GXSXG lipid-binding motif and the Ser-His-Asp catalytic triad are highlighted in magenta, yellow, green, and red, respectively. Multiple sequence alignment was performed using Clustal Omega (Sievers *et al.*, 2011).

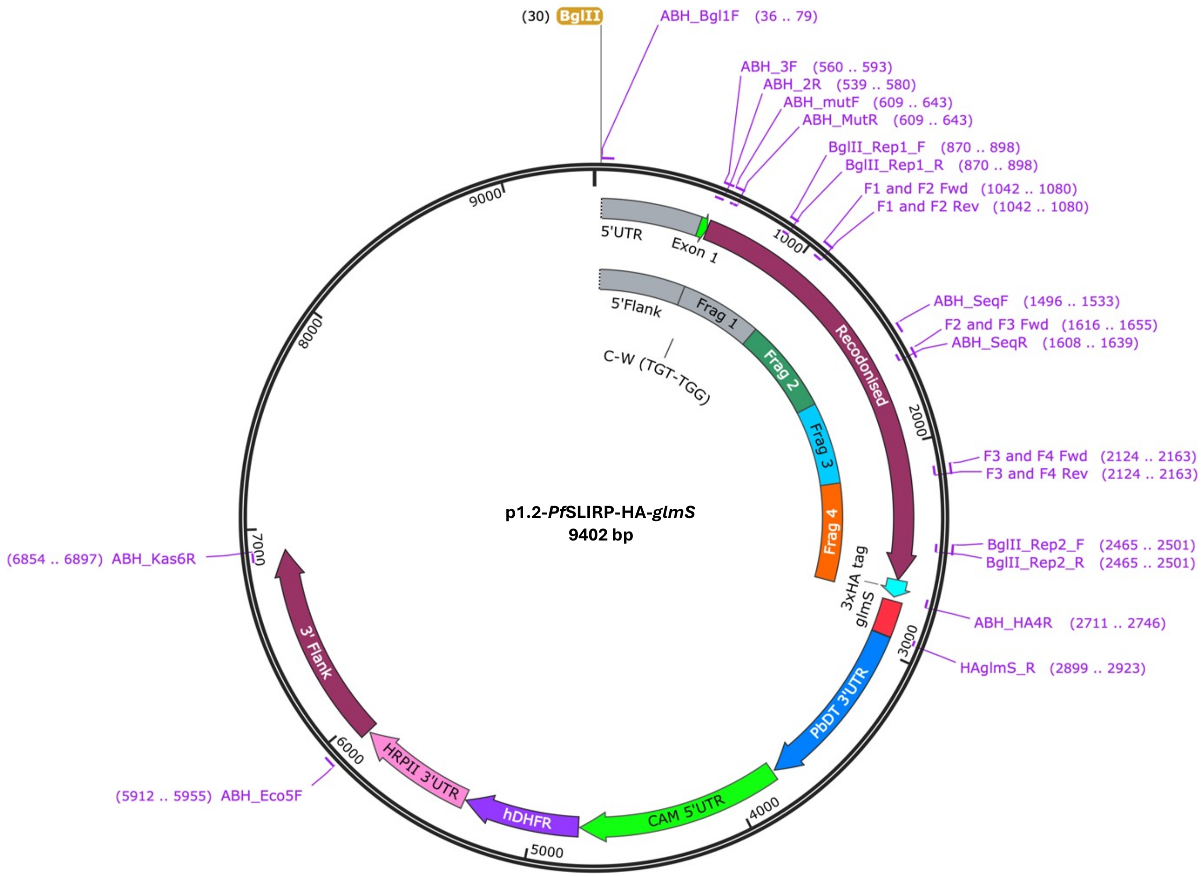


**Figure S8. Map of *Pf*SLIRP CRISPR-Cas9 construct, p1.2-*Pf*SLIRP-HA-*glmS***

The *BglII* site used to linearise the plasmid for transfection is highlighted in yellow. Figure created using SnapGene® software (Insightful Science; available at snapgene.com)

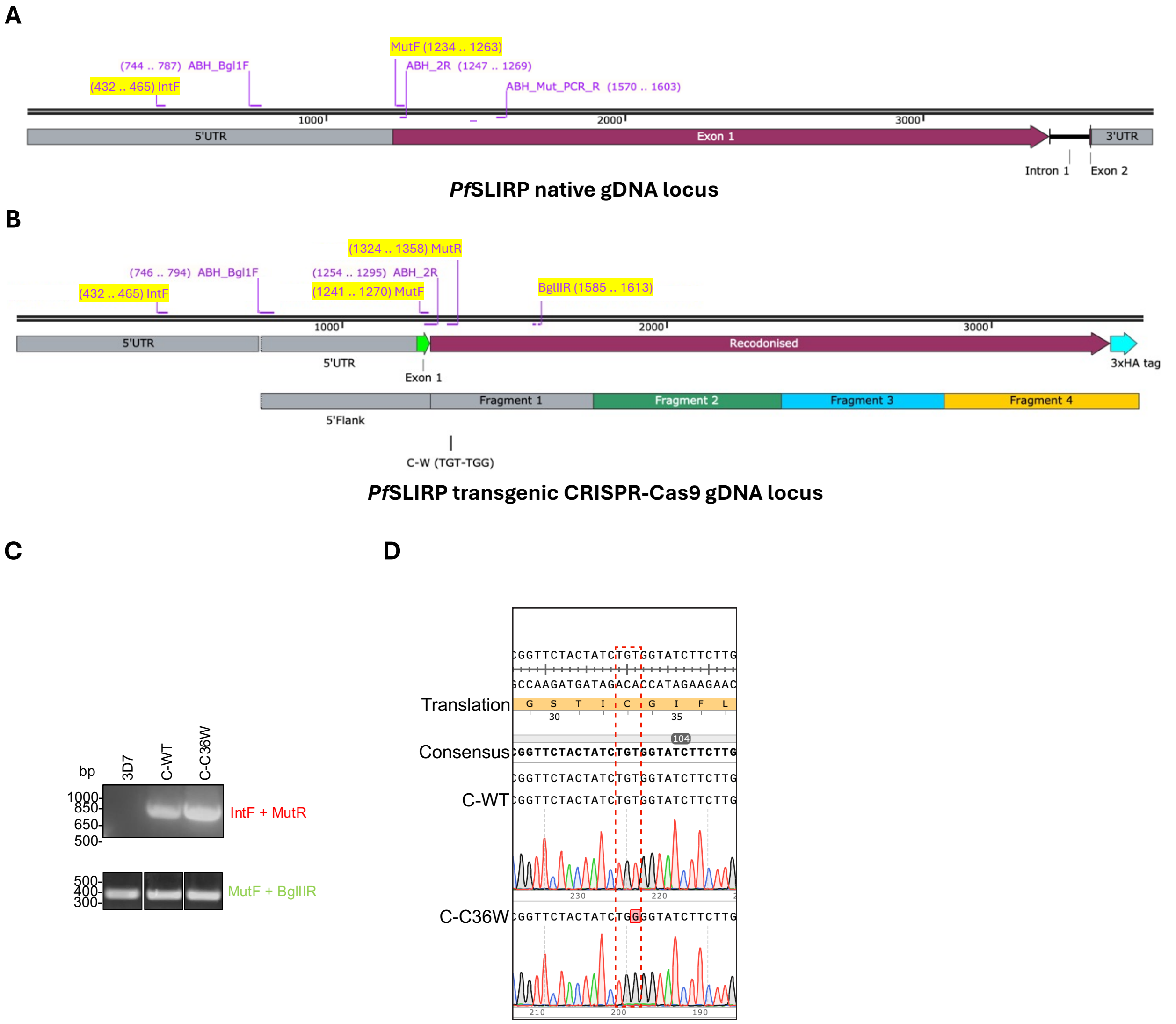


**Figure S9. PCR validation of transgenic *Pf*SLIRP parasites.** Map of **A** wild-type and **B** transgenic CRISPR-Cas9 *Pf*SLIRP gDNA loci. Primers used to confirm correct integration of *Pf*SLIRP 5’ homology flank and C36W mutation in the transgenic *Pf*SLIRP locus are highlighted in yellow. Figure created using SnapGene® software (Insightful Science; available at snapgene.com) **C** PCRs using 3D7, C-WT and C-C36W gDNA and primers IntF and MutR targeted the region containing recodonised sections of the *Pf*SLIRP 5’ homology flank, showing that only C-WT and C-C36W produced a product and not the parental wild-type 3D7 parasites. To validate correct insertion of C36W mutation only in the C-C36W parasites, PCRs with primers MutF and BglIIR targeting the region containing C36W of the *Pf*SLIRP 5’ homology flank were used to amplify regions containing the C36W mutation. This produced a PCR product for all parasites, which were extracted and Sanger sequenced. **D** Chromatograms of the Sanger-sequenced PCR products using primers MutF and BglIIR confirmed that the C36W mutation was correctly inserted into the C-C36W parasite line only.

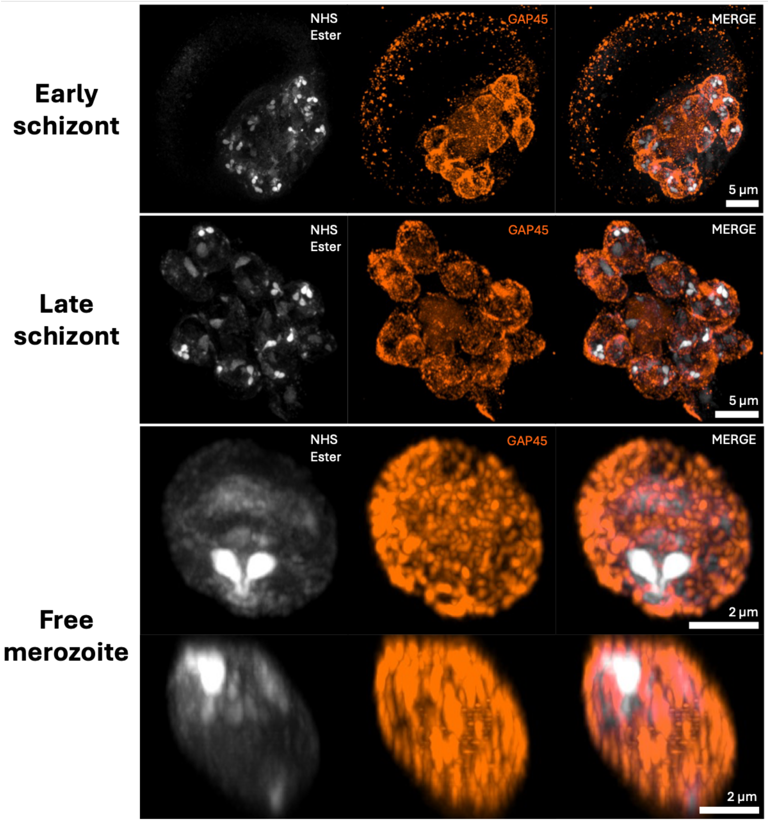


**Figure S10. Ultrastructure expansion microscopy of merozoites at different stages**

Maximum intensity projections of IFAs using C-WT samples shows that NHS ester stains protein density and GAP45 stains the merozoite periphery (inner membrane complex).

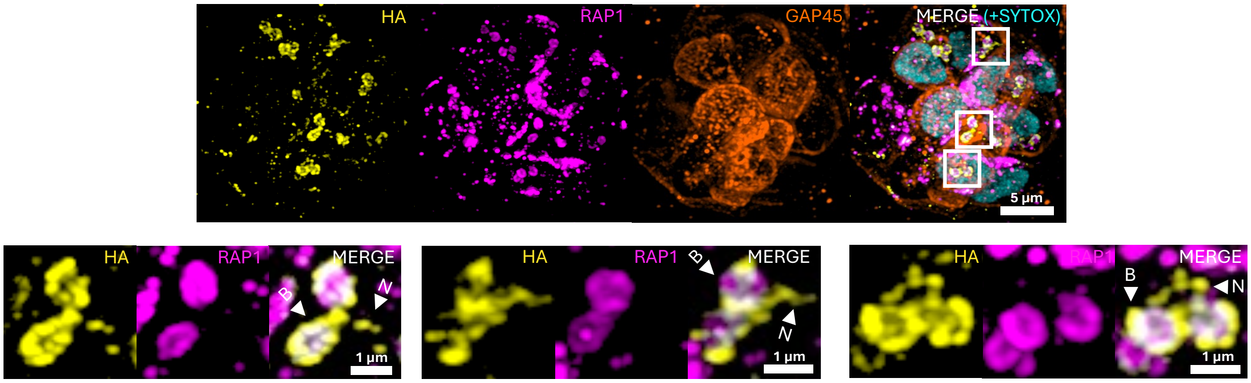


**Figure S11. Ultrastructure expansion microscopy of C-C36W schizonts indicate *Pf*SLIRP localises at the rhoptry bulb membrane** Maximum intensity projections of IFAs using C-C36W samples demonstrate that *Pf*SLIRP (HA) is localised to the rhoptry bulb membrane. RAP1 was used to label the rhoptry bulbs, GAP45 for merozoite periphery, and SYTOX™ blue for DNA. B denotes rhoptry bulb.

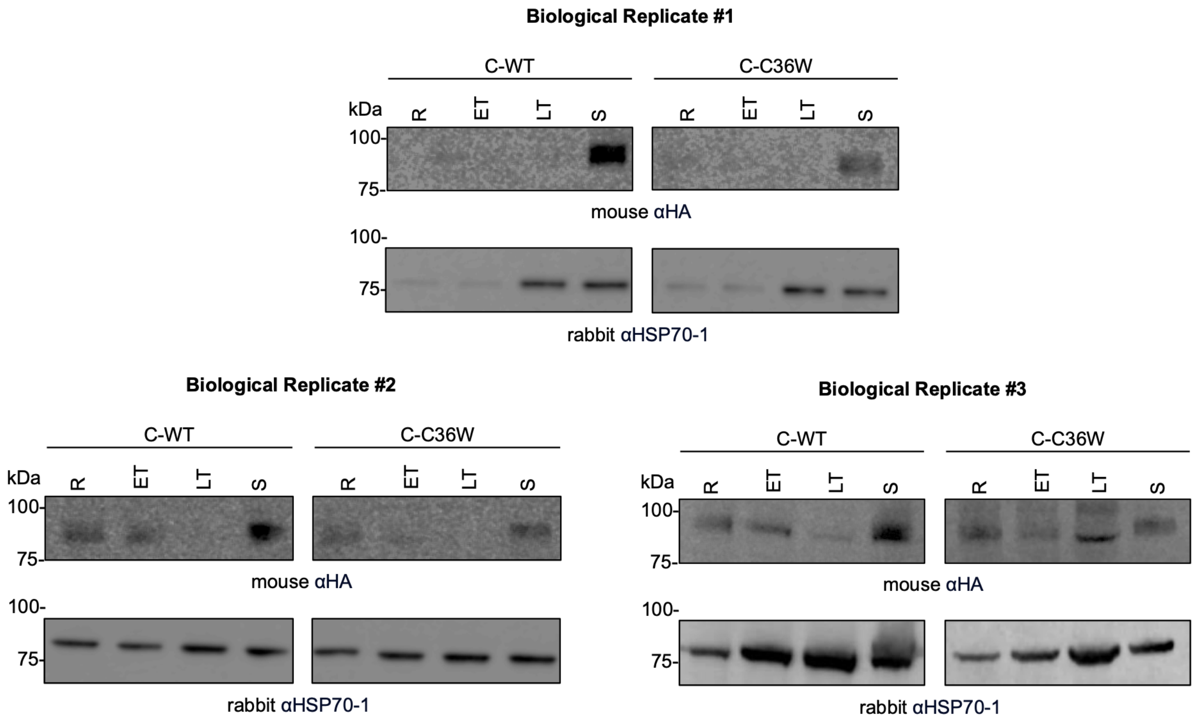


**Figure S12. Western blots used for time course analysis of *Pf*SLIRP expression at the ring, early trophozoite, late trophozoite and schizont stages.** C-WT and C-C36W parasites synchronised to rings (R), early trophozoites (ET), late trophozoites (LT) and schizonts (S) were saponin lysed and used for western blot and densitometry analysis to determine *Pf*SLIRP expression levels. Mouse anti-HA was used to detect *Pf*SLIRP, and rabbit anti-HSP70-1 was used to detect HSP70-1.

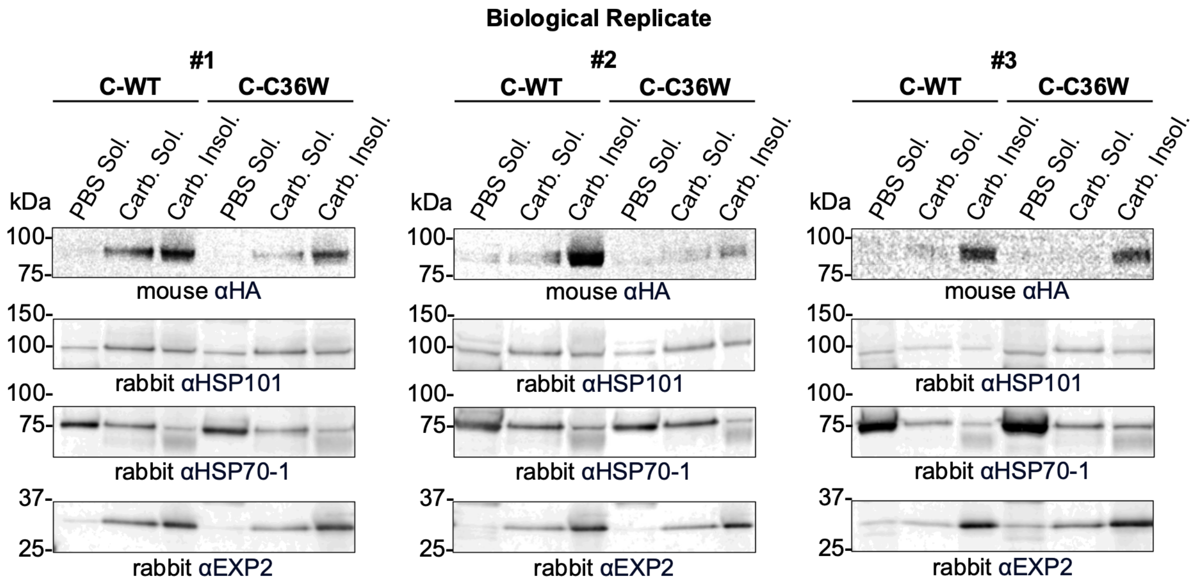


**Figure S13. *Pf*SLIRP is associated with membranes.** Sequential lysis of C-WT and C-C36W schizont saponin pellets via sodium carbonate (Carb.) extraction were conducted to examine the solubility of *Pf*SLIRP. In this assay, soluble proteins are found in the PBS soluble (Sol.) fractions, peripheral membrane proteins in the carbonate soluble fraction, integral membrane proteins in the carbonate insoluble (Insol.) fraction. Western blots show controls behaved as expected with soluble protein HSP70-1 being most concentrated in the PBS soluble fraction, peripheral membrane protein HSP101 most concentrated in the carbonate soluble fraction, and integral membrane protein EXP2 most concentrated in the carbonate insoluble fraction. *Pf*SLIRP was detected in the carbonate soluble and carbonate insoluble fractions and is therefore membrane associated. Mouse anti-HA was used for detecting *Pf*SLIRP. Rabbit anti-HSP101, HSP70-1 and EXP2 were used as controls.

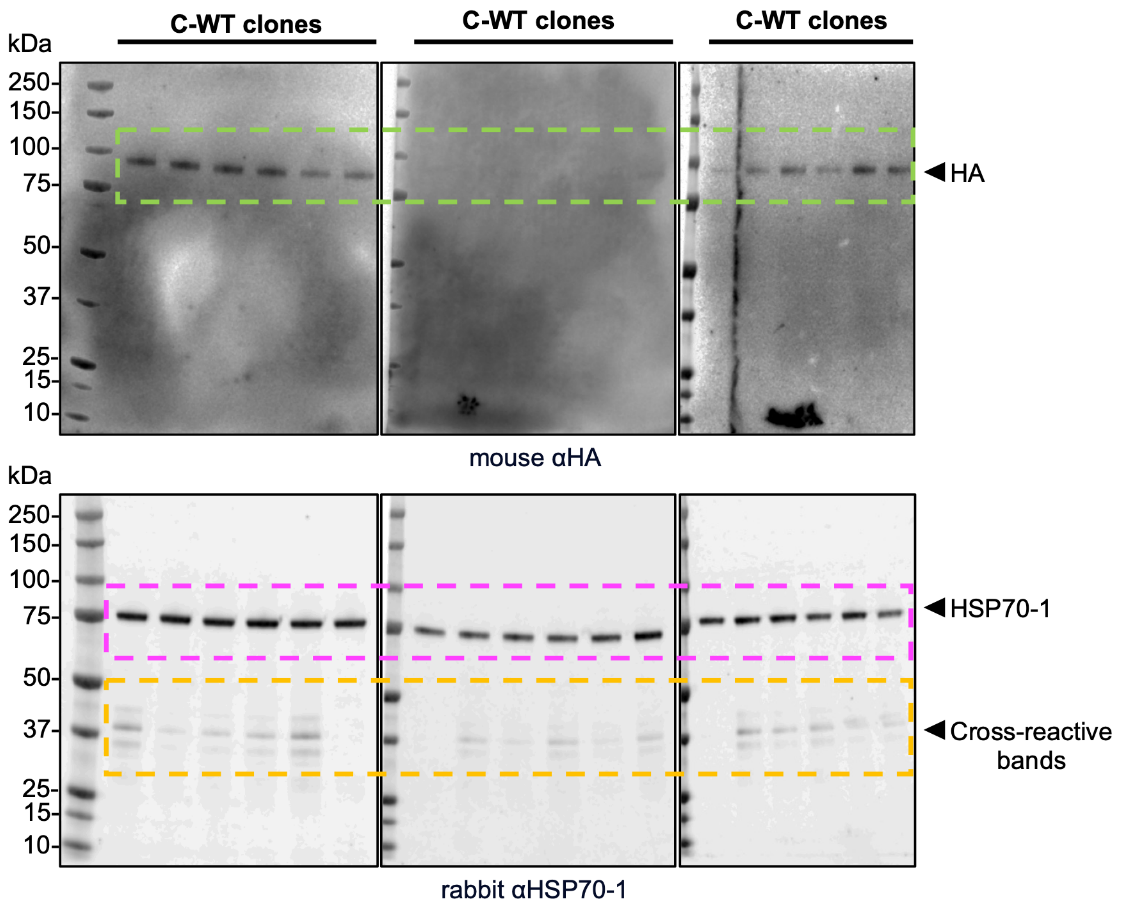


**Figure S14. Rabbit anti-HSP70-1 antibody produces cross-reactive bands in western blots.** C-WT clones isolated via limiting dilution were saponin lysed at schizont stage and validated for HA-tagging by western blot. Mouse anti-HA was used to detect *Pf*SLIRP, and rabbit anti-HSP70-1 was used to detect HSP70-1. Rabbit anti-HSP70-1 produced cross-reactive bands at ~35-45 kDa.

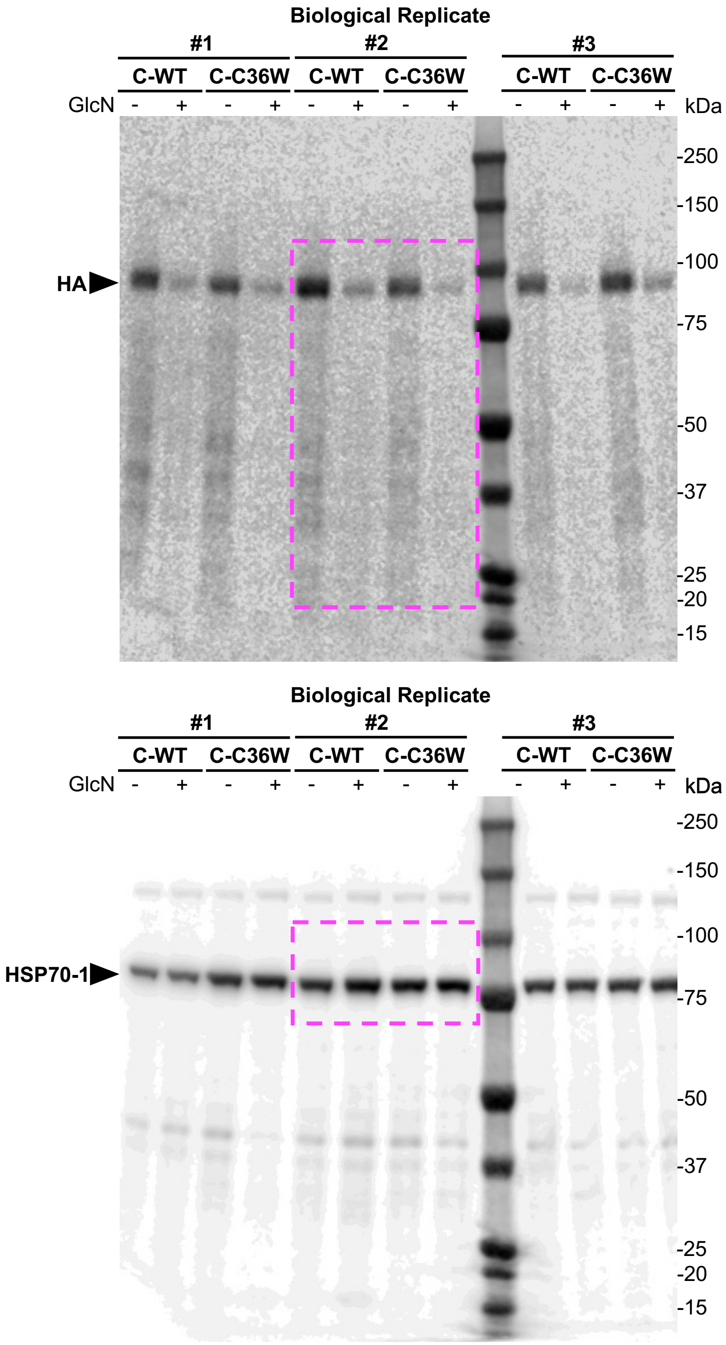


**Figure S15. Western blots used for densitometry analysis of *Pf*SLIRP knockdown.** Western blots of proteins extracted from C-WT and C-C36W parasites ± 2.5 mM glucosamine (GlcN) for 72 h (early trophozoites to schizont) demonstrating *Pf*SLIRP knockdown. Densitometry analyses were performed by normalising signal from *Pf*SLIRP (HA) bands to HSP70-1. Magenta box indicates regions used for Figure 6. Mouse anti-HA was used to probe *Pf*SLIRP, and rabbit anti-HSP70-1 was used as a loading control.

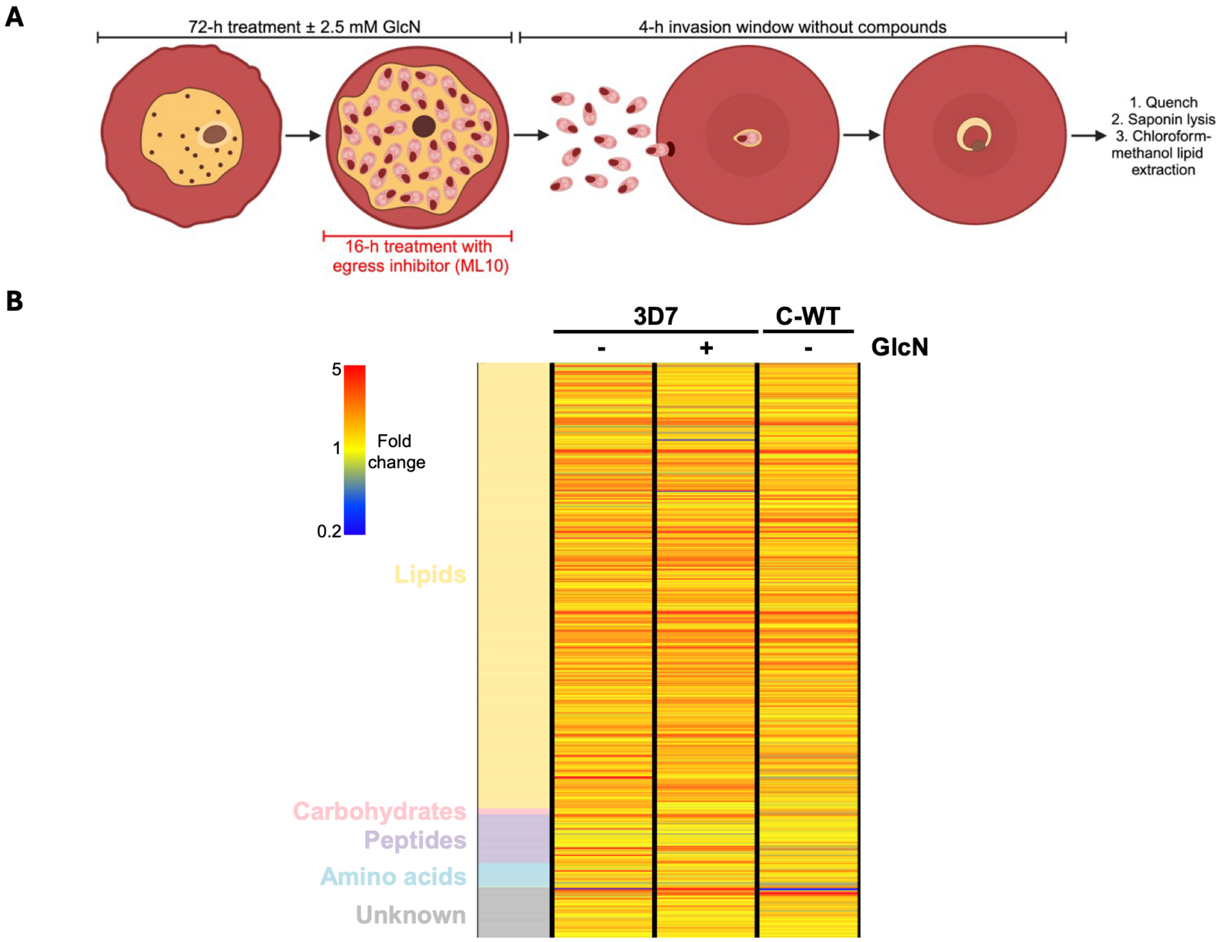


**Figure S16. C-WT parasites have a comparable metabolome to 3D7 parasites and glucosamine does not perturb parasite metabolites. A** Early trophozoite C-WT and C-C36W parasites were treated ± 2.5 mM glucosamine (GlcN) for 72 h, during which the egress inhibitor, ML10, was added at the 56 h timepoint for 16 h (overnight) to stall parasites at the schizont stage. GlcN and ML10 was removed at the 72 h timepoint, and parasites were allowed to egress and invade for 4 h, after which time, lipids were harvested for mass spectrometry analysis. Figure created using BioRender.com. **B** Heatmap analysis of all metabolite abundances (y axis) expressed as mean fold change versus untreated 3D7 (3D7 -) control in all sample groups (x axis). The three parasite metabolite profiles were comparable with no notable changes in particular metabolite group. Each column represents a single biological replicate. Yellow indicates no significant perturbation of metabolite levels relative to control while red and blue indicates increased and decreased abundances, respectively.

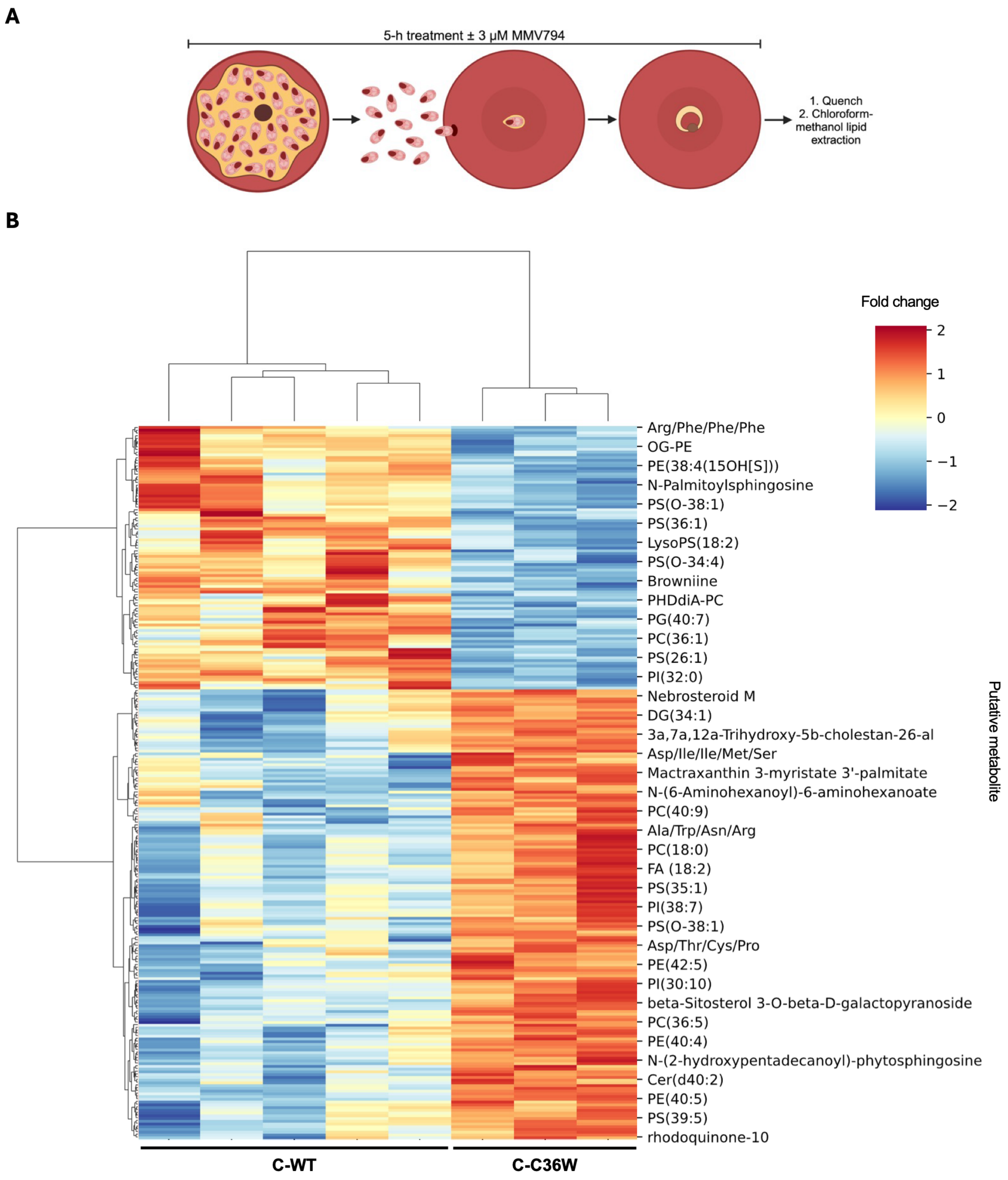


**Figure S17. The C36W mutation in *Pf*SLIRP perturbs the parasite metabolome. A** Purified schizonts were treated ± 3 µM MMV794 for 5 h, during which parasites egressed and invaded red blood cells for 5 h, parasites were subsequently harvested for metabolomic analyses. Figure created using BioRender.com **B** Hierarchical clustering of 260 significantly perturbed putative metabolites (fold change > 1.5 and $p < 0.01$) between untreated C-WT and C-C36W parasites. Vertical clustering shows sample similarities and horizontal clustering shows the relative abundance of the 260 putative metabolites. Each column represents a single biological replicate. The colour scale bar represents fold change (mean-centred and divided by standard deviation of each variable). Annotated metabolites (-) indicate validated sequence by MS/MS.

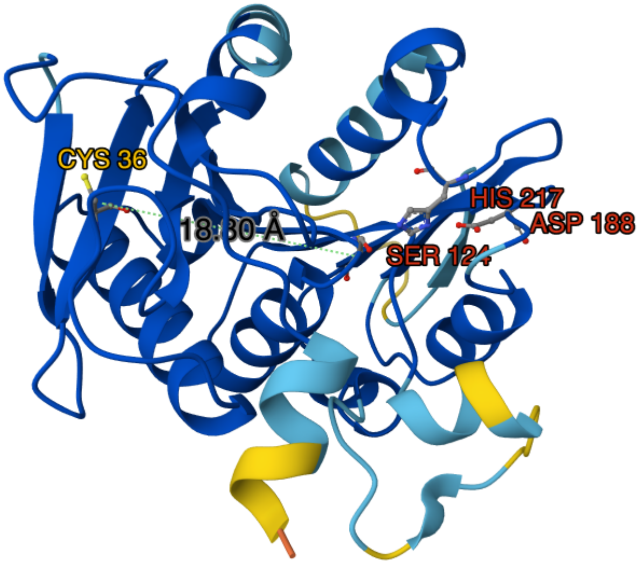


**Figure S18. The C36 in *Pf*SLIRP is approximately 19 Å from its catalytic serine.** The C36 residue in *Pf*SLIRP that underwent mutation (C36W) in MMV794-selected parasites is ~19 Å from the closest residue of the catalytic triad, S124. Model was predicted by AlphaFold (Jumper *et al.*, 2021; Varadi *et al.*, 2024). Orange/yellow denotes the conserved cysteine at position 36, and red denotes the catalytic triad composed of the nucleophile (serine 124), acid (aspartate 188) and histidine (127). Residues are colour-coded by AlphaFold per-residue model confidence score between 0-100: very high > 90 (blue), high > 70 (cyan), low > 50 (yellow) and orange < 50 (very low).

### Supplementary Tables

| Chromosome | Position | Ref | Alt | Effect | Gene_ID | Gene_Description | Codon_ Change | AA_ Change | WT_3D7_F3 | Pop B_F10 | Pop B_F6 | Pop C_C6 | Pop C_C8 | Pop C_D6 |
|---|---|---|---|---|---|---|---|---|---|---|---|---|---|---|
| Pf3D7_04_v3 | 213747 | A | G | nonsynonymous | PF3D7_0403800 | alpha/beta hydrolase, putative | TGT/TGG | C36W | | | | ✓ | ✓ | ✓ |
| Pf3D7_04_v3 | 214947-217591 | N/A | N/A | deletion (2644) | PF3D7_0403800 | alpha/beta hydrolase, putative | N/A | N/A | | ✓ | | | | |
| Pf3D7_04_v3 | 214947-217593 | N/A | N/A | deletion (2646) | PF3D7_0403800 | alpha/beta hydrolase, putative | N/A | N/A | | | ✓ | | | |
| Pf3D7_04_v3 | 340656 | G | A | nonsynonymous | PF3D7_0406500 | NYN domain-containing protein, putative | GAC/GAA | D1133E | | ✓ | | | | |
| Pf3D7_04_v3 | 340715 | T | C | nonsynonymous | PF3D7_0406500 | NYN domain-containing protein, putative | AAA/CAA | K1114Q | | | | | | ✓ |
| Pf3D7_04_v3 | 471930 | G | A | nonsynonymous | PF3D7_0410300 | protein phosphatase PPM1, putative | CCA/CAA | P556Q | | | ✓ | | | ✓ |
| Pf3D7_10_v3 | 1437050 | T | A | nonsynonymous | PF3D7_1036400 | liver stage antigen 1 | CAT/CAA | H245Q | | ✓ | | ✓ | | |
| Pf3D7_10_v3 | 1437082 | G | T | nonsynonymous | PF3D7_1036401 | liver stage antigen 1 | CGT/CTT | R256L | | | | | | ✓ |
| Pf3D7_10_v3 | 1437383 | T | G | nonsynonymous | PF3D7_1036402 | liver stage antigen 1 | GAT/GAG | D356E | | ✓ | | ✓ | | |
| Pf3D7_10_v3 | 1437541 | G | T | nonsynonymous | PF3D7_1036403 | liver stage antigen 1 | CGT/CTT | R409L | | | | ✓ | | |
| Pf3D7_13_v3 | 122989-128525 | N/A | N/A | inversion (5536) | PF3D7_1302200, PF3D7_1302300 | early transcribed membrane protein 13 (ETRAMP13); Plasmodium exported protein, unknown function | N/A | N/A | | | | | ✓ | |
| Pf3D7_13_v4 | 122989-128595 | N/A | N/A | inversion (5606) | | | N/A | N/A | | | | | | ✓ |
| Pf3D7_13_v5 | 123067-128528 | N/A | N/A | inversion (5461) | | | N/A | N/A | | | ✓ | | | |
| Pf3D7_05_v3 | 755918 | G | A | nonsynonymous | PF3D7_0518100 | protein AMR2 | GGT/GAT | G874D | | | | ✓ | | |
| Pf3D7_07_v3 | 1080009 | G | A | nonsynonymous | PF3D7_0725400 | conserved Plasmodium protein, unknown function | AGC/AAC | S95N | | | | ✓ | | |
| Pf3D7_08_v3 | 959318 | C | T | nonsynonymous | PF3D7_0821300 | ATP-dependent RNA helicase DHX36, putative | TGT/TAT | C606Y | | ✓ | | | | |
| Pf3D7_11_v3 | 730107 | C | A | nonsynonymous | PF3D7_1119400 | ubiquitin-protein ligase, putative | CAA/AAA | Q738K | | | | ✓ | | |
| Pf3D7_11_v3 | 1120609 | G | T | nonsynonymous | PF3D7_1128900 | conserved protein, unknown function | TGG/TTG | W598L | | ✓ | | | | |
| Pf3D7_11_v3 | 1861293 | C | T | nonsynonymous | PF3D7_1147000 | sporozoite and liver stage asparagine-rich protein | TGT/TAT | C2022Y | | | | ✓ | | |
| Pf3D7_12_v3 | 646127 | G | A | nonsynonymous | PF3D7_1216000 | serine--tRNA ligase, putative | AGC/AAC | S530N | | | | ✓ | | |
| Pf3D7_14_v3 | 1821851 | G | T | nonsynonymous | PF3D7_1444100 | conserved Plasmodium protein, unknown function | TCT/TAT | S866Y | | | ✓ | | | |

**Table S1. Tabular summary of pooled variant analysis of MMV794-resistant parasites.** Table depicts non-synonymous single nucleotide polymorphisms and structural variants (deletion and inversion) that passed quality filtration and were detected in at least one resistant parasite clone. Ref = reference allele. Alt = alternative allele. AA = amino acid. N/A = not applicable.

| Organism | Gene ID | BLAST Total Score[a] | BLAST Query Cover (%)[b] | BLAST Percentage Identity (%)[c] | Reciprocal BLAST hit[d] | Mammalian host cell type (Genus)[e] |
|---|---|---|---|---|---|---|
| *Plasmodium falciparum* | PF3D7_0403800 | N/A | N/A | N/A | N/A | Hepatocytes, erythrocytes, reticulocytes[1] |
| *Hepatocystis sp.* | HEP_00334600 | 2.00E-156 | 39 | 74.22 | ✓ | Hepatocytes, erythrocytes[2] |
| *Babesia microti* | BmR1_04g07110 | 5.00E-73 | 34 | 46.15 | ✓ | Erythrocytes[3] |
| *Theileria orientalis* | MACJ_002185 | 9.00E-62 | 34 | 43.70 | ✓ | Lymphocytes, leukocytes, erythrocytes[4] |
| *Cystoisospora suis* | CSUI_001334 | 3.00E-61 | 30 | 44.14 | X (PF3D7_0805000) | Nucleated cells only (intestinal epithelial cells)[5] |
| *Toxoplasma gondii* | TGME49_262490 | 4.00E-61 | 30 | 43.24 | X (PF3D7_0805000) | Nucleated cells only (Macrophages, epithelial cells, muscle cells, neurons, fibroblasts, enterocytes)[6,7] |
| *Neospora caninum* | NCLIV_025340 | 7.00E-61 | 30 | 42.79 | X (PF3D7_0805000) | Nucleated cells only (Neurons, macrophages, fibroblasts, vascular endothelial cells, myocytes, hepatocytes and dermal cells)[8] |
| *Besnoitia besnoiti* | BESB_001740 | 2.00E-60 | 30 | 41.44 | X (PF3D7_0805000) | Nucleated cells only (endothelial cells and fibroblasts)[9] |
| *Sarcocystis calchasi* | SCGI_LOCUS1799 | 5.00E-59 | 30 | 42.34 | X (PF3D7_0805000) | Nucleated cells only (Skeletal muscle cell and cardiac muscle cell)[10] |
| *Porospora gigantea* | GNI_096520 | 2.00E-34 | 26 | 40.83 | X (PF3D7_0805000) | Nucleated cells only (intestinal epithelial cells)[11,12] |
| *Cryptosporidium hominis* | CHM_1g2570 | 2.00E-21 | 22 | 35.63 | ✓ | Nucleated cells only (intestinal epithelial cells)[13] |

**Table S2. BLAST search of *Pf*SLIRP search against other Apicomplexan parasites identifies potential *Pf*SLIRP orthologues only in blood-borne parasites.**

[a]Total score as calculated by BLAST (Altschul *et al.*, 1990) based on the sum of alignment scores for all High-Scoring Segment Pairs between the query and subject sequence, derived from substitution matrices, gap penalties, and match/mismatch values. Scores closer to 0 indicates stronger homology.

[b]The percentage of the query sequence (PF3D7_0403800) that aligns with the subject sequence in the BLAST search, indicating the length of the query sequence that is covered by the alignment.

[c]Percentage of exact matches between the query (PF3D7_0403800) and subject sequence within the aligned region. BLAST calculates it by dividing the number of identical matches by the total aligned positions, including gaps, and multiplying by 100 to express it as a percentage.

[d]A reciprocal BLAST hit occurs when two sequences (PF3D7_0403800 and other proteins) from different species are BLAST searched against each other, and each returns the other as its top hit, indicating potential orthology.

[e]Type of mammalian host cell types that may be parasitised by the Genus of Apicomplexan parasites (Feufack-Donfack *et al.*, 2024)[1], (Ejotre *et al.*, 2021)[2], (Uilenberg, 2006)[3], (Lakew *et al.*, 2023)[4], (Shrestha *et al.*, 2015)5, (Attias *et al.*, 2020)[6], (Dubey *et al.*, 1998)[7], (Fisher *et al.*, 2024)8, (Söderström *et al.*, 2021)[9], (Frey *et al.*, 2016)[10], (Valigurová, 2012)[11], (Florent *et al.*, 2021)[12], (Guérin and Striepen, 2020)[13].

N/A = not applicable.

Highlighted rows indicate potential *Pf*SLIRP orthologues with ≥ 40 % similar in percentage identity and reciprocal BLAST hits.

| Organism | Gene ID | BLAST Total Score[a] | BLAST Query Cover (%)[b] | BLAST Percentage Identity (%)[c] | Reciprocal BLAST hit[d] |
|---|---|---|---|---|---|
| *P. reichenowi* | PRSY57_0401400 | 0 | 100 | 92.72 | ✓ |
| *P. gaboni* | PGABG01_0402000 | 0 | 100 | 78.98 | ✓ |
| *P. ovale* | POVWA1_004930 | 0 | 68 | 68.36 | ✓ |
| *P. malariae* | PmUG01_03013800 | 0 | 96 | 49.52 | ✓ |
| *P. gonderi* | PGO_030170 | 5.00E-170 | 51 | 67.11 | ✓ |
| *P. relictum* | PRELSG_0301000 | 6.00E-170 | 50 | 69.35 | ✓ |
| *P. inui* | C922_04786 | 2.00E-169 | 76 | 50.09 | ✓ |
| *P. vivax* | PVW1_030010200 | 9.00E-166 | 51 | 63.81 | ✓ |
| *P. coatneyi* | PCOAH_00006120 | 1.00E-162 | 51 | 64.53 | ✓ |
| *P. knowlesi* | PKNH_0301800 | 1.00E-162 | 96 | 42.16 | ✓ |
| *P. fragile* | AK88_02005 | 5.00E-162 | 51 | 63.2 | ✓ |
| *P. gallinaceum* | PGAL8A_00054400 | 2.00E-160 | 50 | 69.32 | ✓ |
| *P. cyonomolgi* | PCYB_031150 | 6.00E-148 | 51 | 61.33 | ✓ |
| *P. berghei* | PBANKA_1001500 | 2.00E-128 | 49 | 53.3 | ✓ |
| *P. yoelii* | PY17XNL_01002645 | 2.00E-127 | 50 | 53.02 | ✓ |
| *P. vinckei* | YYG_02575 | 1.00E-125 | 55 | 51.21 | ✓ |
| *P. chabaudi* | PCHDS_000221700 | 2.00E-116 | 49 | 52.04 | ✓ |

**Table S3. Tabular summary of basic local alignment search tool results of *Pf*SLIRP search against other *Plasmodum spp*.**

[a]Total score as calculated by BLAST (Altschul *et al.*, 1990) based on the sum of alignment scores for all High-Scoring Segment Pairs between the query and subject sequence, derived from substitution matrices, gap penalties, and match/mismatch values. Scores closer to 0 indicates stronger homology.

[b]The percentage of the query sequence (PF3D7_0403800) that aligns with the subject sequence in the BLAST search, indicating the length of the query sequence that is covered by the alignment.

[c]Percentage of exact matches between the query (PF3D7_0403800) and subject sequence within the aligned region. BLAST calculates it by dividing the number of identical matches by the total aligned positions, including gaps, and multiplying by 100 to express it as a percentage.

[d]A reciprocal BLAST hit occurs when two sequences (PF3D7_0403800 and other proteins) from different species are BLAST searched against each other, and each returns the other as its top hit, indicating potential orthology.

A BLAST search of *Pf*SLIRP was performed against other *Plasmodium* parasites, which found *Pf*SLIRP orthologues in 17 species (≥ 40 % similar in percentage identity and reciprocal BLAST hits).

| Name | Construction of *Pf*SLIRP 5' homology flank for p1.2- *Pf*SLIRP-HA-*glmS* |
|---|---|
| ABH_Bgl1F | AGATCTTGTTTATGTTTGAATAGGTTATTTTTTTGTGTAACCCCT TTGA |
| ABH_2R | TCTGTTCTTAGAGTAAGATGGTGGGTGCGGTCTGAAGATTAG |
| ABH_3F | CCATCTTACTCTAAGAACAGAAAGAACTTGCACT |
| F1 and F2 Rev | ACCGTGGATGAACAAGATTGGACACT |
| F1 and F2 Fwd | AGTGTCCAATCTTGTTCATCCACGGT |
| ABH_SeqR | AACAGAGTTCTTTTCACACAAAGTGTTAGAGT |
| F2 and F3 Fwd | ACTTTGTGTGAAAAGAACTCTGTTCAAAGGAACAACAACA |
| F3 and F4 Rev | TCGATTTGTTCAGAGATGATGTTCTTCTTCATATTATCGA |
| F3 and F4 Fwd | TCGATAATATGAAGAAGAACATCATCTCTGAACAAATCGA |
| ABH_HA4R | ACTAGTTTAGGCATAATCTGGAACATCGTACGGATA |
| **Name** | **Inserting C36W mutation into *Pf*SLIRP 5' homology flank** |
| ABH_MutF (MutF) | ACGGTTCTACTATCTGGGGTATCTTCTTGAACAAC |
| ABH_MutR (MutR) | GTTGTTCAAGAAGATACCCCAGATAGTAGAACCGT |
| **Name** | **Removing internal *BglII* sites from *Pf*SLIRP 5' homology flank** |
| BglII_Rep1_F | TCATCGCTTACGGTAGGTCTTTGGGTTCT |
| BglII_Rep1_R (BglIIR) | AGAACCCAAAGAcCTACCGTAAGCGATGA |
| BglII_Rep2_F | ACTGAAAGAAAGGACAAAATCTCTATCAACAACGACA |
| BglII_Rep2_R | TGTCGTTGTTGATAGAGATTTTGTCCTTTCTTTCAGT |
| **Name** | **Guide RNA for insertion of p1.2- *Pf*SLIRP-HA-*glmS* into WT *P. falciparum*** |
| ABH-sgRNA | CTATTCTTTGAATAGCTAGGTGG (protospacer adjacent motif (PAM) underlined) |

**Table S4. Sequences of primers used to generate the *Pf*SLIRP CRISPR-Cas9 construct, p1.2-*Pf*SLIRP-HA-glmS**

| **Name** | **Confirming integration of the *Pf*SLIRP 5’ homology flank into the genomic locus following transfection of the p1.2 CRISPR plasmid** |
| --- | --- |
| ABH.p1.2-IntF (IntF) | AAATTAAGCTAATATGTTTATATGCTCATGACGA |
| ABH_MutR (MutR) | GTTGTTCAAGAAGATACCCCAGATAGTAGAACCGT |
| **Name** | **Confirming integration of the *Pf*SLIRP C36W mutation into the genomic locus following transfection of the p1.2-*Pf*SLIRP (C36W)-HA-*glmS*** |
| ABH_Mut_PCR_F (MutF) | TGCATTGAACCAACTAATCTTCAGACCGCA |
| BglII_Rep1_R (BglIIR) | AGAACCCAAAGACCTACCGTAAGCGATGA |

**Table S5. Sequences of primers used to validate correct integration of the *Pf*SLIRP 5’ homology flank and C36W mutation in CRISPR-Cas9 constructs, p1.2-*Pf*SLIRP(WT)-HA-glmS and p1.2-*Pf*SLIRP(C36W)-HA-glmS**

| Parameter | Definition |
| --- | --- |
| Egress | Schizont rupture leading to the release of merozoites from schizonts and RBCs. |
| RBC contact | Contact between a merozoite and RBC lasting ≥ 0.25 s. |
| RBC contacts per Egress | The number of contacts between a merozoite and RBC lasting ≥ 0.25 s from a single egress event. |
| RBC deformation | The process by which merozoites cause indentations on the RBC surface after contacting the RBC. |
| RBC deformation per RBC contact | The proportion of merozoite contacting a RBC that proceeded to cause indentations on the RBC surface. |
| Internalisation | The process by which the merozoite drives itself into the RBC. |
| Internalisation per RBC contact | The proportion of merozoite contacting a RBC that proceeded to drives itself into the RBC. |
| Ejection | When a merozoite is ejected from the RBC after internalisation. |
| Invasion | When a merozoite internalises itself within the RBC and remains internalised. |
| Invasion per RBC contact | The proportion of merozoite contacting a RBC that proceeded to internalises itself within the RBC and remains internalised. |

**Table S6. Parameters measured in live-cell imaging**

| Primary Antibody | Dilution | Immunogen | Supplier |
|---|---|---|---|
| Mouse monoclonal anti-HA | 1:1000 | Synthetic peptide from human influenza HA-tag, conjugated to Keyhole Limpet Haemocyanin | Sigma-Aldrich (Catalogue number H3663) |
| Rabbit polyclonal anti-EXP2 | 1:1000 | Recombinant EXP2 (Bullen *et al.*, 2012) | WEHI monoclonal facility |
| Rabbit polyclonal anti-HSP70-1 | 1:500 – 1:1000 | Recombinantly PfHsp70-x (C-terminal region) (Külzer *et al.*, 2012) | WEHI monoclonal facility |
| Rabbit polyclonal anti-HSP101 | 1:500 | Recombinant HSP101 (amino acids 68-170) (de Koning-Ward *et al.*, 2009) | Gifted from Tania de Koning-Ward, Deakin University |
| **Secondary Antibody** | **Dilution** | **Immunogen** | **Supplier** |
| Goat anti-mouse, Alexa Fluor™ 800 | 1:10,000 | Gamma Immunoglobins Heavy and Light chains | Thermo Scientific (Catalogue number A32730) |
| Goat anti-rabbit, Alexa Fluor™ 680 | 1:10,000 | Gamma Immunoglobins Heavy and Light chains | Thermo Scientific (Catalogue number A-21076) |

**Table S7. Antibodies used in western blotting**

| **Primary Antibody** | **Dilution** | **Immunogen** | **Supplier** |
|---|---|---|---|
| Mouse monoclonal anti-HA | 1:500 | Synthetic peptide from human influenza HA-tag, conjugated to Keyhole Limpet Haemocyanin | Sigma-Aldrich (Catalogue number H3663) |
| Rabbit polyclonal anti-EXP2 | 1:500 | Recombinant EXP2 (amino acids 25-287) (Bullen *et al.*, 2012) | WEHI monoclonal facility |
| Rabbit polyclonal bleed anti-RhopH3 | 1:1000 | Recombinant RhopH3 (Counihan *et al.*, 2017) | Gifted from Tania de Koning-Ward, Deakin University |
| Rabbit anti-MSP1-19 | 1:500 | Recombinant MSP1 (amino acids 1672-1766) (de Koning-Ward *et al.*, 2003) | WEHI monoclonal facility |
| Rabbit anti-RON4.2 | 1:500 | Recombinant RON4 (amino acid 437-661) (Richard *et al.*, 2010) | WEHI monoclonal facility |
| Rabbit anti-GAP45 | 1:500 | Recombinant GAP45 (Baum *et al.*, 2006) | WEHI monoclonal facility |
| Rat anti-HA 3F10 | 1:50 | Amino acids 98-106 from the human influenza virus hemagglutinin protein | Merck Life Science (Catalogue number 12158167001) |
| **Secondary Antibody** | **Dilution** | **Immunogen/Reactivity** | **Supplier** |
| Goat anti-mouse Alexa Fluor 594 | 1:2000 | Gamma Immunoglobins Heavy and Light chains | Thermo Scientific (Catalogue number A-11032) |
| Goat ant-rabbit Alexa Fluor™ 488 | 1:2000 | Gamma Immunoglobins Heavy and Light chains | Thermo Scientific (Catalogue number A-11001) |
| Goat anti-rat Alexa Fluor™ Plus 488 | 1:1000 | Gamma Immunoglobins Heavy and Light chains | Thermo Scientific (Catalogue number A48262) |
| Goat anti-mouse Alexa Fluor™ Plus 594 | 1:1000 | Gamma Immunoglobins Heavy and Light chains | Thermo Scientific (Catalogue number A-11032) |
| Goat anti-rabbit Alexa Fluor™ Plus 647 | 1:1000 | Gamma Immunoglobins Heavy and Light chains | Thermo Scientific (Catalogue number A32733) |
| SYTOX™ BLUE | 1:1000 | Gamma Immunoglobins Heavy and Light chains | Thermo Scientific (Catalogue number S11348) |

| | | | |
|---|---|---|---|
| Alexa Fluor™ 488 NHS Ester | 1:200 | Primary amines on proteins and ligands, amine-modified oligonucleotides | Thermo Scientific (Catalogue number A20000) |

**Table S8. Antibodies used in indirect immunofluorescence assays**

### Supplementary Methods

<u>Chemistry Methods</u>

NMR spectra were recorded on a Bruker Ascend™ 300. Chemical shifts are reported in ppm on the δ scale and referenced to the appropriate solvent peak. MeOD, DMSO-$d_6$, $D_2O$, and $CDCl_3$ contain $H_2O$. Chromatography was performed with silica gel 60 (particle size 0.040-0.063 μm) using an automated CombiFlash Rf purification system. LCMS were recorded on an Agilent LCMS system comprised of an Agilent G6120B Mass Detector, 1260 Infinity G1312B Binary pump, 1260 Infinity G1367E HiPALS autosampler and 1260 Infinity G4212B Diode Array Detector. Conditions for LCMS were as follows, column: Luna® Omega 3 μm PS C18 100 Å, LC Column 50 × 2.1 mm at 20 ºC, injection volume 2 μL, gradient: 5-100% B over 3 min (solvent A: $H_2O$ 0.1% formic acid; solvent B: ACN 0.1% formic acid), flow rate: 1.5 mL/min, detection: 254 nm, acquisition time: 4.3 min. Unless otherwise noted, all compounds were found to be >95% pure by this method.

*6-Amino-3,3-dimethyl-8-thioxo-1,4-dihydrothiopyrano[3,4-c]pyran-5-carbonitrile* (Int-1). *2,2*-Dimethyltetrahydropyran-4-one (3.0 g, 23.4 mmol) was dissolved in MeOH (5 mL) and carbon disulfide (2.11 mL, 35.1 mmol) was added. Malononitrile (1.86 g, 28.1 mmol) was added in three portions and triethylamine (1.14 mL, 8.2 mmol) was then added dropwise. The reaction was then stirred for 48 h at 20 ºC and concentrated *in vacuo*. The crude material was then purified by column chromatography eluting with a gradient of 100% DCM to 2% MeOH/DCM to afford Int-1 as a solid (4.2 g, 71%). $^1$H NMR (300 MHz, $CDCl_3$): δ 1.32 (s, 6H) 2.64 (s, 2H) 4.69 (s, 2H) 5.78 (br. s., 2H). LCMS, m/z 253.2 (100) $[M+H]^+$.

*3,3-Dimethyl-8-morpholino-6-sulfanyl-1,4-dihydropyrano[3,4-c]pyridine-5-carbonitrile* (Int-2). Int-1 (3.5 g, 13.9 mmol) was suspended in ethanol (15 mL) and morpholine (3.6 mL, 41.6 mmol) was added. The reaction mixture was heated to reflux under nitrogen for 72 h. The reaction mixture was cooled in an ice bath to 0 ºC and left at this temperature for 2 h. The precipitate that formed was filtrated and washed with cold ethanol to afford Int-2 as a solid (1.43, 33%). $^{1}$H NMR (300 MHz, $CDCl_3$): δ 1.31 (s, 6H) 2.63 (s, 2H) 3.17 - 3.44 (m, 4H) 3.64 - 3.85 (m, 4H) 5.75 (br. s., 2H). LCMS, m/z 306.2 (100) $[M+H]^+$.

*3-Bromo-12,12-dimethyl-8-morpholino-11-oxa-5-thia-4,7-diazatricyclo[7.4.0.02,6]trideca-1(9),2(6),3,7-tetraene* (MMV687794). Bromine (0.72 mL, 14.0 mmol) was added to a stirred solution of Int-2 (1.43 g, 4.68 mmol) in chloroform (10 mL) at 20 ºC and then heated to reflux for 3 h. The reaction was then cooled and dissolved in DCM (50 mL) and washed with saturated $NaHCO_3$ solution (30 mL), brine (30 ml), dried with $Na_2SO_4$ and concentrated. The crude material was then purified by column chromatography eluting with a gradient of 100% DCM to 30% EtOAc/DCM to afford MMV687794 (320 mg, 18%). $^{1}$H NMR (300 MHz, $CDCl_3$): δ 1.40 (s, 6H) 3.17 - 3.34 (m, 4H) 3.42 (s, 2H) 3.81 - 3.91 (m, 4H) 4.73 (s, 2H). LCMS, m/z 384.2 (100) $[M+H]^+$.

*3-Ethyl-12,12-dimethyl-8-morpholino-11-oxa-5-thia-4,7-diazatricyclo[7.4.0.02,6]trideca-1(9),2(6),3,7-tetraene* (Analog 1) and *12,12-dimethyl-8-morpholino-11-oxa-5-thia-4,7-diazatricyclo[7.4.0.02,6]trideca-1(9),2(6),3,7-tetraene* (Analog 2). MMV687794 (30 mg, 0.078 mmol) was added to a solution of dry THF (1 mL). $Pd(PPh_3)_2Cl_2$ (2.7 mg, 0.0039 mmol) was then added and the reaction purged with $N_2$. Diethylzinc (0.156 mL, 0.156 mmol) was then added dropwise and the reaction stirred at reflux for 16 h. The reaction mixture was then cooled to 20 ºC and the reaction filtered through diatomaceous earth. Prep LCMS afforded Analog 1 (5.2 mg, 20%) and Analog 2 (3.1 mg, 13%). Analog 1: $^{1}H$ NMR (300 MHz, $CDCl_3$): δ 1.36 - 1.50 (m, 9H) 3.06 - 3.20 (m, 4H) 3.20 - 3.30 (m, 4H) 3.79 - 3.92 (m, 4H) 4.77 (s, 2H). LCMS, m/z 334.2 (100) $[M+H]^+$. Analog 2: $^{1}H$ NMR (300 MHz, $CDCl_3$): δ 1.41 (s, 6H) 3.06 (s, 2H) 3.25 - 3.32 (m, 4H) 3.81 - 3.94 (m, 4H) 4.76 (s, 2H) 8.77 (s, 1H). LCMS, m/z 306.2 (100) $[M+H]^+$.

*In vitro* culturing of *P. falciparum*

Parasites were cultured using human RBCs (Australian Red Cross) at 2 or 4% HCT in TPP® tissue culture dishes (Thermo Scientific), Falcon® 6-well Clear Multiwell Plates (Thermo Scientific) or Nunc™ cell culture dishes (Thermo Scientific). Parasite cultures were kept in microisolator chambers (Labquip) gassed with 1% $O_2$, 5% $CO_2$, 94% $N_2$ (Supagas) and incubated at 37 °C. Complete RPMI (cRPMI) was used: RPMI 1640 (Merck Life Science), 25 mM HEPES (Gibco), 31.25 µg/mL gentamicin (Gibco), 0.2% $NaHCO_3$ (Thermo Scientific) and 0.5% AlbuMAX™ (Gibco). Parasites were monitored via Giemsa-stained thin blood smears as described (Trager and Jensen, 1976).

Sorbitol synchronisation (Lambros and Vanderberg, 1979) was performed to obtain synchronous ring-stage parasites. Mixed-stage parasite cell pellets (500 g/5-10 mins) were resuspended in 10-15× pellet volume of 5% D-Sorbitol (Merck Life Science), dispensed in a 10- or 50-mL conical tube (Thermo Scientific) and incubated in a water bath for ≥ 10 mins at 37 °C to lyse mature parasites (> 24 hpi). Sorbitol was aspirated from cell pellet (500 g/5-10 mins) and resuspended to 2 or 4% HCT in cRPMI.

Percoll purification (Rivadeneira *et al.*, 1983) was performed to obtain ≥80% purified schizonts. Schizont (≥44 hpi) parasite cell pellets (500 g/5-10 mins) were resuspended in 4-5× pellet volume of cRPMI, layered over 4-6 mL of 67% Percoll (Merck Life Science) in a 10 mL conical tube, and centrifuged (1500 g/15 mins) with reduced deacceleration. The dark middle layer containing schizonts were collected and washed in 10-20 mL cRPMI.

Magnet purification (Ribaut *et al.*, 2008) was performed to obtain ≥95% purified schizonts. MACS® magnetic separation columns (Miltenyi Biotec) were flushed with with 5 mL 80% ethanol, equilibrated 2× in 5 mL iRPMI (cRPMI without $NaHCO_3$ and AlbuMAX™) and loaded onto an in-house magnetic stand. A Dispoflex 3-way stopcock (Disposafe) and a flat-end protein gel-loading tip (Merck Life Science) was attached to the column to regulate flow rate. Schizont (≥44 hpi) parasite cell pellets (500 g/5-10 mins) were resuspended to ~20-24% HCT and dispensed into the columns for passage. Columns were washed 5× with 5 mL iRPMI once iRBCs completely pass through to remove RBCs. Bound cells were eluted with 5 mL iRPMI via a 10 mL syringe into a 50 mL conical tube and washed with 10-20 mL cRPMI (500 g/5-10 mins).

<u>Generation of p1.2-PF3D7_0403800-HA-*glmS* constructs for CRISPR-Cas9 transfection</u>

The p1.2-*Pf*SLIRP-HA-*glmS* plasmid for clustered regularly interspaced short palindromic repeats (CRISPR)-Cas9 transfection of *P. falciparum* contained three haemagglutinin (HA) epitope tags and a *glmS* riboswitch directly appended to *Pf*SLIRP to enable the study of its function within *P. falciparum* (Prommana *et al.*, 2013). As MMV794-resistant parasites 3C-C6 have a C36W mutation in *Pf*SLIRP, this mutation was also engineered into the plasmid to validate that MMV794 targets *Pf*SLIRP. The complete *Pf*SLIRP CRISPR-Cas9 construct, p1.2-*Pf*SLIRP-HA-*glmS*, was generated in a 14-step PCR involving the separate construction of the 3' and 5' homology flanks, removal of additional *BglII* restriction enzyme sites, insertion of the C36W mutation into the 5' homology flank, and ligation of the 3' and 5' homology flanks together.

The 3' homology flank was amplified with ABH_Eco5F and ABH_Kas6R from *P. falciparum* genomic DNA (gDNA) and ligated into a pJET1.2/blunt cloning vector. The sequence on the recombination flank was confirmed by Sanger sequencing and then ligated into the pUF1-Cas9G (Volz *et al.*, 2016) vector via *EcoRI* and *KasI* restriction sites.

For the construction of the 5' homology flank, as the C36W mutation is near the start of the *Pf*SLIRP (PF3D7_0403800) gene (Figure S6, C-W (TGT-TGG)), parts of the 5' untranslated region upstream of the start codon (Figure S6, grey box) had to be used as a 5' homology flank to stimulate recombination of the DNA construct into the gene's locus. The 5' homology flank was first amplified with ABH_Bgl1F and ABH_2R from *P. falciparum* gDNA (Figure S6). To restrict recombination to within the 5' homology flank, the region downstream of the 5' homology flank incorporating most of the ABH's coding sequence including the C36W mutation, was recodonised to that preferentially used by *Saccharomyces cerevisiae*. The recodonised region was synthesised as a series of overlapping DNA primers called gBlock gene fragments (Integrated DNA Technologies). Due to repetitive elements within the coding sequence the whole region could not be made as a single gBlock and had to be made as four smaller fragments (Figure S6). Fragment 4 was appended with the sequence for 3× HA tags so the protein could be detected by HA-specific antibodies. The four fragments and the 5' homology flank were joined using overlapping PCRs (Figure S6). Once the full 5' homology flank was complete, two internal *BglII* restriction sites had to be removed so that the only *BglII* site remaining was the cloning site at the 5' end of the 5' block. This was achieved by designing primers over the internal *BglII* sites that changed a single base to destroy the site without changing the amino acid (Figure S6). Following removal of internal *BglII* sites, a series of overlapping PCRs were performed to reconstruct the 5' homology flank and another series of PCRs were performed with the ABH_MutF and ABH_MutR primers to insert the C36W mutation into the 5' block (Figure S6).

After confirming the 5' homology flank was accurate by DNA sequencing, it was ligated into a pJET1.2/blunt cloning vector before ligation into the above pUF1-Cas9G vector containing the 3' recombination flank via *BglII* and *SpeI* restriction sites (Figure S6).

<u>Validation of CRISPR-Cas9 *Pf*SLIRP transgenic parasite lines</u>

To validate the correct integration of p1.2-*Pf*SLIRP(WT)-HA-*glmS* and p1.2-*Pf*SLIRP (C36W)-HA-*glmS* constructs into the genomic locus of transfected parasites, mixed-stage parasites at 4% HCT and ≥ 3% parasitaemia were harvested via saponin lysis. gDNA was extracted from the saponin pellet and correct integration of the 5' homology flank and C36W mutation was confirmed by DNA sequencing (Figure S7). Parental wild-type parasites (3D7) were also sequenced as a control. Previously generated MMV794-resistant parasites 3C-C6 (containing C36W mutation) and 3B-F10 (containing a deletion in the promoter region (Δ5'UTR) were sequenced to confirm the presence of the C36W mutation.

Saponin lysis

Parasite cultures were pelleted (500 g/5-10 mins) and resuspended in 10× pellet volumes of 0.1 or 0.15% saponin in ice-cold 1× PBS+Protease Inhibitor Cocktail Tablet (PI; Roche). Samples were incubated on ice for 10 mins and pelleted (3,200 g/10 mins) at 4 °C prior to washing 3× in 1× PBS+PI (Christophers and Fulton, 1939). Pellets were stored at -80 °C or used immediately.

gDNA extraction

gDNA was extracted from parasite pellets (from saponin lysis) following the manufacturer's protocol (DNeasy Blood & Tissue kit protocol (Qiagen)) with an additional centrifugation step. Specifically, following incubation with proteinase K, samples were pelleted (13,200 rpm/5 mins) to remove hemozoin crystals released upon parasite lysis. Pelleting was repeated (with transfer to a fresh microcentrifuge between each spin) until the supernatant was clear. Following this step, the protocol was resumed as per the manufacturer's instructions.

Western blotting (*Pf*SLIRP-HA validation, *Pf*SLIRP parasite stage expression, *Pf*SLIRP knockdown, carbonate extraction)

To validate the HA tagging of *Pf*SLIRP, mixed-stage (predominantly trophozoites and schizonts) 10 mL parasite cultures at 4% HCT and ≥ 5% parasitaemia were harvested via saponin lysis.

To examine stage-specific expression of *Pf*SLIRP, individual 30 mL ring (0-12 hpi), early trophozoite (18-24 hpi), late trophozoite (30-36 hpi), and schizonts (40-46 hpi) cultures were prepared. Contamination of asynchronous parasites was minimised by 2× sorbitol treatment of cultures. Post-sorbitol treatment, ring cultures were harvested immediately, early trophozoites and late trophozoites were harvested upon reaching the desired stage, schizonts were treated with 30 nM ML10 and Percoll purified before harvest via saponin lysis.

To examine the level of *Pf*SLIRP knockdown, early-trophozoite (18-24 hpi) C-WT and C-C36W parasites at 1% parasitaemia were treated with 2.5 mM glucosamine for 72 h until they reached the schizont stage, which were harvested via saponin lysis.

The following procedures were used for gel loading and western blotting for the above samples: Parasite pellets harvested from saponin lysis were resuspended in 10-20× pellet volume of 1× non-reducing sample buffer (NRSB), comprising 50 mM pH 6.8 Tris•HCl (Astral Scientific•Sigma-Aldrich), 10% glycerol (Astral Scientific), 2 mM EDTA (Merck Life Science), 2% SDS (Merck Life Science) and 0.01% bromophenol blue (Bio-Rad) in Mili-Q water. The samples were sonicated for 10 cycles (30 s on/off) at 4 °C in Bioruptor® Pico (Diagenode) before the addition of dithiothreitol (Merck Life Science) to a final concentration of 100 mM and denatured at 80 °C for 10 mins before SDS-PAGE and western blotting.

The following procedure was used for carbonate extraction (Grüring *et al.*, 2012). Saponin pellets harvested from 10 mL schizont-stage cultures at 4% HCT and ≥ 5% parasitaemia were resuspended in 20× pellet volume of PBS+PI and freeze-thawed for five cycles in a dry ice-ethanol slurry and a shaking heat block at 37 °C (PHMT Thermoshaker). Samples were pelleted (17,800 rpm/30 mins) at 4 °C and the supernatant was collected containing soluble proteins (PBS soluble fraction). The remaining pellet was resuspended in 20× initial pellet volume of 0.1 M $Na_2CO_3$ (pH 11), mixed on a tube roller at RT for 30 mins, and pelleted (17,800 g/30 mins) at 4 °C. The supernatant was collected containing peripheral membrane proteins (carbonate soluble fraction). The remaining pellet was washed 1× in 500 µL 1× PBS and centrifuged (17,800 rpm/5 mins), which contained integral membrane proteins (carbonate insoluble fraction). The PBS and carbonate soluble fractions were resuspended in $^{1}/_{3}$ volume of 4× NRSB to a final concentration of 1× NRSB, and the carbonate insoluble fraction was resuspended to the same volume in 1× NRSB. The samples were reduced with dithiothreitol and denatured with heat as above.

Resuspended samples were loaded into precast 4-12% Bis-Tris SDS-PAGE gels (Thermo Scientific) and electrophoresed in 1× MOPS buffer comprised of 50 mM MOPS (Merck Life Science), 50 mM Tris (Astral Scientific), 3.47 mM SDS (Merck Life Science), 1.03 mM EDTA (Merck Life Science) in distilled water at 200 V for 1 h (Mini gels) or 150 V for 1 h 45 mins (Midi gels). Precision Plus Protein™ All Blue Prestained Protein Standards (Bio-Rad) were used as protein markers. Gels were transferred to nitrocellulose membranes for western blotting using the iBlot 2 (20V, 7 mins; Merck Life Science) and blocked with 1% casein for 10-60 mins on orbital shaker at RT and probed with primary antibodies (Table S7) in 1 % casein for 1 h (RT) or overnight (4 °C). Unbound primary antibodies were removed by 3× 1-5 mins washes of 1× PBS before probing with fluorescent-conjugated secondary antibodies in 1 % casein (containing 0.01% azide) (Table S7). Unbound secondary antibodies were washed off as above and visualised using Odyssey® XF imaging system (LI-COR) and Image Studio™ v.1.0 programme (LI-COR).

<u>Indirect immunofluorescence assays (IFAs)</u>

IFAs were prepared either through iRBC smears or cultures. Images were captured using a Zeiss Axio Observer Z1 inverted widefield microscope equipped with a Plan-Apochromat 100×/1.40 Oil DIC objective, and the super-resolution radial fluctuations (SRRF) algorithm was used to capture super-resolution images. All images were processed using Fiji software.

IFAs using iRBC smears were prepared as follows: Thin smears of Percoll-purified iRBCs were made on glass slides and air dried for at least 1 h in the biosafety cabinet and stored at -20 °C until required for use. Glass slides were thawed at RT for 1 h prior to fixation with ice-cold 90% acetone/10% methanol for 2 mins. Hydrophobic boundaries were drawn around the smears with a Dako Pen (Agilent) before blocking with 3% BSA/0.02% TX100 for 1 h at RT within a humidity chamber. Cells were probed with primary antibodies (Table S7) overnight at 4 °C and unbound antibodies were removed by extensive washing in 0.05% polysorbate 20 (Astral Scientific) in 1× PBS before incubation with Alexa Fluor secondary antibodies (Thermo Scientific; Table S7) for 1 h in darkness at RT. Washes were completed as before and coverslips were mounted on slides with 2-5 µL of VECTASHIELD® Antifade Mounting Medium with DAPI (Vector Laboratories).

IFAs using iRBC cultures were prepared as follows: Percoll-purified iRBCs were settled onto poly-L-lysine- (Sigma-Aldrich) coated 10 mm round coverslips and fixed with 4% paraformaldehyde/0.0075% glutaraldehyde for 20 mins before quenching in 0.1 M glycine/0.1% TX100 in PBS. Cells were blocked in 3% BSA/0.02% TX100 in PBS for 1 hour at RT. To minimise non-specific binding, 1 mL primary antibodies in 3% BSA/0.02% TX100 were incubated with RBCs at 20% HCT at RT for 90 mins. The cross-adsorbed primary antibodies were then added to the cells overnight at 4 °C. Cells were washed with 0.02% TX100 in PBS to remove unbound antibodies and probed with Alexa Fluor secondary antibodies as above. Washes were performed as before. Coverslip mounting was performed as above.

Ultrastructure expansion microscopy (U-ExM) IFA

U-ExM (Gambarotto *et al.*, 2019; Bertiaux *et al.*, 2021; Liffner *et al.*, 2023) was performed as described with modifications. #1.5 13 mm round coverslips (Bio-Strategy) were rinsed in 80% ethanol (in mili-Q water), placed in a 24-well tissue culture plate (Merck Life Science) and washed 3× with filtered (0.22 μm) 500 μL 1X PBS. The dried coverslips were coated with 300 μL 0.1 mg/mL poly-D-lysine in 1× PBS (Gibco) for > 1 h at RT and rinsed 3X with filtered 500 μL 1× PBS, keeping the coverslips submerged during the final rinse. Percoll-purified schizont pellets of C-WT parasites were prepared as outlined in lattice light sheet imaging and resuspended to a final 4% HCT, and 50-100 μL of this resuspension is settled onto the coverslips for ≥ 30 mins at 37 °C. The supernatant is removed from the coverslips, and cells were fixed with methanol-free 4% formaldehyde/0.01% glutaraldehyde (Thermo Scientific/Merck Life Science) for 10-15 mins at RT. After the fixative was removed, cells were quenched/permeabilised with 500 μL 0.1 M glycine + 0.1% Triton X-100 in 1× PBS for ≥ 5 mins at RT, which were then removed and rinsed 2× with 500 μL of filtered 1× PBS. Cell-fixed coverslips were then incubated with 300 μL 1.4% formaldehyde/2% acrylamide (Merck Life Science) at 37 °C for 1-3 h or at 4 °C for > 3 h.

A gelation apparatus was assembled using a small shallow polystyrene foam box filled with ice and an aluminium heating/cooling block (48 × 1.5 mL tube; Merck Life Science) in the centre, levelled evenly. A piece of parafilm was placed in the centre of the aluminium cooling block. Before gelation, the coverslips were removed from the 24-well culture plate and dried by dabbing their edges against a Kimwipe and left cell-side up halfway on the edge of the parafilm for ≥ 15 mins. For gelation, 5 μL of 10% tetramethylethylendiamine (TEMED; Thermo Scientific) and 5 μL of 10% ammonium persulfate (APS; Thermo Scientific) were added to 90 μL of monomer solution (19% sodium acrylate (Merck Life Science), 10% acrylamide, 0.1% N,N'-methylenbisacrylamide (BIS; Merck Life Science)) (final concentration: 17.1% sodium acrylate, 1% acrylamide, 0.5% TEMED, 0.5% APS, 0.09% BIS, 1× PBS). The mixture was quickly mixed by pipetting and 35 μL was dispensed onto the parafilm and coverslips were placed cell-side down for gelation in the dark for ≥ 5 mins within the apparatus. Then, parafilm was placed in a humidity chamber wrapped in parafilm and aluminium foil and incubated at 37 °C for ≥ 30 mins. Post-incubation, the gels were gently removed from the coverslip with a small paintbrush and placed in a 2 mL microcentrifuge tube containing 1 mL denaturation buffer (200 mM SDS (Bio-Rad), 200 mM NaCl, 50 mM pH 9 Tris), which was topped up to 2 mL. The 2 mL microcentrifuge tubes were incubated at 95 °C for 60-90 mins, and the denaturation buffer was aspirated. The gels were gently transferred into a 90 mm petri dish (Thermo Scientific) with a paintbrush and washed 2× for ≥ 10 mins with Mili-Q water on a platform rocker at low speed for expanding. Mili-Q water was removed from the dish and a ruler was placed underneath the petri dish to measure the diameter of the gel for calculation of expansion factor (≥ 4×).

For staining, the gels were washed 2× for ≥ 10 mins with 1× PBS on a platform rocker at low speed for shrinking, cut in half with a spatula, and a half is placed in a Falcon® 6-well Clear Multiwell Plate for blocking with 1 mL of filtered (0.22 μm) 2% BSA (Merck Life Science) at RT for ≥ 20 mins. The gels were then probed with 1 mL of primary antibodies in 2% BSA at 4 °C overnight in the dark on a platform rocker, washed 3× for ≥ 5 mins with 1× PBS at RT on a platform rocker, then probed with 1 mL of secondary antibodies in 1× PBS at RT for 2-3 h in the dark on a platform rocker. The gels were washed 2× for ≥ 5 mins with 1× PBS at RT in the dark on a platform rocker. Then, the gels were transferred into a 90 mm petri dish and washed 2× for ≥ 10 mins with Mili-Q water at RT in the dark on a platform rocker for expansion.

For imaging, #1.5 25 mm round glass coverslips (Fisher Biotec) were placed in a staining jar (3D-printed in-house), washed 1× with 80% ethanol, coated with 0.1 mg/mL poly-D-lysine for ≥ 1 h at RT, and washed 2× with filtered 1× PBS For mounting, the water was aspirated from the gel and a rectangular section from the centre of the gel was cut. The rectangular section was cut again in half with one side flipped, and both halves of the rectangular section were placed onto a poly-D-lysine-coated 25 mm round coverslip on an Attofluor™ cell chamber (Thermo Scientific), with one half flipped, and the chamber was mounted onto a Piezo stage and imaged on the Zeiss laser scanning microscope 980 with Airyscan 2 using the Plan-Apochromat 63×/1.40 Oil DIC objective. The eyepieces (DAPI channel) were used to examine both halves of the gel to determine the gel orientation that contains the cells (parasite nuclei) to determine the correct gel orientation. Then, the following settings were used to acquire z-stacks with Airyscan 2: full Z-stack per Track, optimal sampling (0.148 μm), bidirectional frame scanning, 35 × 35 × 150 nm pixel size, 2 sampling rate, 0.70 μs pixel dwell time, 850 V detector gain, and 0.5% (405 nm), 1% (488 nm), 1% (561 nm) and 0.3% (639 nm) laser power. Images were processed using Imaris 10.2.

Baseline parasite metabolomic analysis of C-WT

C-WT and 3D7 parasite cultures were prepared as follows: 30 mL ring-stage parasite cultures at 4% HCT and 1% parasitaemia were sorbitol treated, and 3D7 cultures were treated ± 2.5 mM GlcN for 72 h until parasites reached the schizont stage. Magnet purification were performed to isolate schizonts, which were resuspended in cRPMI with 10 µM for 2-4 h to obtain synchronised late-stage schizonts (44-48 hpi). The parasite cultures were pelleted (500 g/10 mins) and quenched in 1 mL ice-cold 1× PBS, pelleted (1,000 g/5 mins) and the supernatant was aspirated.

Parasite lipids were extracted as above (Parasite lipidomics analysis (post-MMV794 treatment)) using single-phase chloroform/methanol/water extraction.

The LCMS data was processed as follows: Metabolomic samples were analysed by LC-MS using a Dionex UltiMate 3000 RS HPLC system fitted with Ascentis Express C8 column (2.7 µm, 2.1 × 100 mm, Supelco, Merck) and coupled with Q Extractive Orbitrap mass spectrometer (Thermo Scientific) as previously described (Aurelio *et al.*, 2016) with modifications. Mass spectrometry instrument used heated electrospray source operating in both positive and negative ion mode. The pooled biological quality control and blank samples were analysed periodically throughout each batch, and all batches analysed sequentially to avoid any impact of systematic instrument drift on metabolite signals. Metabolomics data were analysed using IDEOM (Creek *et al.*, 2012). A total number of 743 metabolites were putatively identified in *Pf*SLIRP experiment. Univariate statistical analysis used Welch's t-test (α = 0.05) in IDEOM, and multivariate statistical analysis used the web-based analytical tool MetaboAnalyst (Xia *et al.*, 2015).

Parasite lipidomics analysis (post-*Pf*SLIRP knockdown)

C-WT and C-C36W parasite cultures were prepared as follows: 30 mL ring-stage parasite cultures at 2% HCT and 2-3% parasitaemia were sorbitol treated, grew to early trophozoites, split into 2× 10 mL cultures each, and treated ± 2.5 mM GlcN for 72 h until parasites reached the schizont stage. During the 72 h-GlcN treatment period, the cultures were also treated with 30 nM ML10 at the 56 h timepoint overnight for 16 h to stall parasites at the late-schizont stage (44-48 hpi) at the 72 h timepoint. Then, these cultures were monitored to ensure schizont fitness via microscopy examination of Giemsa-stained thin blood smears. The schizont cultures were pelleted (500 g/10 mins), washed 1× with 50 mL cRPMI, dispensed into new culture dishes with 50 mL cRPMI and incubated at normal culture conditions for 4 h for invasion. Parasite invasion was validated via monitoring of cultures via microscopy examination of Giemsa-stained thin blood smears. Then, the cultures were pelleted (500 g/10 mins), resuspended in 10 mL cRPMI and quenched in a dry ice-ethanol slurry until they reached 10 °C. Finally, the cultures were pelleted (500 g/10 mins), and the pellets were harvested via 0.15% saponin lysis.

Parasite lipids were extracted and prepared as follows: Parasite lipids were extracted from saponin pellets using chloroform:methanol:water 1:3:1 (v/v/v) (Charital *et al.*, 2024). Firstly, 10 µL tridecanoic acid (C13:0) and phosphatidylcholine (PC; C21:0) were added to the pellets as internal standards, followed by 10 µL SPLASH® LIPIDOMIX® Mass Spec Standard (Avanti). This is followed by the addition of 1 µL 1% butylated hydroxytoluene (Sigma-Aldrich) in ethanol, 50 µL UltraPure™ DNase/RNase-free distilled $H_2O$ (Gibco), 300 µL methanol and sonication for 10 s (on/off) at RT until pellets were resuspended. Then, 100 µL chloroform was added to the resuspension, which were sonicated again for 10 s (on/off) at RT until pellets were resuspended. The pellets were kept on ice for 15 mins before centrifugation (5,000 rpm/5 mins), and the supernatant was transferred to a clear N8 screw neck glass vial (Macherey-Nagel). Then, 10 µL 0.1 M HCl and 150 µL methanol were added to the pellets and further resuspended with sonication for 10 s (on/off) at RT to ensure complete lipid extraction. Then, 300 µL chloroform was added to the pellet, which was resuspended with sonication for 10 s (on/off) at RT and left on ice for 10 mins before centrifugation (10,000 rpm/10 mins). During centrifugation, 200 µL chloroform and 300 µL aqueous phase 0.2% KCl/0.5% acetic acid were added to the previous N8 vial. Then, the supernatant from the pellets was added to the N8 vial to pool the organic phase, and the vial was vortexed for 5 s and centrifuged (2,000 rpm/30 s). The bottom layer (organic phase; 800 µL) was transferred to an amber N8 screw neck glass vial (Macherey-Nagel), and 300 µL were aliquoted into each of 2× clear N9 screw neck glass vials (Macherey-Nagel) with 0.25 mL glass conical micro-inserts (Macherey-Nagel). The samples were dried down in a SpeedVac with the drying rate set to high and stored at -20 °C until subsequent gas or liquid chromatography-mass spectrometry analysis. Samples were analysed in random sequence along with blank samples.

Total parasite lipids were analysed by LCMS as follows: Dried-down lipids were reconstituted in 80 µL methanol and incubated at 30 °C for 5 mins with vigorous vortexing. Total lipids in methanol (1 µL) were injected into an Agilent 1290 infinity/Infinity II LCMS system equipped with ZORBAX Eclipse Plus C18, 2.1 × 100 mm, 1.8 µm columns (Agilent) with Infinity II inline filter, 0.3 µm (Agilent) maintained at 45 °C. Samples were separated by the charge in the gradient of solvent A (water:acetonitrile:isopropanol, 5:3:2, (v/v/v), 1 mM ammonium formate) and solvent B (isopropanol:acetonitrile:water, 90:9:1 (v/v/v), 1 mM ammonium formate). The gradient was as follows: starting with a flow rate of 0.4 mL/min at 15% B and increasing to 50% over 2.5 mins, then to 57% at 2.6 mins, to 70% over 9 mins, to 93% at 9.1 mins, to 96% over 11 mins, to 100% at 11.1 mins, and hold to 12 mins, then back to 15% at 12.2 mins to 16 min (total of 16 mins). Equilibration was as follows; solvent was decreased from 100% B to 10% B over 0.1 mins and held for an additional 0.9 mins. The flow rate was then switched to 0.6 mL/mins for 1 min before returning to 0.4 mL/mins over 0.1 mins. Solvent B was held at 10% B for a further 0.9 mins at 0.4 mL/mins for a total cycle time of 15 mins. MS (Agilent 6495c triple quadrupole) was operated in for targeted analysis with DMRM (dynamic multiple reaction monitoring) with 650 ms of cycle time with following setting: Agilent Jet stream ion source with positive and negative switching with gas temperature at 150 °C, drying gas ($N_2$) 17 L/mins, nebuliser gas 20 psi, sheath gas temperature at 200 °C with flow at 10 L/mins, capillary voltage 3500 V for positive and -3000 V for negative mode, nozzle voltage at 1000 V for positive and -1500 V for negative mode. Acquisition DMRM method used was referred to (Huynh *et al.*, 2019) with a modification for *P. falciparum* lipid species. The resultant LCMS data was subjected to targeted analysis using Mass Hunter Quantification software (Agilent). Each lipid species was quantified using a calibration curve of each representative lipid with known abundance. All solvents used for this LCMS analysis were purchased from Sigma-Aldrich.

<u>Parasite lipidomics analysis (post-MMV794 treatment)</u>

C-WT and C-C36W parasite cultures were prepared as follows: 30 mL ring-stage parasite cultures at 4% HCT and ≥ 10% parasitaemia were sorbitol treated and stalled at the early schizont stage (40-44 hpi) with overnight (16 h) treatment with 30 nM ML10 at the early trophozoite stage. Magnet purification was performed on the 30 mL cultures to isolate schizonts, which were split equally into 2× 10 mL cultures. The cultures were treated with ± 3 μM MMV794 (10× 72-h parasite growth $EC_{50}$), incubated at normal culture conditions for invasion for 5 h and pelleted (500 g/10 mins). 160 μL of the supernatant was aliquoted for the analysis of MMV794 abundance, while the rest was aspirated. The parasite pellet was quenched in 1 mL ice-cold 1× PBS, pelleted (1,000 g/5 mins) and the supernatant was aspirated.

Parasite lipids were extracted as follows: Parasite lipids were extracted from parasite pellets using chloroform:methanol:water 2:6:1 (v/v/v). 200 μL chloroform/methanol/water was added to the pellets, immediately vortexed and mixed on a tube roller mixer (Ratek BTR5-12V) for 1 h at 4 °C. Samples were centrifuged at (16,100 g/1 min) and 160 μL of supernatant (lipids) was removed. The supernatants were dried down for lipidomics in the Savant® DNA110 SpeedVac® concentrator (Thermo Scientific) with the drying rate set to high. Dried-down samples were wrapped in Parafilm and stored at -80 °C until LCMS analysis.

MMV794 and total parasite lipids were analyzed by LCMS as follows: Dried-down samples were reconstituted in 80 μL butanol:methanol:water (45:45:10), sonicated at RT for 30 mins and centrifugated (21,000 g/10 mins) to remove cell debris. A pooled biological quality control comprised of 10 μL aliquots of each sample combined was included to identify metabolites and to monitor downstream sample stability. Samples (10 μL) were injected into a Dionex UltiMate™ rapid separation liquid chromatography system (Thermo Scientific) equipped with Supelco® Ascentis Express® C8 100 × 2.1 mm, 2.7 μm reversed-phase column (Merck) with a Luna C8 2 × 2 mm, 5 µm guard column (Phenomenex) maintained at 40 °C. Samples were separated by the change in gradient of solvent A (water:isopropanol, 60:40. (v/v), 8 mM ammonium formate) and solvent B (isopropanol:water, 98:2, (v/v) 8 mM ammonium formate and 2 mM formic acid (FA)). The gradient was as follows: starting with a flow rate of 0.2 mL/min at 0% B and increasing to 100% B over 25 mins, followed by a 2-min wash at 100% B and a 3-min re-equilibration at 0% B for a total cycle time of 30 mins. MS was operated in full scan mode with positive and negative polarity switching at 70,000 resolution with detection range of 140-1,300 *m/z* to collect positive and negative ion mode data. Automatic gain control target was $10^6$ ions with a 50 ms maximum injection time. The following settings were used: electro-spray ionisation source (ESI; 4.0 kV for both positive and negative mode), sheath gas (50), auxiliary gas (20), sweep gas (2), capillary temperature (300 °C), probe heater temperature (120 °C).

The LCMS data was processed as follows: The LCMS data obtained was subjected to untargeted processing using open-source software identification and evaluation of metabolomics data from LCMS (IDEOM) (Creek *et al.*, 2012) with an updated metabolite database encompassing peptides of proteinergic amino acids up to five amino acids long. Raw LCMS files were first converted to mzXML format using ProteoWizard, and extensible computational MS (XCMS) algorithm was used to identify peaks and converted them to peakML files. The Mzmatch.R algorithm was used to align and annotate related metabolite peaks with specific criteria: minimum detectable intensity of 100,000, relative standard deviation (RSD) of < 0.5 (reproducibility), and peak shape (codadw) of > 0.8. Default IDEOM parameters were used to eliminated noise and artefact peaks. Proton changes were corrected in negative (loss) or positive (gain) modes and putative metabolites were identified by correct mass within 3 ppm mass error searching against the IDEOM metabolite database. Algorithms for principal component analysis and hierarchical clustering of metabolites were also used in MetaboAnalyst (Chong *et al.*, 2018). Metabolomics data were presented as relative abundance values and statistical analyses were performed using Welch's *t*-test where significant interactions were observed and determined at $p < 0.05$.

<u>Untargeted comparative proteome profiling</u>

Parasite pellets were resuspended in 1 mL of lysis buffer (containing 5% SDS (Sigma-Aldrich), 100 mM 2-[4-(2-hydroxyethyl)piperazin-1-yl]ethanesulfonic acid (HEPES, pH 8.1) and heated at 95 °C for 10 mins. The samples were subject to probe sonication for 10 s (three cycles), and heated again at 95 °C for 10 mins before centrifugation 13,000 g/5 mins to obtain the soluble proteome. Lysate protein concentration was measured using Pierce™ Dilution-Free™ Rapid Gold BCA Protein Assay Kit (Thermo Scientific) and diluted to a concentration of 2 $mg \cdot mL^{-1}$. Supernatant containing the soluble protein fraction was taken (100 µL) for reduction (10 mM tris(2-carboxyethyl)phosphine hydrochloride (Pierce™)) and alkylation (40 mM 2-chloroacetamide) for 15 mins at 55 °C. The samples were then subjected to ProtiFi S-trap™ mini clean-up steps using the standard protocol with trypsinisation (added at a ratio of 1:50 enzyme/protein w/w) and desalted using in-house generated C18 StageTips (ProTiFi; Rappsilber *et al.*, 2003). Samples were then dried, and resuspended in 12 µL of 2% acetonitrile (ACN) and 0.1% FA, spiked with indexed retention time (iRT) peptides for LC-MS/MS analysis.

Solvent proteome profiling

3D7 parasite lysates were prepared as follows: 30 mL ring-stage parasite cultures at 4% HCT and ≥ 10% parasitaemia were sorbitol treated and stalled at the early schizont stage (40-44 hpi) with overnight (16 h) treatment with 30 nM ML10 at the early trophozoite stage. The cultures were washed 1× with 30 mL cRPMI and pelleted (500 g/10 mins). Parasite cell lysate were prepared as described (Ji *et al.*, 2025): The pellets were lysed in 10× pellet volume of hypotonic lysis buffer (150 mM ammonium chloride, 10 mM potassium bicarbonate, 1 mM EDTA in Mili-Q water), incubated on ice for 10 mins and pelleted (3,200 g /10 mins). The pellets were transferred into 1.5 mL microcentrifuge tubes and washed (16,100 g/1 min) with ice-cold 1× PBS until the supernatant was clear. Pellets were resuspended in 1 mL of lysis buffer (containing 1× PBS), 1.5 mM $MgCl_2$, 150 mM KCl, 0.4% Nonidet P-40 (NP-40), 250 U/mL Benzonase, protease inhibitor [Pierce Protease Inhibitor Mini Tablets] and phosphatase inhibitor [Roche PhosSTOP™ Tablets]) with shaking at 4 °C for 20 mins before centrifugation at 20,000 g for 10 mins at 4 °C to obtain the soluble proteome. Lysate protein concentration was measured using Pierce™ Dilution-Free™ Rapid Gold BCA Protein Assay Kit (Thermo Scientific) and diluted to a concentration of 2 $mg \cdot mL^{-1}$. The supernatant was then divided into equal aliquots ($n$ = 2-3) for vehicle (DMSO) and compound treatment (3 μM MMV794) and was incubated at room temperature for 15 mins. Solvent-induced proteome profiling was done as previously described (Zhang *et al.*, 2020; Van Vranken *et al.*, 2021). An equal volume of the lysate from drug/vehicle treated conditions was added to tubes containing additional lysis buffer and a mixture of acetone/ethanol/acetic acid (50:50:0.1 v/v/v, AEA) so that the percentage of AEA was adjusted to 9, 11, 13, 15 or 19% ($n$ = 2-4). Upon the addition of AEA, the samples were then incubated at 37 °C with vigorous shaking for 20 mins and then centrifuged at 4 °C (21,000 g, 15 min). Precipitated proteins were removed and the supernatant containing the soluble protein fraction was taken (85 µL) for reduction (10 mM tris(2-carboxyethyl)phosphine hydrochloride (Pierce™)) and alkylation (40 mM 2-chloroacetamide) for 15 min at 55 °C. The samples were then subjected to ProtiFi S-trap™ mini clean-up steps, desalted using in-house generated C18 StageTips (ProTiFi; Rappsilber *et al.*, 2003), dried and resuspended for LC-MS/MS analysis as previously described.

LC-MS/MS analysis with Orbitrap Astral using LFQ-DIA

Samples were analysed on a Vanquish Neo HPLC coupled to an Orbitrap Astral mass spectrometer (Thermo Scientific) equipped with NanoSpray Flex source (Thermo Scientific), using a 50 cm µPAC™ HPLC column (2.5 µm, 5 × 28 $\mu m^2$, C18, 100-200 Å, Thermo Fisher Scientific). Mobile phase A and B were 0.1% FA in water (Thermo Scientific, Optima LC-MS grade) and 0.1% FA/80% ACN (Thermo Scientific, Optima LC-MS grade), respectively. The column was heated to 50 °C, with the flow rate initially set to fast load (dependent on the column pressure limit of 700 nL/min and 450 bar) to minimise delay time. At the start of the active gradient, the flow rate was turned to 350 nL/min. The initial conditions of 4% B were ramped up to 30% from 0 to 45 mins, followed by an increase to 45% over 15 mins. The column was then washed for 5 mins at 97.5% B with a flow rate of 700 nL/min at the end of the active gradient, followed by fast equilibration on the Vanquish Neo LC, with a maximum pressure limit of 450 bar. For data independent acquisition (DIA) experiments on the Orbitrap Astral MS, MS1 spectra were collected in the Orbitrap at a resolving power of 240,000 over an *m/z* of 380-980. The MS1 normalised AGC target was set to 500% with a maximum injection time of 5 ms. DIA MS2 scans were acquired using the Astral analyser, covering the 380-980 *m/z* range, with a normalised AGC target of 800%, a maximum injection time of 3 ms and an HCD collision energy of 25%, with a default charge state of +2. Window placement optimisation was enabled, with isolation widths of 2 Th and active gradient length of 60 mins.

Analysis of SPP LC-MS/MS data

The proteomics data was processed using Spectronaut Pulsar (v16.1, Biognosys, Switzerland) (version 19.2.240905.62635) referencing an in house *P. falciparum* protein spectral library as per default settings (Siddiqui *et al.*, 2022). The raw data file exported from Spectronaut was analysed in DIA analyst (https://analyst- suites.org/apps/dia-analyst/). These applications automatically processed the data from Spectronaut, identified peptides, and quantified protein abundances. Briefly, proteins lacking quantitative values (reverse sequences, potential contaminant sequences, proteins identified solely by site) were filtered from the dataset to ensure robustness of subsequent analyses. Following pre-processing, all intensity values were $log_2$-transformed and grouped according to solvent and drug conditions. Protein-wise linear models incorporating empirical Bayes statistics were applied to perform differential expression analyses. Utilising the Bioconductor package limma, significantly stabilised proteins in each pairwise comparison were identified using a threshold of $p \leq 0.05$, controlled by the Benjamini-Hochberg method (Monash Proteomics and Metabolomics Facility, 2024) Additionally, proteins were considered to show significant stabilisation only if they had a *p* value $< 0.05$ and a log2 fold change $\geq 1.2$ between control and experimental groups.

<u>Assessment of MMV687794 warhead reactivity using β-mercaptoethanol</u>

The intrinsic warhead reactivity with β-mercaptoethanol was assessed following an adaptation of a previously described procedure(Galbiati *et al.*, 2023). Briefly, compounds were prepared in 10 mM DMSO stock solutions. A solution of 1 mM of test compound in MeOH and phosphate-buffered saline (PBS) (2.5:1 *v:v*) with and without the addition of β-mercaptoethanol (275 equivalents) were incubated at 37 ºC for 1 h and analysed by LCMS. Solutions were analyzed on an Agilent LCMS system comprised of an Agilent G6120B Mass Detector, 1260 Infinity G1312B Binary pump, 1260 Infinity G1367E HiPALS autosampler and 1260 Infinity G4212B Diode Array Detector. Conditions for LCMS were as follows, column: Luna® Omega 3 µm PS C18 100 Å, LC Column 50 × 2.1 mm at 20 ºC, injection volume 2 µL, gradient: 5-100% B over 3 min (solvent A: $H_2O$ 0.1% formic acid; solvent B: ACN 0.1% formic acid), flow rate: 1.5 mL/min, detection: 254 nm, acquisition time: 4.3 min.

Altschul, S.F., Gish, W., Miller, W., Myers, E.W., and Lipman, D.J. (1990). Basic local alignment search tool. *J Mol Biol* 215**,** 403-410.

Attias, M., Teixeira, D.E., Benchimol, M., Vommaro, R.C., Crepaldi, P.H., and De Souza, W. (2020). The life-cycle of Toxoplasma gondii reviewed using animations. *Parasites & Vectors* 13**,** 588.

Aurelio, L., Scullino, C.V., Pitman, M.R., Sexton, A., Oliver, V., Davies, L., Rebello, R.J., Furic, L., Creek, D.J., Pitson, S.M., and Flynn, B.L. (2016). From Sphingosine Kinase to Dihydroceramide Desaturase: A Structure-Activity Relationship (SAR) Study of the Enzyme Inhibitory and Anticancer Activity of 4-((4-(4-Chlorophenyl)thiazol-2-yl)amino)phenol (SKI-II). *J Med Chem* 59**,** 965-984.

Baum, J., Richard, D., Healer, J., Rug, M., Krnajski, Z., Gilberger, T.W., Green, J.L., Holder, A.A., and Cowman, A.F. (2006). A conserved molecular motor drives cell invasion and gliding motility across malaria life cycle stages and other apicomplexan parasites. *J Biol Chem* 281**,** 5197-5208.

Bertiaux, E., Balestra, A.C., Bournonville, L., Louvel, V., Maco, B., Soldati-Favre, D., Brochet, M., Guichard, P., and Hamel, V. (2021). Expansion microscopy provides new insights into the cytoskeleton of malaria parasites including the conservation of a conoid. *PLoS biology* 19**,** e3001020-e3001020.

Bullen, H.E., Charnaud, S.C., Kalanon, M., Riglar, D.T., Dekiwadia, C., Kangwanrangsan, N., Torii, M., Tsuboi, T., Baum, J., Ralph, S.A., Cowman, A.F., De Koning-Ward, T.F., Crabb, B.S., and Gilson, P.R. (2012). Biosynthesis, Localization, and Macromolecular Arrangement of the Plasmodium falciparum Translocon of Exported Proteins (PTEX)*. *Journal of Biological Chemistry* 287**,** 7871-7884.

Charital, S., Shunmugam, S., Dass, S., Alazzi, A.M., Arnold, C.S., Katris, N.J., Duley, S., Quansah, N.A., Pierrel, F., Govin, J., Yamaryo-Botté, Y., and Botté, C.Y. (2024). The acyl-CoA synthetase TgACS1 allows neutral lipid metabolism and extracellular motility in Toxoplasma gondii through relocation via its peroxisomal targeting sequence (PTS) under low nutrient conditions. *mBio* 15**,** e0042724.

Chong, J., Soufan, O., Li, C., Caraus, I., Li, S., Bourque, G., Wishart, D.S., and Xia, J. (2018). MetaboAnalyst 4.0: towards more transparent and integrative metabolomics analysis. *Nucleic Acids Res* 46**,** W486-w494.

Christophers, S.R., and Fulton, J.D. (1939). Experiments with Isolated Malaria Parasites (Plasmodium Knowlesi) Free from Red Cells. *Annals of Tropical Medicine & Parasitology* 33**,** 161-170.

Counihan, N.A., Chisholm, S.A., Bullen, H.E., Srivastava, A., Sanders, P.R., Jonsdottir, T.K., Weiss, G.E., Ghosh, S., Crabb, B.S., Creek, D.J., Gilson, P.R., and De Koning-Ward, T.F. (2017). Plasmodium falciparum parasites deploy RhopH2 into the host erythrocyte to obtain nutrients, grow and replicate. *Elife* 6.

Creek, D.J., Jankevics, A., Burgess, K.E., Breitling, R., and Barrett, M.P. (2012). IDEOM: an Excel interface for analysis of LC-MS-based metabolomics data. *Bioinformatics* 28**,** 1048-1049.

De Koning-Ward, T.F., Gilson, P.R., Boddey, J.A., Rug, M., Smith, B.J., Papenfuss, A.T., Sanders, P.R., Lundie, R.J., Maier, A.G., Cowman, A.F., and Crabb, B.S. (2009). A newly discovered protein export machine in malaria parasites. *Nature* 459**,** 945-949.

De Koning-Ward, T.F., O'donnell, R.A., Drew, D.R., Thomson, R., Speed, T.P., and Crabb, B.S. (2003). A new rodent model to assess blood stage immunity to the Plasmodium falciparum antigen merozoite surface protein 119 reveals a protective role for invasion inhibitory antibodies. *J Exp Med* 198**,** 869-875.

Dubey, J.P., Lindsay, D.S., and Speer, C.A. (1998). Structures of Toxoplasma gondiiTachyzoites, Bradyzoites, and Sporozoites and Biology and Development of Tissue Cysts. *Clinical Microbiology Reviews* 11**,** 267-299.

Ejotre, I., Reeder, D.M., Matuschewski, K., and Schaer, J. (2021). Hepatocystis. *Trends in Parasitology* 37**,** 456-457.

Feufack-Donfack, L.B., Baldor, L., Roesch, C., Tat, B., Orban, A., Seng, D., Salvador, J., Khim, N., Carias, L., King, C.L., Russell, B., Nosten, F., Ong, A.S.M., Mao, H., Renia, L., Lo, E., Witkowski, B., and Popovici, J. (2024). The PvRBP2b-TfR1 interaction is not essential for reticulocytes invasion by Plasmodium vivax isolates from Cambodia. *npj Vaccines* 9**,** 232.

Fisher, C., Seferidis, N., Zilli, J., Roberts, T., and Harcourt-Brown, T. (2024). Insights into the clinical presentation, diagnostics and outcome in dogs presenting with neurological signs secondary to infection with Neospora caninum: 41 cases (2014-2023). *Journal of Small Animal Practice* 65**,** 582-588.

Florent, I., Chapuis, M.P., Labat, A., Boisard, J., Leménager, N., Michel, B., and Desportes-Livage, I. (2021). Integrative taxonomy confirms that Gregarina garnhami and G. acridiorum (Apicomplexa, Gregarinidae), parasites of Schistocerca gregaria and Locusta migratoria (Insecta, Orthoptera), are distinct species. *Parasite* 28**,** 12.

Frey, C.F., Regidor-Cerrillo, J., Marreros, N., García-Lunar, P., Gutiérrez-Expósito, D., Schares, G., Dubey, J.P., Gentile, A., Jacquiet, P., Shkap, V., Cortes, H., Ortega-Mora, L.M., and Álvarez-García, G. (2016). Besnoitia besnoiti lytic cycle in vitro and differences in invasion and intracellular proliferation among isolates. *Parasites & Vectors* 9**,** 115.

Galbiati, A., Bova, S., Pacchiana, R., Borsari, C., Persico, M., Zana, A., Bruno, S., Donadelli, M., Fattorusso, C., and Conti, P. (2023). Discovery of a spirocyclic 3-bromo-4,5-dihydroisoxazole covalent inhibitor of hGAPDH with antiproliferative activity against pancreatic cancer cells. *Eur J Med Chem* 254**,** 115286.

Gambarotto, D., Zwettler, F.U., Le Guennec, M., Schmidt-Cernohorska, M., Fortun, D., Borgers, S., Heine, J., Schloetel, J.G., Reuss, M., Unser, M., Boyden, E.S., Sauer, M., Hamel, V., and Guichard, P. (2019). Imaging cellular ultrastructures using expansion microscopy (U-ExM). *Nat Methods* 16**,** 71-74.

Grüring, C., Heiber, A., Kruse, F., Flemming, S., Franci, G., Colombo, S.F., Fasana, E., Schoeler, H., Borgese, N., Stunnenberg, H.G., Przyborski, J.M., Gilberger, T.W., and Spielmann, T. (2012). Uncovering common principles in protein export of malaria parasites. *Cell Host Microbe* 12**,** 717-729.

Guérin, A., and Striepen, B. (2020). The Biology of the Intestinal Intracellular Parasite Cryptosporidium. *Cell Host & Microbe* 28**,** 509-515.

Huynh, K., Barlow, C.K., Jayawardana, K.S., Weir, J.M., Mellett, N.A., Cinel, M., Magliano, D.J., Shaw, J.E., Drew, B.G., and Meikle, P.J. (2019). High-Throughput Plasma Lipidomics: Detailed Mapping of the Associations with Cardiometabolic Risk Factors. *Cell Chemical Biology* 26**,** 71-84.e74.

Ji, Y., Morrow, J.P., Macraild, C.A., Zhang, H., Giannangelo, C., Schittenhelm, R.B., Creek, D.J., and Siddiqui, G. (2025). Validation of solvent proteome profiling for antimalarial drug target deconvolution. *Int J Parasitol Drugs Drug Resist* 29**,** 100626.

Jumper, J., Evans, R., Pritzel, A., Green, T., Figurnov, M., Ronneberger, O., Tunyasuvunakool, K., Bates, R., Žídek, A., Potapenko, A., Bridgland, A., Meyer, C., Kohl, S.a.A., Ballard, A.J., Cowie, A., Romera-Paredes, B., Nikolov, S., Jain, R., Adler, J., Back, T., Petersen, S., Reiman, D., Clancy, E., Zielinski, M., Steinegger, M., Pacholska, M., Berghammer, T., Bodenstein, S., Silver, D., Vinyals, O., Senior, A.W., Kavukcuoglu, K., Kohli, P., and Hassabis, D. (2021). Highly accurate protein structure prediction with AlphaFold. *Nature* 596**,** 583-589.

Külzer, S., Charnaud, S., Dagan, T., Riedel, J., Mandal, P., Pesce, E.R., Blatch, G.L., Crabb, B.S., Gilson, P.R., and Przyborski, J.M. (2012). Plasmodium falciparum-encoded exported hsp70/hsp40 chaperone/co-chaperone complexes within the host erythrocyte. *Cellular Microbiology* 14**,** 1784-1795.

Lakew, B.T., Eastwood, S., and Walkden-Brown, S.W. (2023). Epidemiology and Transmission of Theileria orientalis in Australasia. *Pathogens* 12.

Lambros, C., and Vanderberg, J.P. (1979). Synchronization of Plasmodium falciparum erythrocytic stages in culture. *J Parasitol* 65**,** 418-420.

Liffner, B., Cepeda Diaz, A.K., Blauwkamp, J., Anaguano, D., Frolich, S., Muralidharan, V., Wilson, D.W., Dvorin, J.D., and Absalon, S. (2023). Atlas of Plasmodium falciparum intraerythrocytic development using expansion microscopy. *Elife* 12.

Monash Proteomics and Metabolomics Facility (2024). "Manual for DIA-Analyst".).

Prommana, P., Uthaipibull, C., Wongsombat, C., Kamchonwongpaisan, S., Yuthavong, Y., Knuepfer, E., Holder, A.A., and Shaw, P.J. (2013). Inducible knockdown of Plasmodium gene expression using the glmS ribozyme. *PLoS One* 8**,** e73783.

Protifi S-TrapTM mini spin column digestion protocol.

Rappsilber, J., Ishihama, Y., and Mann, M. (2003). Stop and Go Extraction Tips for Matrix-Assisted Laser Desorption/Ionization, Nanoelectrospray, and LC/MS Sample Pretreatment in Proteomics. *Analytical Chemistry* 75**,** 663-670.

Ribaut, C., Berry, A., Chevalley, S., Reybier, K., Morlais, I., Parzy, D., Nepveu, F., Benoit-Vical, F., and Valentin, A. (2008). Concentration and purification by magnetic separation of the erythrocytic stages of all human Plasmodium species. *Malaria Journal* 7**,** 45.

Richard, D., Macraild, C.A., Riglar, D.T., Chan, J.A., Foley, M., Baum, J., Ralph, S.A., Norton, R.S., and Cowman, A.F. (2010). Interaction between Plasmodium falciparum apical membrane antigen 1 and the rhoptry neck protein complex defines a key step in the erythrocyte invasion process of malaria parasites. *J Biol Chem* 285**,** 14815-14822.

Rivadeneira, E.M., Wasserman, M., and Espinal, C.T. (1983). Separation and concentration of schizonts of Plasmodium falciparum by Percoll gradients. *J Protozool* 30**,** 367-370.

Shrestha, A., Abd-Elfattah, A., Freudenschuss, B., Hinney, B., Palmieri, N., Ruttkowski, B., and Joachim, A. (2015). Cystoisospora suis - A Model of Mammalian Cystoisosporosis. *Front Vet Sci* 2**,** 68.

Siddiqui, G., De paoli, A., Macraild, C.A., Sexton, A.E., Boulet, C., Shah, A.D., Batty, M.B., Schittenhelm, R.B., Carvalho, T.G., and Creek, D.J. (2022). A new mass spectral library for high-coverage and reproducible analysis of the Plasmodium falciparum–infected red blood cell proteome. *GigaScience* 11.

Sievers, F., Wilm, A., Dineen, D., Gibson, T.J., Karplus, K., Li, W., Lopez, R., Mcwilliam, H., Remmert, M., Söding, J., Thompson, J.D., and Higgins, D.G. (2011). Fast, scalable generation of high-quality protein multiple sequence alignments using Clustal Omega. *Mol Syst Biol* 7**,** 539.

Söderström, M., Malkamäki, S., Sukura, A., Sainmaa, S., and Airas, N. (2021). Sarcocystis calchasi in a captive Patagonian conure (Cyanoliseus patagonus) in Finland. *Journal of Comparative Pathology* 189**,** 135-140.

Trager, W., and Jensen, J.B. (1976). Human malaria parasites in continuous culture. *J Parasitol* 91**,** 484-486.

Uilenberg, G. (2006). Babesia—A historical overview. *Veterinary Parasitology* 138**,** 3-10.

Valigurová, A. (2012). Sophisticated adaptations of Gregarina cuneata (Apicomplexa) feeding stages for epicellular parasitism. *PLoS One* 7**,** e42606.

Van Vranken, J.G., Li, J., Mitchell, D.C., Navarrete-Perea, J., and Gygi, S.P. (2021). Assessing target engagement using proteome-wide solvent shift assays. *eLife* 10**,** e70784.

Varadi, M., Bertoni, D., Magana, P., Paramval, U., Pidruchna, I., Radhakrishnan, M., Tsenkov, M., Nair, S., Mirdita, M., Yeo, J., Kovalevskiy, O., Tunyasuvunakool, K., Laydon, A., Žídek, A., Tomlinson, H., Hariharan, D., Abrahamson, J., Green, T., Jumper, J., Birney, E., Steinegger, M., Hassabis, D., and Velankar, S. (2024). AlphaFold Protein Structure Database in 2024: providing structure coverage for over 214 million protein sequences. *Nucleic Acids Res* 52**,** D368-d375.

Volz, J.C., Yap, A., Sisquella, X., Thompson, J.K., Lim, N.T., Whitehead, L.W., Chen, L., Lampe, M., Tham, W.H., Wilson, D., Nebl, T., Marapana, D., Triglia, T., Wong, W., Rogers, K.L., and Cowman, A.F. (2016). Essential Role of the PfRh5/PfRipr/CyRPA Complex during Plasmodium falciparum Invasion of Erythrocytes. *Cell Host Microbe* 20**,** 60-71.

Xia, J., Sinelnikov, I.V., Han, B., and Wishart, D.S. (2015). MetaboAnalyst 3.0--making metabolomics more meaningful. *Nucleic Acids Res* 43**,** W251-257.

Zhang, X., Wang, Q., Li, Y., Ruan, C., Wang, S., Hu, L., and Ye, M. (2020). Solvent-Induced Protein Precipitation for Drug Target Discovery on the Proteomic Scale. *Anal. Chem.* 92**,** 1363-1371.
